# Charge-Driven Condensation of RNA and Proteins Suggests Broad Role of Phase Separation in Cytoplasmic Environments

**DOI:** 10.1101/2020.04.23.057901

**Authors:** Bercem Dutagaci, Grzegorz Nawrocki, Joyce Goodluck, Ali Akbar Ashkarran, Charles G. Hoogstraten, Lisa J. Lapidus, Michael Feig

## Abstract

Phase separation processes are increasingly being recognized as important organizing mechanisms of biological macromolecules in cellular environments. Well established drivers of liquid-liquid phase separation are multi-valency and intrinsic disorder. Here, we show that globular macromolecules may condense simply based on electrostatic complementarity. More specifically, phase separation of mixtures between RNA and positively charged proteins is described from a combination of multiscale computer simulations with microscopy and spectroscopy experiments. Condensates retain liquid character and phase diagrams are mapped out as a function of molecular concentrations in experiment and as a function of molecular size and temperature via simulations. The results suggest a more general principle for phase separation that is based primarily on electrostatic complementarity without invoking polymer properties as in most previous studies. Simulation results furthermore suggest that such phase separation may occur widely in heterogenous cellular environment between nucleic acid and protein components.

**STATEMENT OF SIGNIFICANCE:** Liquid-liquid phase separation has been recognized as a key mechanism for forming membrane-less organelles in cells. Commonly discussed mechanisms invoke a role of disordered peptides and specific multi-valent interactions. We report here phase separation of RNA and proteins based on a more universal principle of charge complementarity that does not require disorder or specific interactions. The findings are supported by coarse-grained simulations, theory, and experimental validation via microscopy and spectroscopy. The broad implication of this work is that condensate formation may be a universal phenomenon in biological systems.

## INTRODUCTION

Biological cells compartmentalize to support specific functions such as stress response (1, 2), regulation of gene expression (3, 4) and signal transduction (5). Compartmentalization by organelles that are surrounded by lipid membranes is well known. In addition, membrane-less organelles that result from coacervation have been described (6–9). In the nucleus they include the nucleolus (10, 11), nuclear speckles (12, 13), and cajal bodies (14–16); stress granules (2, 17, 18), germ granules (19, 20), and processing bodies (21, 22) have been found in the cytoplasm. The formation of coacervates via condensation and phase separation depends on the composition and concentration of the involved macromolecules (9) as well as environmental conditions such as pH, temperature, and the concentration of ions (6, 23, 24). Multivalent interactions, the presence of conformationally flexible molecules (25–27), and electrostatic interactions between highly charged molecules (25, 27–31) are well-known as the key factors that promote phase separation (PS), in particular via complex coacervation (32, 33). In biological environments, nucleic acids such as RNA have been found to play a prominent role in condensate formation due to their charge (34–40). Another component often found in biological condensates are intrinsically disordered peptides (IDPs) that may phase separate alone or in combination with RNA (24, 38, 41–43), although disorder may not be essential for phase separation (44, 45). Condensates often materialize as droplets, where experiments such as fluorescence recovery after photobleaching (FRAP) (46, 47) or direct visualization of merging droplets (39, 48) may confirm liquid-like behavior. However, a variety of other types of less-liquid condensates involving biomolecules have been described including clusters, gels, and aggregation to fibrils or tangles (18, 49–53). In those cases, internal diffusional dynamics may be highly retarded or lost. The high degree of polydispersity in biological multicomponent systems presents additional changes. An especially intriguing aspect of polydisperse systems is the propensity for multiphasic behavior (10, 44, 54) which imparts a potential for fine-grained tunable spatial patterning of biomolecules in cellular systems (44).

Biomolecular condensates have been studied extensively (55). Microscopy (23, 56, 57), nuclear magnetic resonance (NMR) spectroscopy (17, 41), fluorescence spectroscopy (10, 58, 59), X-ray diffraction (60, 61), and scattering methods (58, 62, 63) have characterized *in vitro* (10, 17, 23, 41, 57, 59–61, 63) and *in vivo* systems (19, 64, 65). Theoretical studies have complemented experiments (55, 66), including particle-based simulations (67) and analytical approaches based on polymer (43, 68) and colloid theories (69–71). Additional insights into specific interactions have come from molecular dynamics (MD) simulation studies (59, 72, 73). Polymer aspects of IDPs and unstructured RNA were emphasized in applications of Flory-Huggins theory in combination with simulations (10, 67, 74, 75). Related studies in the colloid field have described the phase behavior of macromolecules and nanoparticles as single spherical particles (69–71). However, most of the latter studies so far have focused on liquid-solid transitions and the formation of finite size clusters in monodisperse systems. Despite progress, it has remained unclear what components can lead to condensation, especially in highly heterogeneous cellular environments.

As most previous studies have focused on specific biomolecules undergoing PS, we focus here on the question of how general of a phenomenon PS may be in biological environments and what factors may determine the propensity for PS in a heterogeneous system. The starting point is a molecular model of a bacterial cytoplasm that was established by us previously (76, 77) and that was simulated here again but using colloid-like spherical particles with a potential parameterized against atomistic MD simulations of concentrated protein solutions. We found that distinct phases enriched with highly negatively charged RNA and positively charged proteins were formed in the simulations, consistent with a generic electrostatic mechanism that does not require specific interaction sites or elements of disorder and may apply broadly to mixtures of nucleic acids and proteins. The phase behavior seen in the cytoplasmic system was reproduced in reduced five- and two-component models and described by an analytical model where we could systematically vary molecular charge, size, and concentrations. The main prediction of the formation of condensates between RNA and positively charged proteins was confirmed experimentally via confocal microscopy and FRET spectroscopy and the nature of the condensates was analyzed further via dynamic light scattering and nuclear magnetic resonance spectroscopy. The details of the findings from simulation, theory, and experiment are described in the following.

## RESULTS

### Liquid condensates enriched in tRNA and ribosomes form in a model bacterial cytoplasm

A model of the cytoplasm of *Mycoplasma genitalium* established previously (76, 77) was simulated at a coarse-grained (CG) level with one sphere per macromolecule or complex (SI Methods and SI Sheet 1). CG particle interactions were calibrated against results from atomistic MD simulations of concentrated protein solutions. The parameters involve only two particle-dependent properties, namely size and charge (SI Methods). Droplet-like condensates formed spontaneously within 20 μs (Fig. 1A/B) and remained present during 1 ms simulation time. Similar results were obtained with an alternate effective charge model that resulted in better agreement between theory and experiment (see below; Fig. S1) Two types of condensates were observed: one type contained predominantly tRNA and positively charged proteins; the other type contained ribosomes (RP) and positively charged proteins. The RP condensates also attracted the weakly negatively charged GroEL particles at the surface (Fig. 1A). The condensates increased in size as the system size was increased from 100 to 300 nm (Fig. 1A). This observation is consistent with PS rather than finite-size cluster formation. The presence of multiple droplets in the 300 nm system suggests incomplete convergence, but as the droplets grow in size, further merging becomes kinetically limited due to slowing diffusion. We did not find evidence for growth via Ostwald ripening where particles preferentially evaporate from smaller condensates and redeposit onto larger condensates. Further analysis focused on the condensates observed in the 100-nm system.

**Figure 1.**
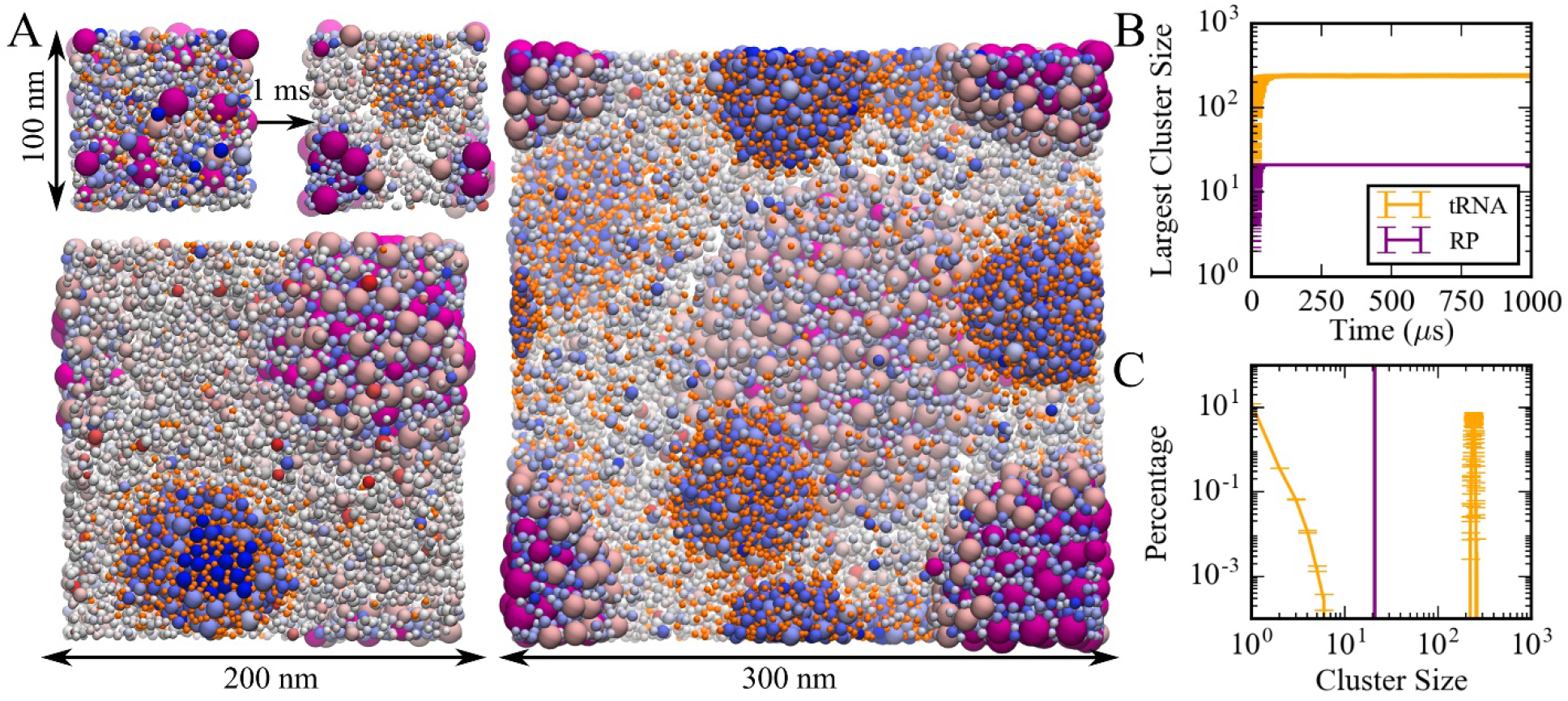
(A) Coarse-grained simulations of a model bacterial cytoplasm. Initial and final frames for 100 nm box and final frames for 200 and 300 nm boxes are shown with tRNAs in orange, ribosomes in magenta, and other molecules colored according to their charges (blue towards positive charges; red towards negative charges). Sphere sizes are shown proportional to molecular sizes. Large pink spheres correspond to GroEL particles. (B) Size of the largest cluster vs. simulation time in 100 nm system. (C) Cluster size distributions for tRNA and RP during the last 500 μs in the 100 nm system.

Cluster analysis (SI Methods) considered interactions between the nucleic acids and positively charged proteins to obtain trajectory-averaged cluster size distributions (Fig. 1C). Most tRNA (87%) was part of a condensate. The remaining fraction of tRNA existed as monomers or small clusters, suggesting coexistence of dilute and condensed phases. RP particles were only found in the RP condensates and therefore underwent a complete phase transition. Total macromolecular volume fractions inside tRNA and RP condensates were 0.42 and 0.58, respectively, whereas volume fractions for just tRNA and RP inside their respective condensates were 0.07 and 0.26. The condensates had significantly higher macromolecular densities than the rest of the simulated system (Fig. S2). The moderately high volume fractions for tRNA condensates are still within the range of concentrated liquid phases (78), but the higher volume fractions in the RP condensate tend towards solid- or gel-like phases (78). Radial distribution functions of tRNA and RP from the center of the corresponding condensates show a relatively smooth decay with a soft boundary for tRNA condensates (Fig. S3), also consistent with a liquid-like phase, whereas distinct peaks and a sharper boundary for RP indicate a highly ordered arrangement in the RP condensates.

Both tRNA and RP condensates contained (positively charged) proteins at high concentrations. tRNA and RP interactions with those proteins were favorable as evidenced by a strong peak in the pairwise radial distribution function *g(r)* at contact distance (Fig. S4). The charge and size of the proteins attracted to the condensates differed between tRNA and RP condensates (Fig. S5). In the tRNA condensates, large proteins with radii of 3 nm and above and with charges of 10 and above were preferred. In contrast, the proteins in the RP condensates were smaller, with radii of 3 nm or less, and many proteins had charges below 10.

The dynamics inside and outside the condensates was analyzed further in terms of translational diffusion coefficients (*D_tr_*) calculated based on mean-squared displacements (Fig. S6). Diffusion during the last 1 μs of the simulation was compared with diffusion during the first 1 μs when condensates were not yet formed. Molecule-specific values of *D_tr_* are given in SI Sheet 1. As a function of the radius of the macromolecules (Fig. S7), *D_tr_* values follow a similar trend as observed before in atomistic simulations of the same system. Diffusion outside the condensates resembled diffusion in the dispersed phase. In tRNA condensates, the diffusion of macromolecules is similar to the dispersed phase or is moderately retarded, depending on the molecule, and consistent with reduced diffusion in increased protein concentrations seen in experiment (79, 80). In RP condensates, diffusion is reduced to a greater extent, but liquid-like dynamics is still maintained for most types of macromolecules as they diffuse around a relatively static RP cluster (SI Movie 1).

### Factors promoting RNA condensation in a reduced five-component model system

A simplified five-component system was constructed to reproduce the RNA condensation observed in the cytoplasmic model. The simplified model consisted of tRNA, ribosomes (RP), large (POS_L_, *q* = 20, *r* = 3.5 nm) and small (POS_S_, *q* = 1, *r* = 2.52 nm) positively charged proteins as well as neutral crowders (CRW, *q* = 0, *r* = 2.52 nm). tRNA and RP concentrations were initially set as in the cytoplasmic model while concentrations, sizes, and charges of the other three particle types were adjusted to match the total number of particles, total molecular volume, and total charge of the cytoplasmic system as closely as possible. Subsequently, a series of simulations were run at different concentrations and with different parameters (SI Sheet 2).

In simulations of the five-component model, tRNA and RP condensed separately as in the cytoplasmic model (Fig. S8). Again, the condensates formed quickly, within 50 μs (Fig. S8), and cluster size distributions of tRNA and RP resembled the results from the cytoplasmic system (*cf*. Figs. 1 and S8). However, in contrast to the cytoplasmic system, we found a small fraction (2% on average) of RP particles in the dilute phase. As in the cytoplasmic model, tRNA strongly preferred interactions with the larger POS_L_ particles, whereas RP interacted favorably with both POS_S_ and POS_L_ (Fig. S9). tRNA condensates remained fully liquid as in the cytoplasmic system. From the last 100 μs of the simulation, we obtained diffusion coefficients *D_tr_* for tRNA of 28.3 ± 0.7 and 59.0 ± 0.5 nm^2^/μs inside and outside of the condensates, respectively, similar to values of 16.3 ± 0.1 and 55.5 ± 0.8 nm^2^/μs in the cytoplasmic system. Diffusion coefficients for RP inside and outside of the RP condensates were 0.49 ± 0.01 and 0.80 ± 0.4 nm^2^/μs, respectively, compared to *D_tr_* = 0.34 ± 0.01 nm^2^/μs for RP in the cytoplasmic condensates.

RP and POS_L_ concentrations were varied systematically, while the concentration of POS_S_ was kept constant and the number of CRW particles was adjusted to maintain a constant total molecular volume (SI Sheet 2). Cluster size distributions were extracted (Fig. S10) and the fraction of tRNA and RP in the large clusters was determined (Fig. 2). Some degree of clustering occurs at all concentrations, but condensation requires that a significant fraction of particles is found in the largest clusters. Based on a criterion that at least half of the particles are found in one or few large clusters, tRNA and RP condensation occurs for [POS_L_] > 100 μM (Fig. 2).

**Figure 2.**
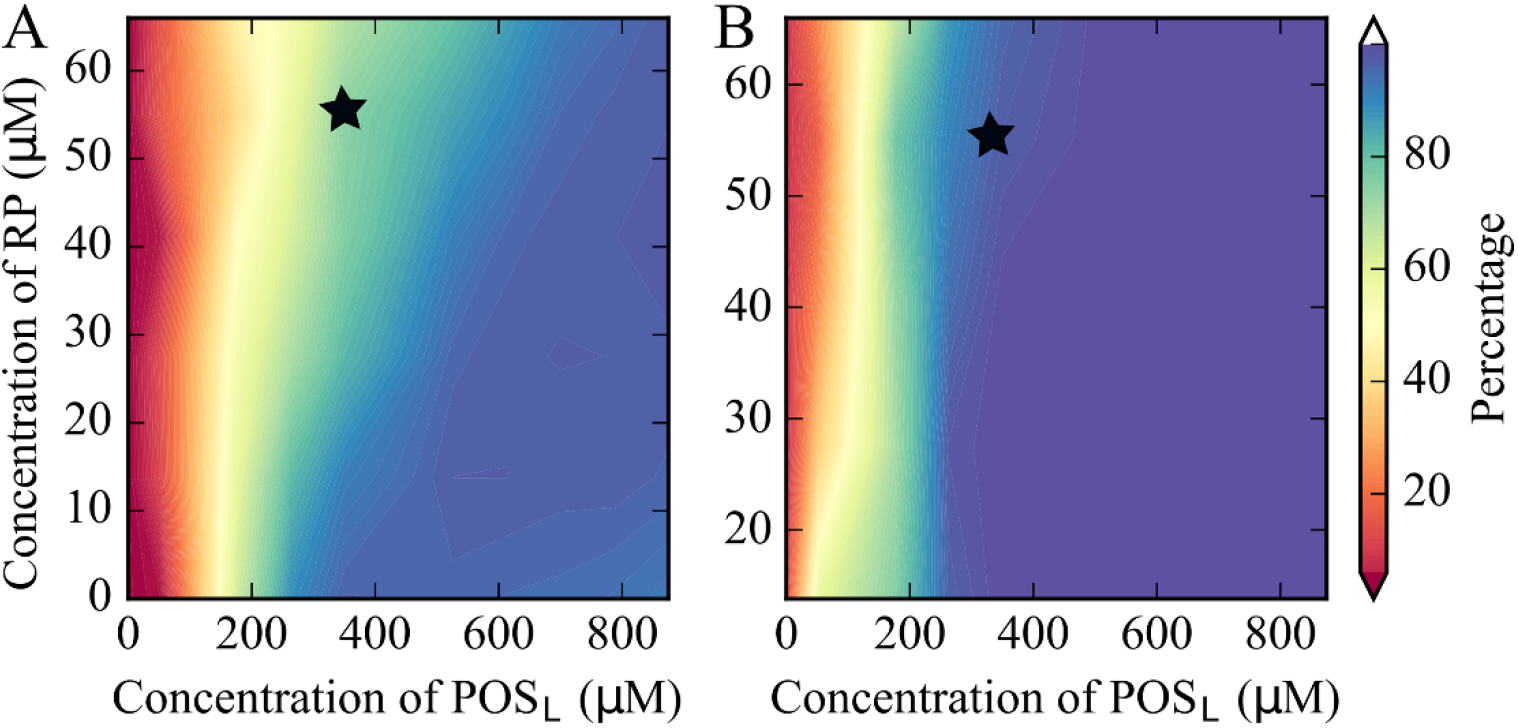
Percentage of tRNA (A) and RP (B) particles in largest clusters in coarse-grained simulations of the five-component model system as a function of [RP] and [POS_L_]. The black star indicates the conditions that match the cytoplasmic model.

Increasing [RP] reduces the amount of tRNA in the tRNA condensates and effectively raises the critical POS_L_ concentration above which tRNA forms condensates (Fig. 2). This can be understood from competition for POS_L_. tRNA only interacts significantly with POS_L_ (Fig. S11) and needs POS_L_ to form condensates, whereas RP interacts with both POS_S_ and POS_L_ (Fig. S12) and therefore draws POS_L_ from tRNA condensates (Fig. S13). For [POS_L_] > 500 μM, the fraction of tRNA particles in the tRNA condensates is relatively constant (Fig. 2). However, the number of POS_L_ particles in the condensates increases as the total [POS_L_] increases (Fig. S13A). This results in larger clusters and lower effective [tRNA] in the condensates at the highest values of [POS_L_] (Fig. S14). The effect of increasing [RP] is again a depletion of POS_L_ in the tRNA condensates, so that [tRNA] in the condensates increases with [RP] for a given value of [POS_L_] (Fig. S14).

In the simulations described so far, the total volume fraction of the system was kept constant by reducing the crowder (CRW) concentration as [POS_L_] and [RP] increased. Therefore, an alternative explanation of the decrease in [tRNA] inside the condensates with increasing [POS_L_] could be reduced crowding by the condensate environment. To test this further, we reduced [CRW] without changing [POS_L_]. Reduced [CRW] also led to reduced [tRNA] in the condensate, but the effect is much smaller than when [CRW] is reduced along with an increase in [POS_L_] (Fig. S15).

### tRNA condensation is a phase separation process

In order to construct phase diagrams, simulations of the five-component model phases were carried out at a range of temperatures for selected values of [RP] and [POS_L_]. Cluster size distributions were extracted (Figs. S16-S18) and the volume fractions of tRNA in dilute and condensed phases as a function of temperature were determined based on the number of tRNA outside and inside the largest tRNA clusters. The resulting curves (Fig. 3) show the typical features of phase diagrams with phase coexistence below critical temperatures *T_c_* of 400 to 535 K. In the absence of ribosomes, *i.e*. [RP] = 0, an increase in [POS_L_] lowers *T_c_* and narrows the two-phase regime (Fig. 3C). This is consistent with reentrant phase behavior expected for complex coacervation of a binary mixture. However, in the presence of ribosomes, i.e. [RP] = 55 μM, *T_c_* increased at the same time as the two-phase regime narrowed with increasing [POS_L_] (Fig. 3D). Moreover, when [POS_L_] = 180 μM, near the minimum needed for PS, an increase in [RP] slightly decreased *T_c_* (Fig. 3A/E), whereas, at a higher concentration, i.e. [POS_L_] = 880 μM, *T_c_* increased with increasing [RP] up to a maximum at 55 μM before decreasing (Fig. 3B/E). These observations reflect competition between ribosomes and tRNA for interactions with POSL and more generally highlight the effects of a complex interplay between interactions in non-binary mixtures that are more representative of biological environments than simple binary mixtures.

**Figure 3.**
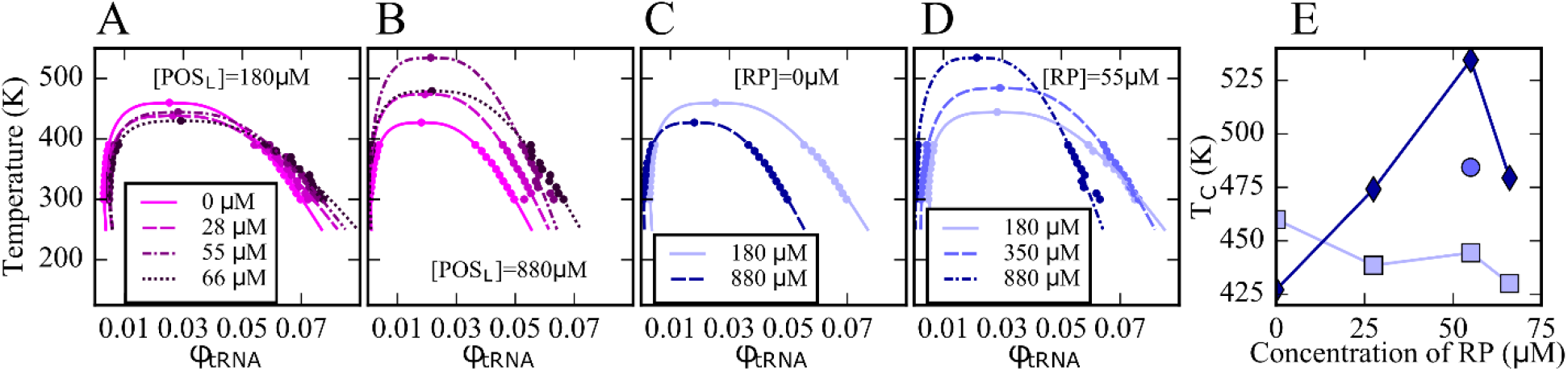
Phase diagrams for tRNA with [POS_L_] = 180 μM and varying RP concentrations (A); with [POS_L_] = 880 μM and varying RP concentrations (B); with [RP] = 0 at two [POS_L_] concentrations (C); and with [RP] = 55 μM and varying POS_L_ concentrations (D); critical temperatures as a function of [RP] at [POS_L_] =180 μM (squares), at [POS_L_] = 880 μM (diamonds), and at [POS_L_] = 350 μM (sphere) (E). Lines in A-D were fitted according to Eqs. M-6 and M-7 (SI Methods).

### Phase separation in experiments for binary mixtures of globular RNA and proteins

The results presented so far have focused on multi-component systems that were modeled to reflect the density and distribution of particle sizes and charges in cytoplasmic environments. A key prediction is that PS due to complex coacervation may occur for a wide range of nucleic acids and positively charged proteins simply based on electrostatic complementarity. To test this idea experimentally, we now turn to binary mixtures of globular RNA and positively charged proteins. We focused on the 47-nucleotide J345 Varkud satellite ribozyme RNA, that folds into an approximately globular shape (81) and that was mixed at high concentration with common proteins with positive charges and varying sizes for which we may expect PS: myoglobin (*q* = +2), trypsin (*q* = +6), lysozyme (*q* = +8), lactate dehydrogenase (LDH; *q* = +4), and alcohol dehydrogenase (ADH; *q* = +8). Bovine serum albumin (BSA; *q* = −17, *r* = 2.58 nm) was added as a control, for which condensate formation is not expected due to its negative charge.

**Figure 4.**
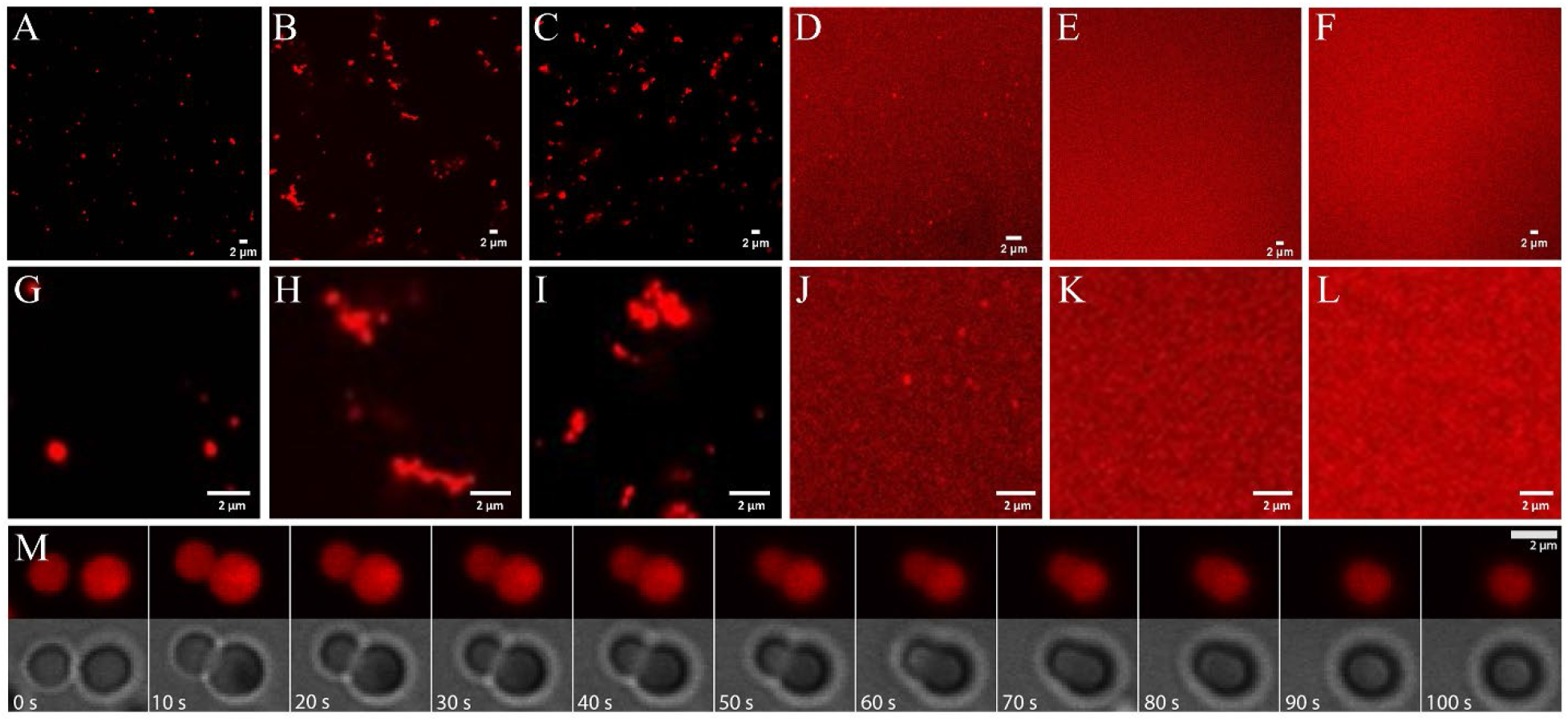
Phase separation in mixtures of J345 RNA at 0.45 mM and various globular proteins at 0.35 mM from confocal microscopy of labeled RNA: trypsin (A; G), ADH (B; H), lysozyme (C; I), LDH (D; J), myoglobin (E; K), BSA (F; L). Time lapse of droplet merging in RNA-trypsin mixture from fluorescence and bright-field microscopy imaging (M).

Imaging via confocal microscopy of dye-labeled RNA (Fig. 4 and Figs. S19 to S24) shows well defined fluorescent clusters for mixtures of RNA with trypsin, ADH, lysozyme, and LDH, but not for RNA with myoglobin or BSA. The background fluorescence varies significantly with protein. It is especially high for the mixtures with LDH, suggesting that only a fraction of RNA is participating in the condensates and a larger fraction of RNA remained in the dilute phase.

Individual condensates are relatively small, and many appear to have sizes near or below the diffraction limit of the microscope. For RNA-trypsin mixtures we clearly observe single droplet-shaped condensates of varying sizes that follow roughly an exponential distribution (Fig. S25).

We note that the concentration of Cy3-labeled RNA is only 8 μM, corresponding to 1 in 56 RNA at 0.45 mM total RNA concentration. Therefore, the fluorescent images in Fig. 4 are biased towards clusters that contain at least 50 RNA molecules, whereas smaller clusters are imaged incompletely. RNA-LDH condensates appear similar but we did not attempt a quantitative size analysis due to the high background fluorescence of the RNA-LDH sample. Diffusing droplets in the RNA-trypsin mixture merge over the course of 1 minute when they come into proximity (Fig. 4M and SI movies M-2 and M-3), indicative of liquid behavior inside the condensates.

For other proteins (lysozyme and ADH) we found more complex condensate morphologies (Fig. 4), where smaller condensates associate to form larger, irregular-shaped condensates without merging as seen for RNA-trypsin condensates. This suggests that the condensates with these proteins are less liquid-like.

To further study the particle size distributions, we carried out dynamic light scattering (DLS) analysis on RNA/lysozyme and RNA/trypsin samples (Fig. 5, Figs. S26-S27, and Table S2; see also SI Methods for analysis details). The light scattering correlation functions indicate a polydisperse sample that is dominated by very long correlation times up to 1 s (Fig. 5). Those long correlation times theoretically correspond to macroscopic-size particles (82), but since no such particles were readily visible in the sample, we may conclude that a significant fraction of condensates exhibited very slow diffusion due to surface adsorption. From the correlation function at shorter times, multi-exponential fits suggest particles in two size regimes for RNA-trypsin and in three regimes for RNA-lysozyme. In both cases, the data indicate the presence of 10 nm-scale particles that are consistent with oligomer-size clusters of RNA and protein molecules. Such small clusters between RNA and/or proteins are expected to be present in the dilute phase due to transient associations (83–86). In both, RNA-trypsin and RNA-lysozyme sample, the DLS analysis suggests the presence of μm-size particles (somewhat smaller for trypsin than for lysozyme). In addition, the DLS data indicate the presence of particles at the light microscopy diffraction limit, around 300 nm, for the RNA-lysozyme system but not for RNA-trypsin mixtures. In fact, the DLS results are qualitatively consistent with the microscopy images and provide additional insights into the particle size distributions at and below the light diffraction limit. However, an exact quantitative interpretation of the DLS results is challenging due to the polydispersity and dynamic nature of our samples and for that reason we also did not attempt to quantify what fraction of particles would be expected in the different size regimes.

**Figure 5.**
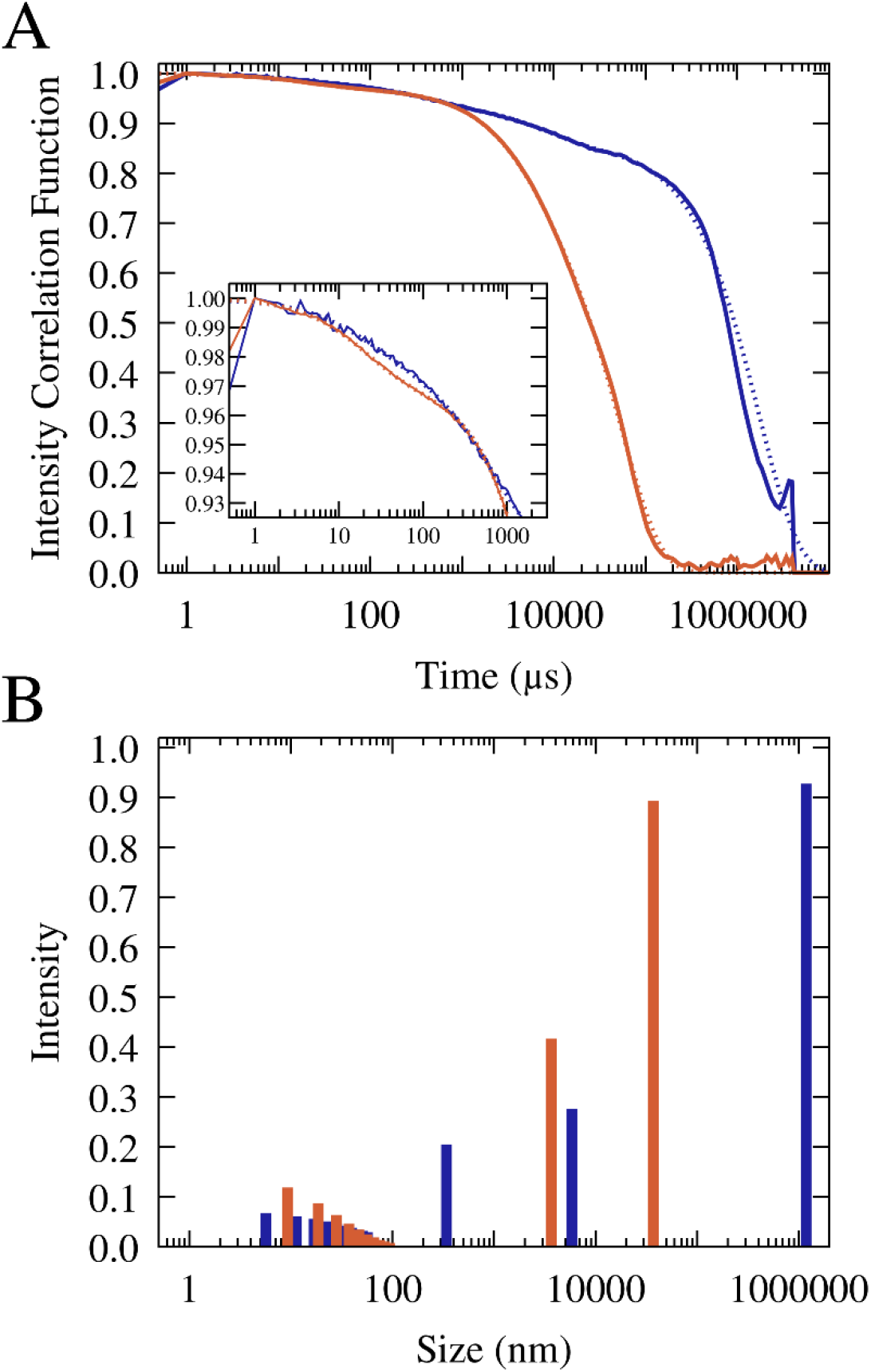
Normalized and averaged scattering intensity correlation functions from triplicate dynamic light scattering experiments of mixtures of 0.1 mM J345 RNA with 0.166 mM trypsin (orange) and 0.4 mM RNA with 0.675 mM lysozyme (blue) (A). Scattering intensity as a function of particle size from multi-exponential fits to the correlation functions (shown as dotted lines in A; see SI Methods) for trypsin (orange) and lysozyme (blue) (B).

To map out a phase diagram, we prepared RNA-trypsin mixtures at various, experimentally feasible RNA and protein concentrations. PS required a minimum protein concentration, e.g. with [RNA] = 100 μM, PS was found with [trypsin] = 150 μM but not with [trypsin] = 50 μM (Fig. S28). At the same time, PS was lost when RNA concentrations were too high. The resulting phase diagram based on confocal microscopy imaging is shown in Fig. 6 in comparison with results from theory that are discussed below.

**Figure 6.**
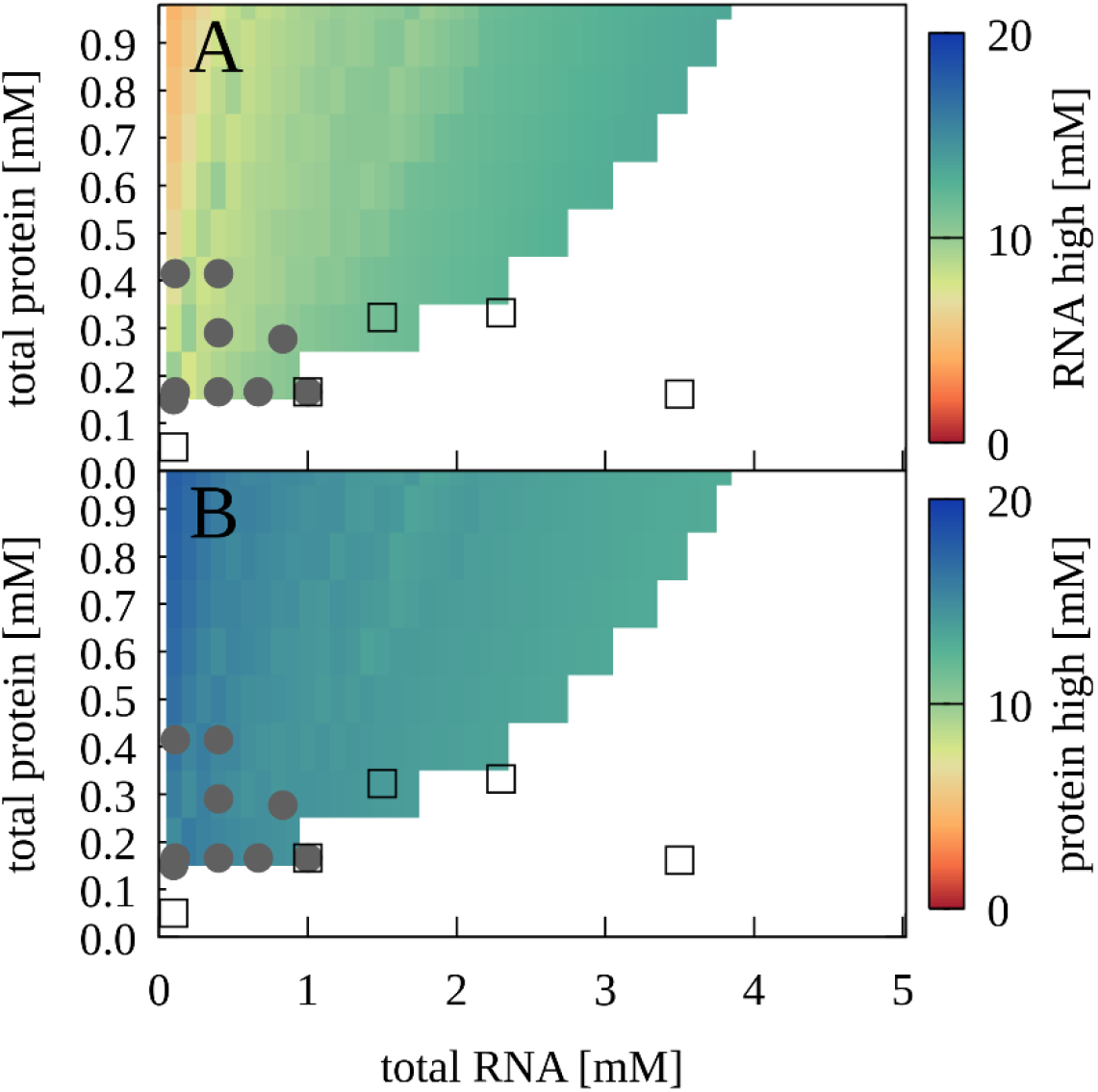
Phase separation for mixtures of J345 RNA and trypsin as a function of total protein and RNA concentrations from experiment and theory. Grey filled circles indicate concentrations for which phase separation was observed experimentally based on confocal microscopy; empty squares indicate concentrations for which microscopy imaging did not show phase separation. Colors indicate predicted concentrations from theory for RNA (A) and proteins (B) in the condensed phases. No phase separation is predicted for white areas.

Förster resonance energy transfer (FRET) experiments also showed a significant increase in FRET efficiencies from 50 μM to 150 μM (Fig. 7A). The comparison between the microscopy and FRET results furthermore establishes that RNA condensates at this RNA concentration can be recognized by FRET efficiencies above 0.26, whereas lower values may indicate a disperse phase. The gradual increase in FRET efficiencies from 0.24 to 0.26 upon increase of trypsin concentrations from 0 to 50 μM is interpreted to result from increasing non-condensate cluster formation (see cluster size distributions in Fig. S10 at [RP] = 0 with increasing protein concentration). However, as in the confocal microscopy experiments, the low concentration of fluorescence-labeled RNA limits the detection of very small clusters where only one or zero of the RNA would be labeled. The FRET results are compared with theoretical predictions (Fig. 7B) as detailed below.

**Figure 7.**
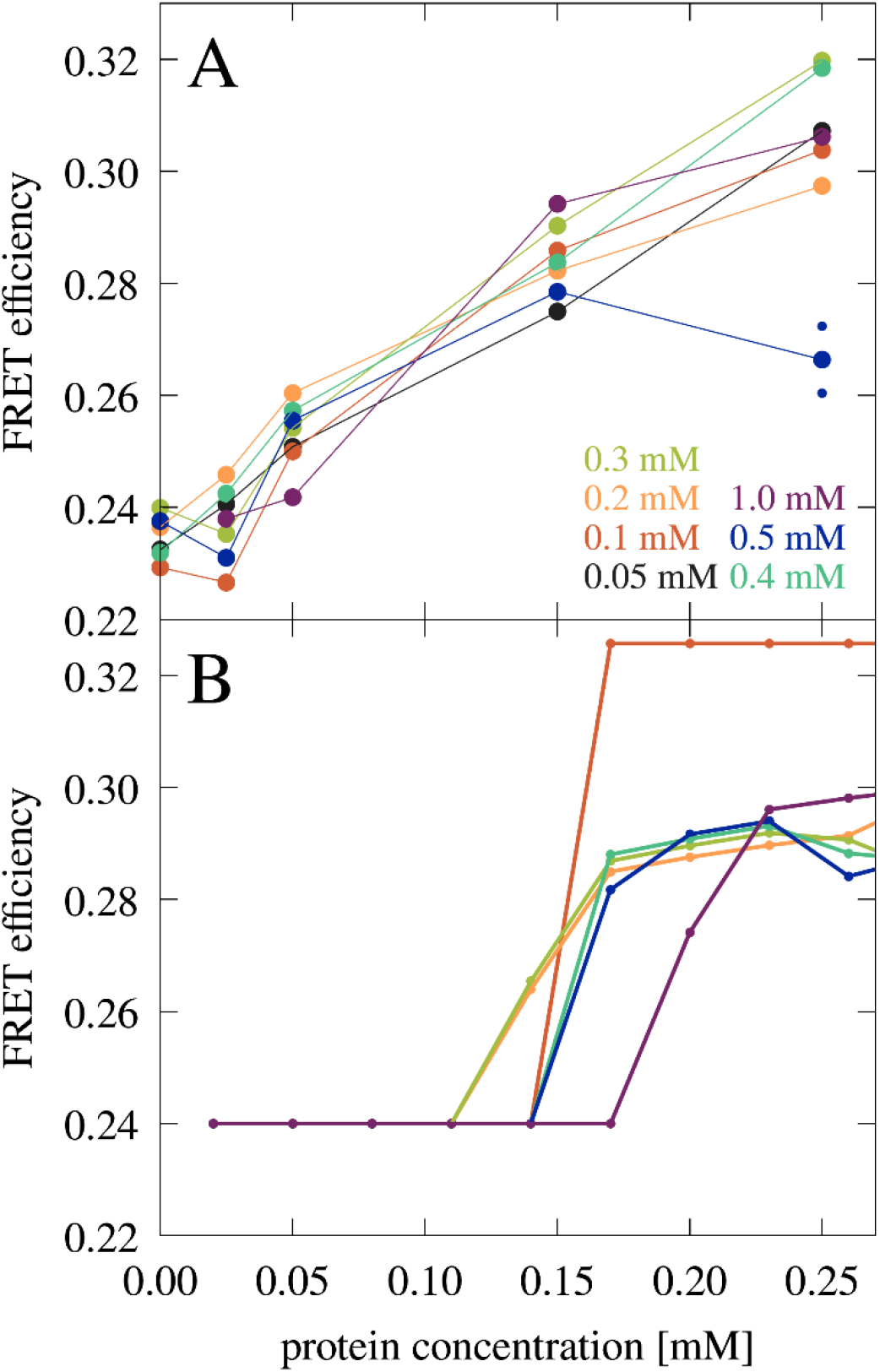
FRET efficiency in mixtures of J345 RNA with trypsin as a function of protein concentration at different RNA concentrations (as indicated by color). The average of two measurements is shown for 0.25 mM protein and 0.5 mM RNA concentrations with smaller points indicating individual measurements. (A). FRET efficiency estimated from the fraction of RNA in the condensed phase from theory (SI Theory) (B). In every measurement, the concentration of Cy3- and Cy5-labeled RNA is constant, 8 μM and 42 μM, respectively.

We applied circular dichroism (CD) and nuclear magnetic resonance (NMR) spectroscopy with the goal of examining whether the proteins and RNA retain their folded states upon condensate formation. The CD spectra in Fig. S29 show that there is no substantial change in the shape of the spectrum of trypsin in the presence of the RNA from 225-250 nm, which would be expected if the protein had unfolded, as a random coil spectrum has essentially no ellipticity in this wavelength range and the spectrum. The key feature of the RNA spectrum, i.e. the broad peak at 250-290 nm is also retained in the mixture. In fact, the spectrum of the trypsin-RNA mixture appears to be simply a linear combination of the spectra of each of the components measured separately.

NMR spectroscopic analysis of RNA-trypsin and RNA-lysozyme samples at PS-inducing concentrations focused on the structure of the RNA. We observed the characteristic ^1^H spectrum of a solution containing only J345 RNA that matches previously matched spectra for the same structure (81) (Fig. S30). In the presence of proteins, the characteristic peaks were retained at the same positions, although with greatly attenuated intensities (Fig. S30). This was interpreted to mean that only a fraction of RNA remained sufficiently dynamic to achieve rotational averaging via molecular tumbling. From comparing the signal-to-noise ratios we estimate that about 80% of the RNA is not visible in the RNA-lysozyme sample and 90% is invisible in the RNA-trypsin sample. Since the majority of RNA is expected to be found in the condensates, this suggests that rotational diffusion of individual RNA molecules in the condensates is retarded significantly since the condensates themselves are too large (>100 nm) to tumble on time scales allowing NMR signals to be observed (<100 ns). Moreover, if one assumes that only RNA in the dilute phases remains visible in NMR spectroscopy, the experiments provide an estimate of the fraction of RNA in the dilute vs. condensed phases, i.e. 20:80 in the presence of lysozyme and 10:90 in the presence of the trypsin for the concentrations studied here. Unfortunately, that also implies that there is no information about the structure of RNA inside the condensates from these experiments.

### Phase separation of RNA and proteins described by simulations and theory

To compare with the experimental findings, we carried out CG simulations again with the model described above but for binary mixtures of spherical particles equivalent in size and charge to the experimentally studied systems, *i.e.* J345 RNA (q=-46, r = 1.47 nm), myoglobin (*q* = +2, *r* = 1.64 nm), trypsin (*q* = +6, *r* = 1.81 nm), lysozyme (*q* = +8, *r* = 1.54 nm), lactate dehydrogenase (LDH; *q* = +4, *r* = 2.68 nm), alcohol dehydrogenase (ADH; *q* = +8, *r* = 2.79 nm), and bovine serum albumin (BSA; *q* = −17, *r* = 2.58 nm). We also tested a spherical particle equivalent to cytochrome C (*q* = +11, *r* = 1.45 nm) which was not studied experimentally because of heme absorption. We observed the formation of condensates at sufficiently high salt concentrations. With κ = 0.7 (about 20 mM salt, see SI Methods), condensates formed with lysozyme, trypsin, LDH, and ADH, but not with cytochrome C, myoglobin, or BSA (Fig. 8). Very similar results were also found with an alternative effective charge model (according to Eq. M-3) as shown in Fig. S31.

The simulation results qualitatively match the experimental results in terms of which proteins promote PS. Moreover, the fraction of RNA in the dilute phase is higher with lysozyme than with trypsin (32% vs. 26-27% using Eq. M-2 or M-3 from averages over the last 100 μs) in qualitative agreement with the estimates from the NMR experiments. We note that an overall larger fraction of RNA is expected in the dilute phase in the simulations due to an excess concentration of RNA (0.439 mM) compared to the protein concentration (0.350 mM) whereas concentrations of RNA and protein were equal in the NMR experiments (0.150 mM). However, the scale of the simulations is too small to directly compare the condensate sizes with the experimental size distributions.

**Figure 8.**
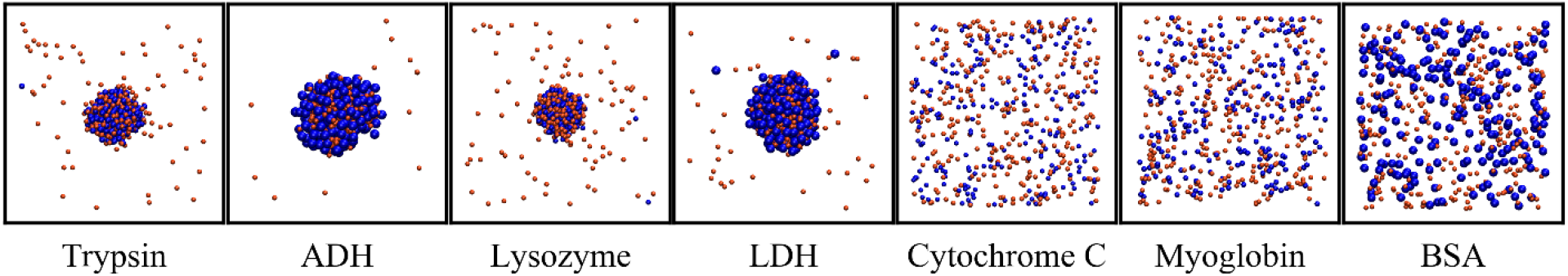
Snapshots after 1 ms for binary RNA-protein mixtures at T = 298K, with κ = 0.7 and using effective charges according to Eq. M-2. [RNA] = 0.493 mM and [protein] = 0.350 mM. Orange and blue spheres show RNA and proteins, according to size. Concentrations inside the condensates were [RNA:lysozyme] = 20.2:20.2 mM; [RNA:trypsin] = 16.5:15.2 mM; [RNA:LDH] = 9.6:7.2 mM; [RNA:ADH] = 9.5:6.7 mM.

To generate more extensive phase diagrams, a theoretical model was developed based on the CG simulations. Briefly, the model approximates the chemical potential for either RNA or proteins in condensed and dilute phases based on a decomposition into enthalpy and entropy: μ = *Δh* - *TΔs*. The enthalpy is determined from convoluting the coarse-grained interaction potential *U(r)* (Eq. 3) with radial distribution functions 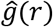 of RNA-RNA, RNA-protein, and protein-protein interactions in the condensed and dilute phases extracted from CG simulations and scaled by particle densities *ρ* (see SI Theory):

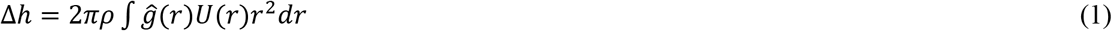

The entropy was estimated from the ratio of particle densities ρ between the entire system and either the dilute or condensed phase:

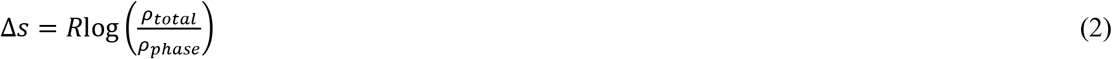

Solutions with respect to the concentrations of protein and RNA in dilute and condensed phases were determined numerically under the conditions that *μ_condensed_ = μ_dilute_* for either RNA, protein, or both, and that molecular volume packing fractions did not exceed maximum liquid packing densities (SI Theory). Total free energies were then calculated, taking also into account mixing entropy contributions between RNA and protein particles. PS was predicted based on the solution with the lowest free energy.

The theoretical approach is essentially a variation of Voorn-Overbeek theory (87) for spherical particles. While this theory has seen numerous applications, especially to polyelectrolyte fluids (88, 89), the specific model described here emphasizes an interaction potential that is parameterized based on atomistic simulations of biological macromolecules and that was further tuned to match experimental data. Therefore, the theory is expected to make predictions that are more relevant for globular biological macromolecules than previous studies.

In developing the theory, we found that using the alternative effective charge model according to Eq. M-3 (Fig. S32) results in better agreement between theory and experiment and therefore we used this model here. We also use a slightly different Debye-Hückel screening term, *i.e.* κ = 1.17, which gave better agreement between theory and experiment.

Application of the theory predicts that PS should occur for a wide range of protein radii and charges as long as proteins are large enough and carry sufficiently positive charge (Fig. 9). More specifically, radius/charge combination corresponding to lysozyme, trypsin, LDH, and ADH are predicted to lead to PS as in the experiments and CG simulations. The radius and charge corresponding to myoglobin is just outside the PS region (Fig. 9) again consistent with the lack of PS in the experiment and simulations. The theory also predicts PS for cytochrome C, for which PS was not seen in the simulations.

The theory reproduces an expected temperature dependence of PS with protein-dependent critical maximal temperatures (Fig. S33). The electrostatic nature of PS also suggests that changes in salt concentrations would affect the findings and the results are indeed sensitive to the value of κ. However, the theoretical treatment is too limited due to the mean-field nature of the Debye-Hückel formalism to make meaningful predictions of salt effects. More specifically, the model is only valid for low ionic strengths and ignores entropic consequences of ion partitioning between condensed and dilute phases that are an important contribution to PS in complex coacervates (90).

Using the theory, we constructed concentration-dependent phase diagrams that can be compared with experiment. Fig. 6 shows the prediction of the two-phase region for RNA-trypsin in good agreement with the experimental data. Figs. S34-S39 show the phase diagrams for all proteins studies here over a wider range of concentrations. All phase diagrams exhibit reentrant behavior with minimal and maximal protein and RNA concentrations as expected for complex coacervates. It should be noted, though, that the full range of concentrations cannot be realized in practice for all systems due to limited solubilities.

Predictions from the theory also allowed a quantitative interpretation of the FRET experiments. Using the predicted fraction of RNA in the condensates for the RNA-trypsin mixtures at different RNA and protein concentrations (Fig. S40) FRET efficiencies were estimated (Fig. 7B and SI Theory). The theoretical predictions qualitatively reproduce the experimental data with an onset of increased FRET efficiencies due to condensation. Moreover, the gradual increase in FRET efficiencies after condensates form is predicted from a growing number of RNA in the condensed phase as protein concentration increases.

**Figure 9.**
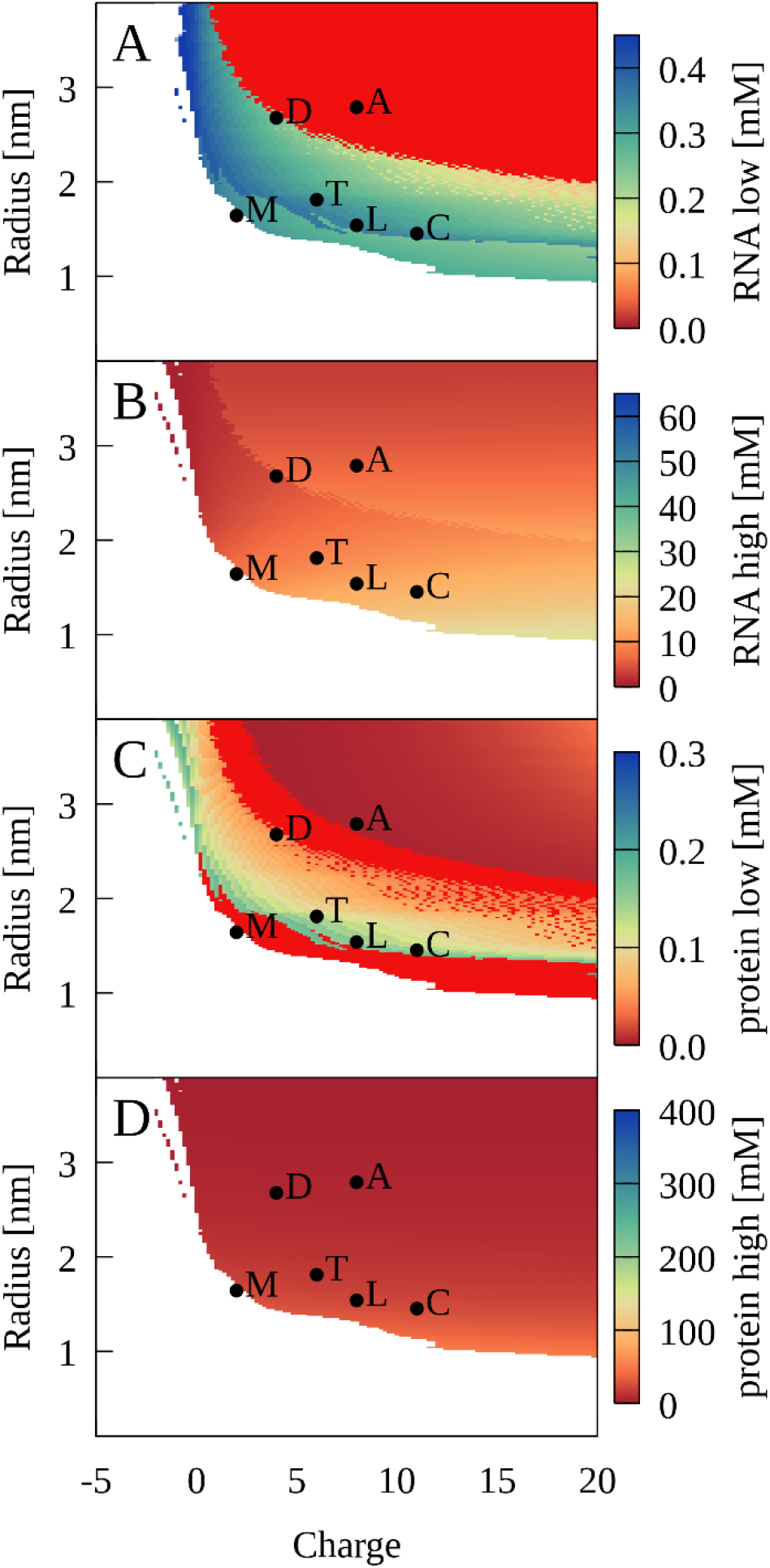
Phase separation for binary RNA-protein mixtures as a function of protein charge and radius from theory. Colors show [RNA] (A, B) and [protein] (C, D) in dilute (A, C) and condensed (B, D) phases. Red indicates zero concentration. [RNA] = 0.45 mM, [protein] = 0.35 mM, κ = 1.17, and T = 298 K. Corresponding properties for proteins are denoted as follows: myoglobin (M); trypsin (T); lysozyme (L); cytochrome C (C); LDH (D); ADH (A).

## DISCUSSION

This study presents a general view on charge-driven biomolecular PS supported by simulation, theory, and experiments. More specifically, we report a potential for PS between negatively charged RNA and positively charged proteins without requiring polymer-character of either component or specific binding interactions. Our simulations and the theoretical model are based on isotropic spheres, whereas experimental validation is based on a compact, approximately globular RNA and a variety of globular proteins that are not known to specifically interact with RNA. This implies that PS may be a very general phenomenon in biological cells depending on the concentrations, charge, and size distribution of available nucleic acid and protein components. In fact, our simulations of a bacterial cytoplasm provide examples of separately forming tRNA-protein and ribosome-protein condensates involving a variety of proteins in a cytoplasmic environment.

The idea of strong complementary electrostatic interactions playing a major role in PS via complex coacervate formation is well-established for a variety of different molecules (25, 27–31) and also for PS involving biomolecules (91). While almost all of the LLPS studies to-date involve polymers and in particular IDPs (42), there are also examples in the literature that discuss PS involving folded proteins (44, 45, 56, 62, 92, 93). In most of those cases, the ability to form condensates is generally ascribed to specific multi-valent interactions and evidence for a more generic electrostatic-only mechanism are only just beginning to emerge (29, 44). The results presented here provide evidence for a more general principle that does not require flexible polymers, specific interaction sites, or specific secondary structures (93). The central principle is simply electrostatic complementarity at the molecular level, but a more generalized concept of multi-valency is implicitly assumed. Isotropic spheres without any directional preference for interactions are in fact infinitely multi-valent. On the other hand, globular proteins with basic amino acids distributed widely across their surface and diffuse positive electrostatic potentials over most of the molecular surface (Fig. S41) are effectively poly-valent particles with respect to interactions with nucleic acids. The key insight from this study is that proteins not known to interact specifically with nucleic acids under dilute conditions may form condensates with nucleic acids, if the proteins are present at sufficient amounts, simply based on a principle of generic poly-valency and an overall charge attraction.

Our study suggests that size and charge are essential determinants of PS between RNA and proteins. Favorable condensates require optimal packing and a balance of attractive and repulsive interactions between oppositely charged RNA and protein particles. Fig. S42 shows a snapshot from the cytoplasmic system illustrating how such packing may be achieved. The optimal balance depends on the size of the RNA particles: Larger proteins are required for the smaller RNA molecules to phase separate whereas smaller proteins allow the larger ribosomal particles to phase-separate (Fig. S5). This can be seen more clearly in the five-component model system, where a relatively modest reduction in the radius of the larger positively charged particle leads to a loss of close tRNA contacts (Fig. S43), therefore preventing condensate formation. The theoretical model for binary RNA-protein mixtures also predicts a minimum protein radius for PS, at least at lower charges (Fig. 9). Myoglobin is outside the predicted range and although it has a net-positive charge, PS was not observed in the experiment at protein concentrations below the RNA concentrations (Fig. 4) consistent with the theory.

The total concentration of the protein is in fact another determinant for PS. Simulations and theory predict minimum protein concentrations depending on the protein charge and size around 0.05 mM or more (Figs. 34-39). For trypsin, this was validated experimentally via microscopy and FRET spectroscopy (Figs. 6 and 7). While many cellular proteins may not be present individually at such high concentrations, our cytoplasmic model shows that a heterogeneous mixture of similar-sized and similar-charged proteins may promote PS equally well. At the lower end, the RNA concentration appears to be a less critical factor for observing PS, although a larger amount of RNA allows more numerous and larger condensates to form assuming that there is enough protein available, at least until reaching a critical RNA concentration beyond which PS is not favorable anymore. In binary mixtures, this is simply a question of the total protein concentration. In the heterogeneous cytoplasmic model, we found competition for the larger positively charged proteins by the ribosomes forming their condensates to be another factor affecting tRNA condensate formation that would need to be considered in cellular environments (Fig. S14).

Since electrostatics is a major driving force of the PS described here, changes in salt concentration are expected to alter the tendency for PS. The theory applied here is not well-suited to examine variations in the salt concentration. At the same time, there is only a limited range of decreased salt conditions that can be applied before either the RNA or the protein structures become destabilized. Therefore, an accurate quantitative understanding of how salt effects may affect the results presented here will have to be deferred to future studies.

A significant interest in PS in biology is related to liquid-state condensates. Such condensates would maintain the dynamics that is necessary for many biological processes as opposed to dynamically retarded gels or amorphous clusters. The simulations suggest that the condensates retain a dynamic character based on calculated self-diffusion rates, albeit with serious limitations on diffusion estimates from CG simulations, especially in the absence of hydrodynamic interactions (94). In experiment, we find evidence of liquid-like behavior for condensates formed in RNA-trypsin mixtures, but the dynamic properties of RNA or proteins in other RNA-protein condensates are less clear and the NMR spectroscopy results generally suggest significant retardation of diffusional dynamics inside the condensates.

Although there are some limitations in the current study that will need to be revisited in future studies to gain a more detailed understanding of the more universal PS between RNA and proteins described here, the main advantage of the CG models and theory is that its simplicity allowed us to explore the large spatial scales and long time scales that can predict phase behavior on experimentally accessible scales. Although the CG models were parameterized based on high-resolution atomistic simulations of concentrated protein solutions, these models lack all but the most basic features of biological macromolecules. Increased levels of realism could be achieved without too much additional computational cost via patchy particles (95), whereas higher-resolution in the form of residue-based coarse-graining (42) to explore the effects of shape anisotropy and inhomogeneous charge distributions across RNA and protein surfaces is in principle attainable but computationally much more demanding. Finally, we expect that further insights could be gained from atomistic simulations of RNA-protein clusters initiated from configurations in the CG simulations to better understand the detailed molecular interactions stabilizing the condensates. On the experimental side, we only focused on RNA without visualizing protein condensation. Moreover, there is a need to follow up on this work with *in vivo* studies to establish how ubiquitous the condensates described here are under cellular conditions.

## CONCLUSIONS

We report phase separation of RNA and proteins based on a universal principle of charge complementarity that does not require polymers or multi-valency via specific interactions. The results are supported by coarse-grained simulations, theory, and experimental validation via microscopy, FRET, and NMR spectroscopy as well as DLS experiments. Condensate formation depends on concentration, size, and charge of the proteins but appears to be possible for typical RNA and common proteins. Simulation results furthermore suggest that such phase separation may occur in heterogenous cellular environment, not just between tRNA and cellular proteins but also, in separate condensates, between ribosomes and proteins. Further computational and experimental studies are needed to gain more detailed insights into the exact molecular nature of the condensates described here.

## METHODS

### Coarse-grained simulations

CG simulations were run using a modified version of a previously introduced colloid-type spherical model (96). In this model, pair interactions consist of a short-range 10-5 Lennard-Jones potential and a long-range Debye Hückel potential according to:

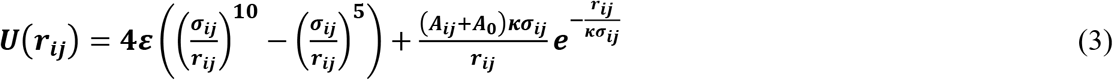

where *r_ij_* is the inter-particle distance, *σ_ij_* is the distance between particles at which the potential is zero, *ε* is the strength of short-range attraction, *A_ij_+A_0_* describes attractive or repulsive long-range interactions, and *κσ_ij_* is the Debye-Hückel screening length. Only *A_ij_* and *σ_ij_* vary between different particles according to charge and size (SI Methods).

Most systems involved box sizes of 100 nm and simulations were extended to 1 ms via Langevin dynamics using OpenMM (97) on graphics processing units (GPUs). The CG simulations were analyzed via in-house scripts and the MMTSB Tool Set (98) (SI Methods).

### Experimental materials and methods

The J345 RNA sequence was synthesized and deprotected by Dharmacon (Horizon Discovery Group), both with and without Cy3 or Cy5 on the 3’ end. The 47-base sequence is GCAGCAGGGAACUCACGCUUGCGUAGAGGCUAAGUGCUUCGGCACAGCACAAGCCCGCUGCG

All measurements were made using the buffer used by Bonneau and Legault for structure determination of this sequence, 10 mM sodium cacodylate (pH 6.5), 50 mM NaCl, .05% sodium azide, 5 mM MgCl_2_. Equine liver trypsin, equine alcohol dehydrogenase, bovine lactic dehydrogenase, equine myoglobin, hen egg lysozyme and bovine serum albumin were obtained from Sigma-Aldrich and used without further modification.

Confocal microscopy images were obtained on a Nikon A1 scanning confocal microscope with 100x magnification. The excitation wavelength was 561 nm and detection was set for Cy3 fluorescence using a GaAsP detector. The diffraction-limited spatial resolution is 260 nm. Images were processed with ImageJ and modified only for contrast and brightness. Images were cropped and enlarged to aid observation of the smallest features.

Fluorescence spectra were obtained with PTI Q4 fluorimeter, excited at 475 nm and emission observed between 525 and 700 nm. The concentration of Cy3-labeled RNA and Cy5-labeled RNA were kept constant at 8 uM and 42 uM, respectively, with the unlabeled concentration varied from 0 to 0.5 mM. The low concentration of labeled RNA limits the possibility of self-quenching but also limits the detection of very small clusters.

The normalized FRET ratio was calculated from the total intensity between 525 and 650 nm for the donor and 650 and 700 nm for the acceptor,

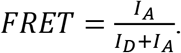

In the absence of protein, the RNA exhibits some baseline transfer, likely due to transient interactions between the dyes, leading to a background FRET level of ~0.24. Upon the addition of protein above the threshold concentration, the mixture is visibly turbid.

The size distribution of the protein-RNA complexes were measured using a dynamic light scattering (DLS) machine (Zetasizer nano series from Malvern company) at room temperature. The samples were mixed freshly before each experiment and all measurements were repeated three times in a single run and the corresponding average results were reported. A Helium Neon laser with a wavelength of 632 nm was used for the size distribution analysis.

NMR spectra were acquired at a ^1^H frequency of 600 MHz on a Varian 600 MHz spectrometer with a room-temperature probe. Solvent was suppressed with a gradient 1-1 echo sequence. Samples were prepared in 90% H_2_O, 10% D_2_O in the buffer described above with DSS as an internal chemical shift reference. 16k points were acquired with a 1-second recycle delay and a total acquisition time of approximately one hour per spectrum. RNA concentrations were 300 μM for J345 only and 135-140 μM for RNA-protein samples; protein concentrations were around 150 μM; the RNA-only spectrum was scaled to account for the differing concentration. Spectra were processed with zero-filling to 32k and a 5 Hz exponential window function.

Circular dichroism measurements were made using an Applied Photophysics Chirascan spectrometer. All measurements were made using a 0.1 mm pathlength cuvette at room temperature.

## Supporting information

SI Sheet 1

SI Sheet 2

## ACKNOWLEDGMENTS

This study was funded by the National Science Foundation (MCB 1817307, to MF and LJL; MCB 2018296, to CGH) and the National Institutes of Health (R35 GM126948, to MF). Computational resources at the Institute for Cyber-Enabled Research/High Performance Computing Cluster (ICER/HPCC) at Michigan State University were used.

