## Supplementary Text, Figures, and Table for "Charge-Driven Condensation of RNA and Proteins Suggests Broad Role of Phase Separation in Cytoplasmic Environments"

**This PDF file includes:**

Supplementary methods
Supplementary theory
Figures S1 to S49
Tables S1 and S2
Caption for movies S1-S3
Captions for spreadsheets S1 and S2
References for SI reference citations

**Other supplementary materials for this manuscript include the following:**

Movies S1 to S3
Spreadsheets S1 and S2

### Supplementary Methods

**Coarse-grained model.** Pair-wise interactions were calculated according to Eq. 3 with the parameters $\varepsilon$, $\sigma_{ij}$, $A$, $A_0$, and $\kappa$. The model was initially parameterized from previously published all-atom simulations of homogeneous mixtures of chicken villin headpiece ("villin") (1) and subsequently validated with heterogeneous mixtures of protein G, villin, and ubiquitin (2) as summarized in Table S1.

A common value of $\varepsilon$ = 4.0 kJ/mol was used for all particles in the short-range 10-5 Lennard-Jones potential. Particle size was taken into account by first determining the radii $r_i$ of spheres with equivalent volumes to the atomistic molecular volumes of a given macromolecule or complex (see for example Sheet S1 for molecules in the cytoplasmic model system). Lennard-Jones parameters $\sigma_i$ were obtained from the radii $r_i$ according to:

$$\sigma_{\mathrm{i}} = 2^{-\frac{1}{6}} \cdot r_i \quad \text{M-1}$$

Pairwise parameters $\sigma_{ij}$ were calculated as $\sigma_{ij} = \sigma_i + \sigma_j$.

In the long-range Debye-Hückel type potential, a common value of $A_0$ =3.0 kJ/mol was used to reflect effective repulsion between charge-neutral, but still polar molecules due to solvation effects. Net charges led to additional repulsive or attractive contributions.

The nominal net charge of a given molecule was converted to effective charges to account for counterion condensation around highly charged macromolecules (3). We distinguish here effectively bound ions that lead to an effectively reduced charge vs. ions that remain mobile in solution and give rise to Debye screening as described below. Generally, the effective charge remains close to nominal charges for small charges, but for highly charged molecules, in particular negatively charged nucleic acids and nucleic acid complexes such as the ribosome, the effective charge is reduced significantly (4-6). Charge neutralization is more pronounced with divalent ions such as $Mg^{2+}$ vs. monovalent ions such as $Na^+$ or $K^+$ (4, 7, 8). But the amount of $Mg^{2+}$ ions in biological systems is limited and typically not high enough to neutralize the charge of all the nucleic acids so that additional charge neutralization by monovalent ions remains a significant factor (9).

Here, we obtain effective charges via either one of the following expressions:

$$q_{\mathrm{eff},1} = \mathrm{sign}(q) \cdot 20 \cdot \log\left(\frac{|q|}{20} + 1\right) \quad \text{M-2}$$

$$q_{\mathrm{eff},2} = \mathrm{sign}(q) \cdot 0.6\sqrt{|q|} \cdot \log\left(\frac{|q|}{2} + 1\right) \quad \text{M-3}$$

Both empirical formulae give effective charges close to nominal charges for molecules with small charges and highly reduced charges for molecules with large formal charges (Fig. S32). For a DNA molecule with a nominal charge of -45, atomistic MD simulations suggest effective charges of -10 to -20 under the assumption that ions within 1 nm from the solute surface are effectively bound (7, 8); at the other end, effective charges between -100 to -800 are estimated for ribosomal particle with a nominal charge of about -4000 based on colloid models (4) or electrostratic potential calculations (6), assuming a mixture of divalent and monovalent ions is involved in neutralization. Equations M-2 and M-3 are both consistent with these estimates. Eq. M-2 was used initially and screens smaller charges less and larger charges more strongly compared to Eq. M-3 which was adopted after adjusting the theory to better match experimental results. Neither expression considers ionic concentration as counterion condensation does not depend strongly on concentration (8). Moreover, negatively and positively charged solutes are treated in the same manner even although the binding strength of biological anions ($Cl^-$) and cations ($K^+$, $Na^+$, $Mg^{2+}$) to oppositely charged macromolecules may be asymmetric. However, since highly positively charged macromolecules are uncommon, this assumption may not have significant consequences for the systems studied here.

The effective charges calculated either via M-2 or M-3 were then converted to $A_i$ values:

$$A_i = \text{sign}(q_i)\sqrt{\frac{3}{4}q_{i,\text{eff}}} \qquad \text{M-4}$$

Pairwise values were determined as $A_{ij}=A_i*A_j$ and the factor ¾ was determined by parameterization against the atomistic MD simulations.

The Debye screening length in Eq. 3 is $\kappa\sigma_{ij}$, *i.e.* it depends on particle size as in the original model by Mani et al. (10) in order to better model screening interactions between particles of very different sizes with screened charges that are mostly near the surface. This complicates interpretation of κ in terms of specific salt concentrations. However, as an illustration one may consider a typical smaller protein or RNA with $\sigma_{ii}$ = 3 nm where κ = 0.5, 1.0, and 1.5 would correspond to monovalent ion concentrations of 40 mM, 10 mM, and 5 mM, respectively. Note, that these ion concentrations reflect excess ion concentrations after subtracting condensed counterions as those are accounted for in the effective charges according to Eq. M-2 or M-3. Therefore, total ion concentrations in experiment corresponding to a given value of κ in our model should be significantly higher by factors of 2 to 10 depending on the charges of the considered macromolecules.

**Molecular dynamics simulations.** MD simulations were run up to 1 ms using OpenMM (11) on GPU machines. The interaction potential from Eq. 3 was implemented as a custom non-bonded interaction potential via OpenMM's Python interface. A Langevin thermostat was applied with a temperature of 298 K unless noted otherwise and a friction coefficient of 1 $ps^{-1}$. A value of κ = 1.5 was used to describe salt screening unless noted otherwise. The timestep for the simulations was set to 1 ps. Frames were saved every 1 ns for simulations of the 100 nm cytoplasm model, every 10 ns for the concentrated protein simulations used for parameterization, and every 100 ns for all other systems. The pairwise potential in Eq. 3 was evaluated with a cutoff 49.5 nm. A switching function was applied to be effective at 49 nm. In total, about 270 ms of combined simulation time was run for all systems described here. The total computational cost for these simulations was around 350 GPU days based on timing on a single NVIDIA GeForce GTX 1080 Ti GPU card.

**Validation of CG model.** For validation, CG simulations of the systems with the same concentrations as in the atomistic simulations were performed for 100 μs. The CG simulations compared favorably with the atomistic simulations based on pairwise radial distribution functions and cluster size distribution (Fig. S44).

**Bacterial cytoplasm model.** We constructed a coarse grained model of *Mycoplasma genitalium* cytoplasm based on our previously established atomistic model (12, 13). All the macromolecules and complexes were converted to single spherical particles where the particle center initially coincided with the center of mass of the molecules in the atomistic model. Sphere radii were determined as described above based on equivalent volumes, and effective charges were determined from nominal charges according to Eq. M-2 or M-3. A list of all particles with their size, charge, effective charge and concentration is given in the SI Sheet 1. The initial system is a cubic box with a size of 100 nm. Additional systems were generated with 200 and 300 nm box sizes by replicating the initial system accordingly. MD simulations were run up to 1 ms as described above.

**Five-component systems.** A representative model of the cytoplasmic system consisted of five components, with an effective charge and volume fraction matching the values in the cytoplasmic system. The components consist of tRNA, ribosomes (RP), positively charged proteins with small ($POS_S$) and large ($POS_L$) sizes and charges and neutral crowders (CRW) (SI Sheet 2). tRNA and RP have the same size and charge as in the full cytoplasmic system. The RP concentration includes RP molecules in the cytoplasmic model as well as the ribosomal fragments RR23, R50P RR16 and R30P (see SI Sheet 1). The tRNA concentration was adjusted to include all particles with a nominal charge between -100 to -25, except for GroEL, which has a very large size and was not found as part of the tRNA condensates in the cytoplasmic simulations. Concentrations of the positively charged proteins were adjusted to keep the total effective charge of the system close to the cytoplasmic model. The system components were then varied to achieve different concentrations of RP and positively charged particles (see SI Sheet 2). Simulations of the five-component system were performed as described above over 1 ms using only effective charges calculated via eq. M-2.

**Two-component systems.** Two-component RNA-protein systems were simulated with the same CG model as described above for 1 ms to make predictions for experimentally testable systems. Effective charges were calculated either via eq. M-2 or M-3. RNA particles were modeled after the 47-nucleotide J345 Varkud satellite ribozyme RNA, that folds into an approximately globular shape(14) with $r_{RNA}$ = 1.47 nm and $q_{RNA}$ = -46. Proteins were considered with the following charges and radii: myoglobin (+2, 1.64 nm), trypsin (+6, 1.81 nm), lysozyme (+8, 1.54 nm), cytochrome C (+11, 1.45 nm), lactate dehydrogenase (+4, 2.68 nm), alcohol dehydrogenase (+8, 2.79 nm), and bovine serum albumin (-17, 2.58 nm).

**Analysis tools.** Analysis of the CG simulations was performed for the simulation time between 500 µs to 1 ms unless stated otherwise using in-house code in conjunction with the MMTSB Tool Set (15).

**Clustering analysis.** We previously analyzed macromolecular clustering using specific distance cutoffs that were suitable for capturing direct molecular interactions leading to transient clusters (1, 2, 16). From those studies, we arrived at a definition of clusters based on contacts where center of mass distances between spherical particles are less than $\sigma_{ij}$ + 0.7 nm. $\sigma_{ij}$ is the pair-wise Lennard-Jones parameters in Eq. 1 defined as described above in Eq. M-1. This criterion was applied to all pairs of particles, of same or different type, and connected graphs were generated from the pairs determined to be in contact. All particles within such a graph were then considered to be part of one cluster.

We initially applied this criterion here as well in a slightly modified version where we only considered contacts based on tRNA-protein and RP-protein pairs in order to be able to separately analyze tRNA and RP clustering in the same system. GroEL-protein pairs were also included when analyzing RP clusters since they were found to associate on the surface of the RP-rich condensates. We found that the $\sigma_{ij}$ + 0.7 nm contact criterion underestimated cluster sizes when visually inspecting condensed states (Fig. S45). This may not be surprising since macromolecules in liquid condensates are not necessarily in direct contact with other molecules while direct interactions are the essential feature of the transient molecular clusters described by us previously. From inspecting radial distribution functions for interactions between tRNA and $POS_L$ and $POS_S$ particles in the five-component system at different concentrations, we found that an increased cutoff of $\sigma_{ij}$ + 2.2 nm would include all the contacts within the first peak (Fig. S46).

We further validated whether this criterion is more generally applicable to the cytoplasmic system by comparing with results from geometry-based scale-free hierarchical clustering. We applied such an algorithm to just tRNA particles during the last 100 µs of the simulation of the cytoplasmic systems so that clusters could be defined without having to invoke any contact-based criteria and without having to define clusters via interactions with other system components. We used the hierarchical clustering method implemented in the MMTSB Tool Set (15), but with a more recently established criterion for determining the optimal number of clusters (17). This approach gave fluctuating cluster sizes between 180 and 260 tRNA molecules with a peak near 240 molecules (see Fig. S47). Clusters based on the $\sigma_{ij}$ + 2.2 nm distance cutoff for tRNA-protein pairs resulted in a narrower distribution but with a peak at the same number of molecules, whereas shorter cutoffs gave significantly smaller clusters. The broader variation in cluster sizes from the geometrical clustering reflects in part a lack of robustness in estimating optimal cluster sizes from scale-free hierarchical clustering (17), and this is also the reason for why we used the contact-based criterion here instead of hierarchical geometrical clustering for determining tRNA and RP clusters.

**Diffusion analysis.** Translational diffusion ($D_{tr}$) was calculated for each molecule in the cytoplasmic system from the mean square displacement (MSD) of molecules between time $t$ and ($t$+$\tau$) for a given lag time $\tau$. Diffusion coefficients were then obtained from linear fits to MSD(τ) vs. τ (see Fig. S6).

$$D_{tr} = \frac{\mathrm{MSD}(\tau)}{6\tau} \qquad \text{M-5}$$

The first and last 1 μs of the cytoplasmic simulations were resampled so that conformations could be saved with a 1-ns interval. This allowed the analysis of all molecules in the dispersed and condensed states at the beginning and end of the trajectory and a comparison with previously published diffusion rates of macromolecules in the same system simulated in atomistic detail during similar time scales (13). In this case, the slope of MSD($\tau$) was fitted up until $\tau$ = 20 ns. Diffusion coefficients were calculated separately for molecules inside the tRNA and RP condensates as well as for molecules in the dilute phase. Molecules were considered to be part of a condensate if they remained part of the condensate during the entire lag time $\tau$.

For the five-component model system, diffusion was analyzed based on the last 100 μs of the simulations based on snapshots saved with a 100-ns interval and determining the slope of MSD($\tau$) up until $\tau$ = 2 μs.

**Phase separation analysis.** In order to determine critical temperatures, CG simulations were performed at temperatures ranging from 300 to 500 K in 10 K increments using the Langevin thermostat. The critical temperatures and concentration were obtained by fitting the temperature to the coexisting volume fractions using the following formulas (18):

$$\phi_H - \phi_L = A(T_c - T)^{0.32} \qquad \text{M-6}$$

$$\frac{1}{2}(\phi_H + \phi_L) = \phi_C + B(T - T_c) \qquad \text{M-7}$$

where $\varphi_H$ and $\varphi_L$ are the volume fractions of tRNA inside and outside of the clusters respectively, $T$ is the temperature, $T_c$ is the critical temperature and $\varphi_c$ is the critical volume fraction. This calculation was done for the model system simulations at different RP and $POS_L$ concentrations (see SI Sheet 2).

**Dynamic light scattering analysis.** The central observable of dynamic light scattering (DLS) experiments are time-dependent scattering intensity correlation functions $g_2(\tau)$ that are related to electric field correlation functions $g_1(\tau)$ according to:

$$g_2(\tau) - 1 = g_1(\tau)^2 \quad \text{M-8}$$

In case of a monodisperse solution with particles of a diameter *d*, a single exponential decay is observed with:

$$g_1(\tau; d) = e^{-2q^2 D(d)\tau} \quad \text{M-9}$$

with the wave vector

$$q = \frac{4\pi n}{\lambda} \sin\left(\frac{\theta}{2}\right) \quad \text{M-10}$$

and the diffusion according to Stokes-Einstein:

$$D(d) = \frac{k_B T}{6\pi\eta d} \quad \text{M-11}$$

where $n$ is the refractive index of the solvent medium (*i.e.* 1.335), $\lambda$ is the wavelength of the incident laser light (*i.e.* 633 nm), $\theta$ is the scattering angle (*i.e.* 173°), $k_B$ is the Boltzmann constant, $T$ is the temperature (*i.e.* 298 K), and $\eta$ is the viscosity of the solvent (*i.e.* 0.8882 cP).

The samples we considered were clearly polydisperse, requiring the fit of multiple exponential decays. Moreover, from previous studies and simulations we expect that at the smallest particle sizes there is an exponential decay of particle sizes due to dynamic cluster formation in the dilute phase (16, 19). Therefore, we fit the experimental data (*i.e.* $g_2(\tau) - 1$ ) to the following function:

$$g_2(\tau) - 1 = g_1(\tau)^2 \approx \sum_{i=1}^{10} a_c^2 e^{-\frac{2i}{t_c}} g_1^2(\tau; d_c) + \sum_{i=1}^{4} a_i^2 g_1^2(\tau; d_i) \quad \text{M-12}$$

Consequently, the parameters of the numerical fits were the size of the smallest particle, $d_c$, its contribution, $a_c$, decreasing according to the decay 'time' $t_c$, and an additional up to four discrete sizes $d_i$ with contributions $a_i$.

Using gnuplot, version 5.2, we fit the function according to M-12 to individual correlation functions as well to an average that was obtained after normalizing individual functions.

### Supplemental Theory

**Prediction of condensation between RNA and proteins.** An analytical model was constructed to reproduce the phase behavior seen in the simulations and allow a wider range of parameters to be explored. The analysis focuses on a two-component system consisting of a mixture of negatively charged particles *R*, equivalent to the RNA in the simulations, and particles *P*, equivalent to proteins, typically with a positive charge. The particles have charges $q_R$, $q_P$ and radii $r_R$, $r_P$. We consider a system of volume V in which R and P particles are present in total concentrations of $c_R$ and $c_P$. However, we do not include any finite-size effects and therefore the following analysis is scale-independent.

We assume that a phase-separated state is formed with a high-density condensate of volume $V_c$ and a low-density dilute phase of volume $V_d = V\text{-}V_c$, *i.e.* there is no change in the total system volume upon phase separation. The concentrations of R and P particles in the dilute and condensed phases are denoted as $c_{R,d}$, $c_{P,d}$, $c_{R,c}$, and $c_{P,c}$. From the concentrations, number densities $\rho_{R,d}$, $\rho_{P,d}$, $\rho_{R,c}$, and $\rho_{P,c}$ for R and P particles in the dilute and condensed phases are obtained according to $\rho = \frac{c}{\text{mM}} \cdot \frac{N_A}{10^{27}\text{nm}^3}$.

Mass conservation requires that:

$$(V - V_c)\rho_{\frac{R}{P},d} + V_c\rho_{\frac{R}{P},c} = V\rho_{\frac{R}{P}} \qquad \text{T-1}$$

leaving $V_c$ and $\rho_{R,c}$, and $\rho_{P,c}$ as independent variables to be determined for a given system in case of phase separation.

In general, the following scenarios are possible:

1) A fully disperse system, where there is no high-density condensate, *i.e.* $V_c = 0$, $\rho_{R,d} = \rho_{R,}$, $\rho_{P,d} = \rho_P$, $\rho_{R,c} = 0$, and $\rho_{P,c} = 0$;

2) a fully condensed system, *i.e.* $\rho_{R,d} = 0$, $\rho_{P,d} = 0$, $\rho_{R,c} = \rho_{R,}$ and $\rho_{P,c} = \rho_P$;

3) a phase-separated system with coexistence of dilute and condensed phases for both R and P particles, *i.e.* $\rho_{R,d} > 0$ and $\rho_{P,d} > 0$;

4) a phase-separated system where only R particles coexist between dilute and condensed phases, *i.e.* $\rho_{R,d} > 0$, $\rho_{P,d} = 0$, and $\rho_{P,c} = \rho_P$;

5) a phase-separated system where only P particles coexist between dilute and condensed phases, *i.e.* $\rho_{R,d} = 0$, $\rho_{P,d} > 0$, and $\rho_{R,c} = \rho_R$.

Which of these possible scenarios is assumed, depends on the total free energy of the system.

In order to determine the total free energy of the system, we begin by estimating the chemical potential for a particle either in the dilute and condensed phase from enthalpies and entropies according to:

$$\mu_{\frac{R}{P},\frac{d}{c}} = \Delta h_{\frac{R}{P},\frac{d}{c}} - T\Delta s_{\frac{R}{P},\frac{d}{c}} \qquad \text{T-2}$$

The enthalpy terms are decomposed into interactions of R-R, P-P, and R-P pairs:

$$\Delta h_{R,\frac{d}{c}} = \Delta h_{\frac{d}{c},RR} + \Delta h_{\frac{d}{c},RP} \qquad \text{T-3}$$

$$\Delta h_{P,\frac{d}{c}} = \Delta h_{\frac{d}{c},PR} + \Delta h_{\frac{d}{c},PP} \qquad \text{T-4}$$

Each pairwise interaction energy is estimated from the coarse-grained interaction potential by assuming a spherically symmetric distribution of particles but modulated as a function of distance according to radial distribution function extracted from simulations for each pair. This amounts to convoluting the pairwise interaction potential $U$ (see Eq. 3) with scaled volume- and density-normalized radial distribution functions $\hat{g}$ as follows:

$$\Delta h_{\frac{d}{c},RR} = \frac{1}{2}\rho_{R,\frac{d}{c}} \int_V \hat{g}_{RR,\frac{d}{c}}(r) U_{RR}(r) d^3r$$
$$= 2\pi\rho_{R,\frac{d}{c}} \int_0^{r_{max}} \hat{g}_{RR,\frac{d}{c}}(r) U_{RR}(r) r^2 dr \qquad \text{T-5}$$
$$\Delta h_{\frac{d}{c},PP} = 2\pi\rho_{P,\frac{d}{c}} \int_0^{r_{max}} \hat{g}_{PP,\frac{d}{c}}(r) U_{PP}(r) r^2 dr \qquad \text{T-6}$$
$$\Delta h_{\frac{d}{c},RP} = 2\pi\rho_{P,\frac{d}{c}} \int_0^{r_{max}} \hat{g}_{RP,\frac{d}{c}}(r) U_{RP}(r) r^2 dr \qquad \text{T-7}$$
$$\Delta h_{\frac{d}{c},PR} = 2\pi\rho_{R,\frac{d}{c}} \int_0^{r_{max}} \hat{g}_{PR,\frac{d}{c}}(r) U_{PR}(r) r^2 dr \qquad \text{T-8}$$

where the factor 1/2 corrects for double-counted self-interactions.

Different radial distribution functions were used for dilute and condensed environments (see Fig. S48). The *g(r)* functions extracted from the simulations were truncated at 20 nm and set to a constant value of 1 for larger radii to remove finite-size artifacts. Although the *g(r)* functions were determined from simulations with specific sizes $r_{R,MD}$, $r_{P,MD}$ of the R and P particles, other particle sizes could be considered by scaling the radial dependence of the *g(r)* functions according to the ratios $r_R/r_{R,MD}$, $r_P/r_{P,MD}$, and $(r_R+r_P)/(r_{R,MD}+r_{P,MD})$ for R-R, P-P, and R-P interactions. The upper integration limit $r_{max}$ was set to 100 nm for all interactions. At that radius and above, *U(r)* is negligible for the range of radii and charges considered here. With the fixed integration limit, the integrals in Eqs. T-5 to T-8 vary only with the charges and radii of particles R and P, and, thus, they are independent of particle concentrations. Then, the enthalpy contributions can be written as:

$$\Delta h_{\frac{d}{c},RR} = \rho_{R,\frac{d}{c}} x_{\frac{d}{c},RR} \qquad \text{T-9}$$
$$\Delta h_{\frac{d}{c},PP} = \rho_{P,\frac{d}{c}} x_{\frac{d}{c},PP} \qquad \text{T-10}$$
$$\Delta h_{\frac{d}{c},RP} = \rho_{P,\frac{d}{c}} x_{\frac{d}{c},RP} \qquad \text{T-11}$$
$$\Delta h_{\frac{d}{c},PR} = \rho_{R,\frac{d}{c}} x_{\frac{d}{c},PR} \qquad \text{T-12}$$

where the $x$ values represent the integrals in Eqs. T-5 to T-8 multiplied by 2π.

The entropy term was calculated based on the change of concentration in either dilute or condensed phases relative to the concentration in a fully disperse, non-separated system, which is the total system concentration, *i.e.* for the dilute phase:

$$\Delta s_{R,d} = R\log\left(\frac{c_R}{c_{R,d}}\right) = R\log\left(\frac{\rho_R}{\rho_{R,d}}\right) \qquad \text{T-13}$$
$$\Delta s_{P,d} = R\log\left(\frac{c_P}{c_{P,d}}\right) = R\log\left(\frac{\rho_P}{\rho_{P,d}}\right) \qquad \text{T-14}$$

where R is the universal gas constant. In estimating the entropy for the condensed phase, the finite volumes of the R and P particles were subtracted from the condensed phase volume $V_c$:

$$\Delta s_{R,c} = R\log\left(\frac{\rho_R}{\rho_{R,c}} \cdot \left(1 - \left(\rho_{R,c}V_R + \rho_{P,c}V_P\right)\right)\right) \qquad \text{T-15}$$
$$\Delta s_{P,c} = R\log\left(\frac{\rho_P}{\rho_{P,c}} \cdot \left(1 - \left(\rho_{R,c}V_R + \rho_{P,c}V_P\right)\right)\right) \qquad \text{T-16}$$

with the molecular volumes calculated from the radii of the spherical R and P particles:

$$V_{\frac{R}{P}} = \frac{4\pi}{3} r_{\frac{R}{P}}^3 \qquad \text{T-17}$$

Coexistence of the dilute and condensed phases assumes equilibrium, *i.e.*:

$$\mu_{R,d} = \mu_{R,c} \qquad \text{T-18}$$

$\mu_{P,d} = \mu_{P,c}$ T-19

In scenario 3), both, Eqs. T-18 and T-19, have to be satisfied simultaneously. For scenario 4), only Eq. T-18 needs to be satisfied under the condition that $\rho_{P,d} = 0$; and for scenario 5), only Eq. T-19 has to be satisfied with $\rho_{R,d} = 0$.

Solutions in terms of $\rho_{R,d}$, $\rho_{P,d}$, $\rho_{R,c}$, $\rho_{P,c}$, and $V_c$ were determined numerically by scanning $V_c$ and solving for the densities in the dilute phase (the densities in the condensed phase follow from Eq. T-1).

Eq. T-18 combined with Eqs. T-1, T-2, T-3, T-9, T-11, T-13, and T-15 gives the following:

$$0 = \mu_{R,d} - \mu_{R,c} \quad \text{T-20}$$

$$= \Delta h_{R,d} - T\Delta s_{R,d} - \Delta h_{R,c} + T\Delta s_{R,c}$$

$$= \rho_{R,d}x_{d,RR} + \rho_{P,d}x_{d,RP} - \rho_{R,c}x_{c,RR} - \rho_{P,c}x_{c,RP} + TR\log\left(\frac{\rho_{R,d}}{\rho_{R,c}} \cdot \left(1 - \left(\rho_{R,c}V_R + \rho_{P,c}V_P\right)\right)\right)$$

$$= \rho_{R,d}\left(x_{d,RR} + \frac{V - V_c}{V_c}x_{c,RR}\right) + \rho_{P,d}\left(x_{d,RP} + \frac{V - V_c}{V_c}x_{c,RP}\right) - \frac{V}{V_c}\left(\rho_R x_{c,RR} + \rho_P x_{c,RP}\right)$$

$$+TR\log\left(\frac{V_c\rho_{R,d} - V_R\rho_{R,d}\left(V\rho_R - (V-V_c)\rho_{R,d}\right) - V_P\rho_{R,d}\left(V\rho_P - (V-V_c)\rho_{P,d}\right)}{V\rho_R - (V-V_c)\rho_{R,d}}\right)$$

$$= f_R\left(\rho_{R,d}, \rho_{P,d}, V_c\right) \quad \text{T-21}$$

An analogous function $f_P\left(\rho_{R,d}, \rho_{P,d}, V_c\right)$ is obtained from Eq. T-19. There is no analytical solution, but $f_R\left(\rho_{R,d}, \rho_{P,d}, V_c\right) = 0$ and $f_P\left(\rho_{R,d}, \rho_{P,d}, V_c\right) = 0$ can be solved via the Newton-Raphson method given $V_c$ and either $\rho_{P,d}$ or $\rho_{R,d}$.

For scenario 4), $f_R\left(\rho_{R,d}, \rho_{P,d}, V_c\right) = 0$ was solved for different values of $V_c$ and $\rho_{P,d} = 0$; for scenario 5), $f_P\left(\rho_{R,d}, \rho_{P,d}, V_c\right) = 0$ was solved for values of $V_c$ and $\rho_{R,d} = 0$. For scenario 3), $\rho_{R,d}$ was scanned as well and the value of $\rho_{P,d}$ was determined for given values of $V_c$ and $\rho_{R,d}$ by first solving $f_R\left(\rho_{R,d}, \rho_{P,d}, V_c\right) = 0$. The resulting value of $\rho_{P,d}$ was then used with $V_c$ to solve $f_P\left(\rho_{R,d}, \rho_{P,d}, V_c\right) = 0$ for a refined value of $\rho_{R,d}$.

Mathematically possible solutions include cases where the volume fractions in the cluster exceed what is physically realistic and what would prevent liquid-like behavior inside the condensed state. In order to exclude such solutions, it was required that the combined macromolecular volume in the condensed phase is less than 30% of the total volume of the condensed phase, *i.e.*:

$$\rho_{R,c}V_R + \rho_{P,c}V_P < 0.3 \quad \text{T-22}$$

The total system energy is calculated according to:

$$\Delta G = \mu_{R,d} \cdot (V - V_c)\rho_{R,d} + \mu_{R,c} \cdot V_c\rho_{R,c} + \mu_{P,d} \cdot (V - V_c)\rho_{P,d} + \mu_{P,c} \cdot V_c\rho_{P,c} - TS_{mix} \quad \text{T-23}$$

where $S_{mix}$ is the overall mixing entropy according to the ratio of particles R and P in the dilute and condensed phases according to:

$$S_{mix} = S_{mix,d} + S_{mix,c} \quad \text{T-24}$$

$$S_{mix,d} = R(V - V_c)\left(\rho_{R,d}\log\frac{\rho_{R,d}}{\rho_{R,d}+\rho_{P,d}} + \rho_{P,d}\log\frac{\rho_{P,d}}{\rho_{R,d}+\rho_{P,d}}\right) \quad \text{T-25}$$

$$S_{mix,c} = RV_c\left(\rho_{R,c}\log\frac{\rho_{R,c}}{\rho_{R,c}+\rho_{P,c}} + \rho_{P,c}\log\frac{\rho_{P,c}}{\rho_{R,c}+\rho_{P,c}}\right) \quad \text{T-26}$$

For the five scenarios described above, total free energies were then calculated as follows:

1) <u>Disperse</u>:

$$\Delta G_1 = \mu_{R,disperse} \cdot V\rho_R + \mu_{P,disperse} \cdot V\rho_P - TRV\left(\rho_R \log\frac{\rho_R}{\rho_R+\rho_P} + \rho_P \log\frac{\rho_P}{\rho_R+\rho_P}\right) \qquad \text{T-27}$$

where $\mu_{\frac{R}{P},disperse}$ were calculated according to Eqs. T-2 to T-8 using RDFs from the disperse phase extracted from our molecular dynamics simulations before condensates started to form.

2) Condensed:

$$\Delta G_2 = \mu_{R,c} \cdot V\rho_R + \mu_{P,c} \cdot V\rho_P - TRV_c\left(\rho_R \log\frac{\rho_R}{\rho_R+\rho_P} + \rho_P \log\frac{\rho_P}{\rho_R+\rho_P}\right) \qquad \text{T-28}$$

3) R and P in phase coexistence:

$$\Delta G_3 = \mu_{R,c} \cdot V\rho_R + \mu_{P,c} \cdot V\rho_P \qquad \text{T-29}$$

$$-TR(V - V_c)\left(\rho_{R,d} \log\frac{\rho_{R,d}}{\rho_{R,d} + \rho_{P,d}} + \rho_{P,d} \log\frac{\rho_{P,d}}{\rho_{R,d} + \rho_{P,d}}\right)$$

$$-TRV_c\left(\rho_{R,c} \log\frac{\rho_{R,c}}{\rho_{R,c}+\rho_{P,c}} + \rho_{P,c} \log\frac{\rho_{P,c}}{\rho_{R,c}+\rho_{P,c}}\right)$$

since $\mu_{R,c} = \mu_{R,d}$ and $\mu_{P,c} = \mu_{P,d}$

4) R in phase coexistence, $\rho_{P,d} = 0$:

$$\Delta G_4 = \mu_{R,c} \cdot V\rho_R + \mu_{P,c} \cdot V\rho_P - TRV_c\left(\rho_{R,c} \log\frac{\rho_{R,c}}{\rho_{R,c}+\rho_{P,c}} + \rho_{P,c} \log\frac{\rho_{P,c}}{\rho_{R,c}+\rho_{P,c}}\right) \qquad \text{T-30}$$

5) P in phase coexistence, $\rho_{R,d} = 0$:

$$\Delta G_5 = \mu_{R,c} \cdot V\rho_R + \mu_{P,c} \cdot V\rho_P - TRV_c\left(\rho_{R,c} \log\frac{\rho_{R,c}}{\rho_{R,c}+\rho_{P,c}} + \rho_{P,c} \log\frac{\rho_{P,c}}{\rho_{R,c}+\rho_{P,c}}\right) \qquad \text{T-31}$$

The scenario with the overall lowest free energy was then considered to be the predicted state. A program implementing this model is available at http://github.com/feiglab/phasesep.

**Prediction of FRET efficiencies**

FRET efficiencies for mixtures of RNA and proteins at different concentrations were estimated from the predicted amount of RNA inside and outside the condensates as follows:

The theory described above predicts phase separation with the densities of RNA in the dilute and condensed phases given as $\rho_{R,d}$ and $\rho_{R,c}$. From the densities the concentration of RNA in the dilute ($[R_d]$) and condensed ($[R_c]$) phases with respect to the total volume is obtained as follows:

$$[R_d] = \rho_{R,d} \cdot \frac{V-V_c}{V} \qquad \text{T-32}$$

$$[R_c] = \rho_{R,c} \cdot \frac{V_c}{V} \qquad \text{T-33}$$

A fraction of RNA is labeled with fluorophores. The total concentration of labeled RNA is denoted as [F]; the concentration in the dilute and condensed phases, again with respect to the total system volume, is denoted as $[F_d]$ and $[F_c]$, respectively. Then:

$$[F] = [F_c] + [F_d] \qquad \text{T-34}$$

and

$$[R_d] = [U_d] + [F_d] \qquad \text{T-35}$$

$$[R_c] = [U_c] + [F_c] \qquad \text{T-36}$$

where $[U_d]$ and $[U_c]$ are the concentrations of unlabeled RNA in the dilute and condensed phases.

We further make an assumption that there is an equilibrium of labeled RNA to exchange between the dilute and condensed phases while maintaining the overall ratio of RNA between the two phases:

$$[F_d] + [U_c] \leftrightarrow [F_c] + [U_d] \quad \text{T-37}$$

with the equilibrium constant $K$ given as:

$$K = \frac{[F_c][U_d]}{[F_d][U_c]} \quad \text{T-38}$$

Because of the hydrophobic character of the FRET labels we expect that labeled RNA has an affinity for the less-hydrated condensate, *i.e.* $K > 1$.

Equations T-34, T-35, T-36, and T-38 can be solved for $[F_c]$ as a function of $[R_d]$, $[R_c]$, $[F]$, and $K$ to give the fraction of labeled RNA in the condensate as:

$$f = \frac{[F_c]}{[F]} \quad \text{T-39}$$

Based on the resulting value of $f$, FRET efficiencies $E$ were then estimated according to:

$$E = E_0(1 - f) + E_c f \quad \text{T-40}$$

where $E_0$ and $E_c$ are the FRET efficiencies at zero protein concentration and in the condensed phase, respectively. $E_0$ was taken from experiment and $E_c$ was estimated by convoluting the distribution of minimum RNA-RNA distances in the condensed phase extracted from the simulations with $1/(1+(r/r_0)^6)$, where $r$ is the distance between RNA molecules and $r_0$ is a constant that depends on the fluorescence label and additional factors such as the anisotropy of the orientational sampling and the index of diffraction of the medium.

We applied this formalism to interpret the FRET experiments on trypsin based on predicted RNA fractions in the condensed phase (Fig. S40) using the minimum distance distribution of RNA shown in Fig. S49. We took $E_0$ = 0.24 from experiment and found good agreement between experiment and theory for $r_0$ = 4.10 nm and $K$ = 100 (Fig. 7). We note that the value $r_0$ = 4.10 nm is lower than typical values assumed for the Cy3-Cy5 pair (20), but the condensed state differs from typical solution conditions, whereas the spherical models used here allow only very approximate estimates of the true donor-acceptor distances and neglect orientational dependence in fluorescent energy transfer (21).

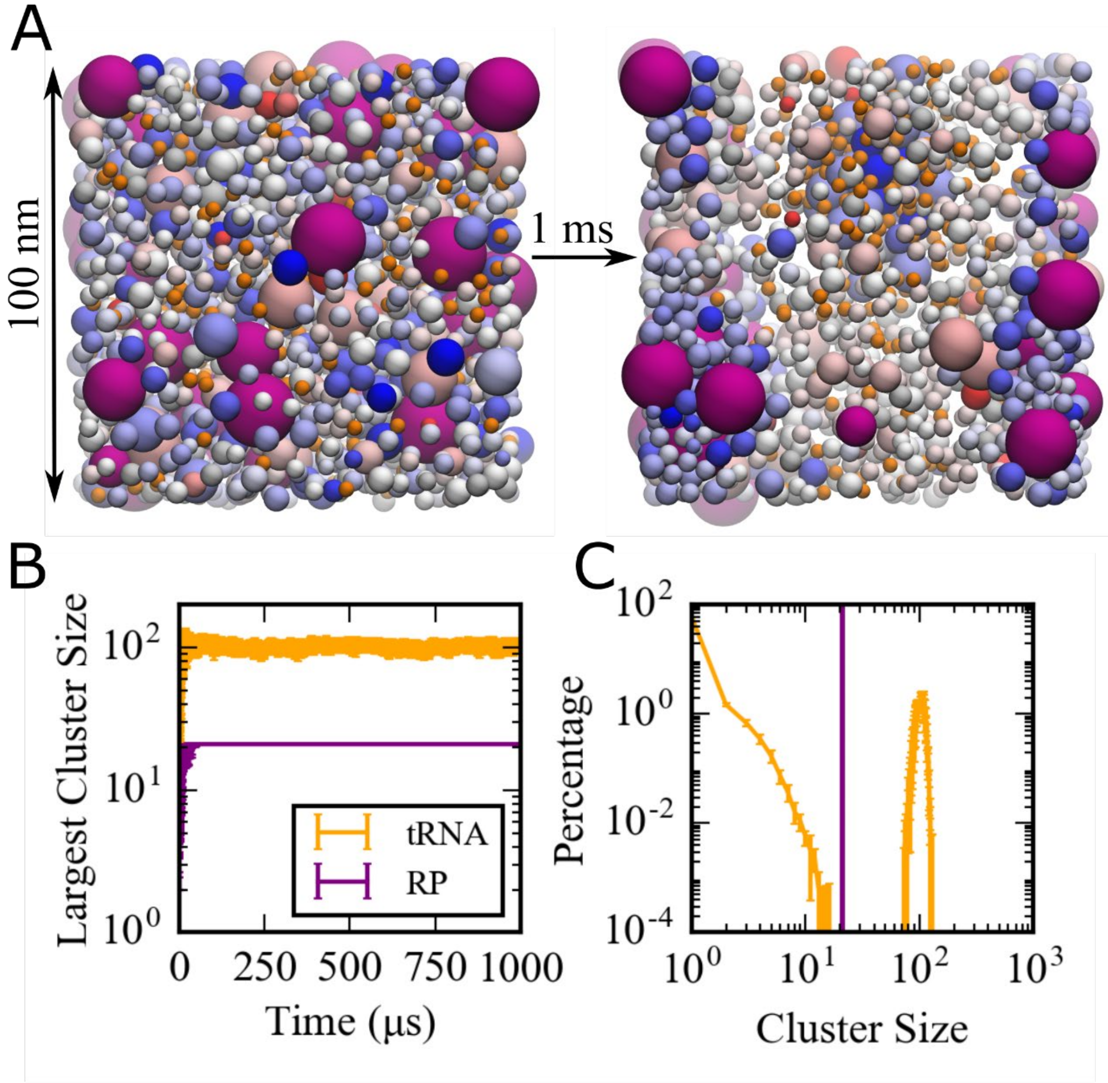


**Fig. S1.** (A) Coarse-grained simulations of a model bacterial cytoplasm with an alternative effective charge model using Eq. M-3. Initial and final frames for a 1 ms simulation of a 100 nm system are shown with tRNAs in orange, ribosomes in magenta, and other molecules colored according to their charges (blue towards positive charges; red towards negative charges). Sphere sizes are shown proportional to molecular sizes. Large pink spheres correspond to GroEL particles. (B) Size of the largest cluster vs. simulation time in 100 nm system. (C) Cluster size distributions for tRNA and RP during the last 500 μs.

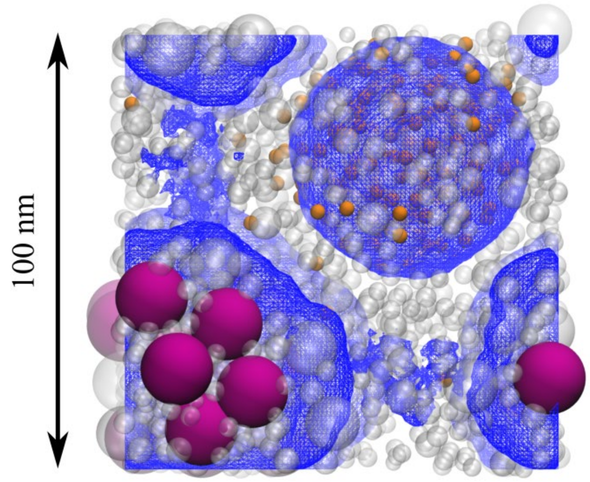


**Fig. S2.** Density variation in the cytoplasmic model system during the last 500 μs of the simulation. Grid-based contours at volume fractions exceeding 10% are indicated in blue and overlaid onto the final snapshot of the system after 1 ms. The density map was calculated using 10 nm voxel sizes and molecules were counted in a specific voxel if their volume based on their van der Waals radii was covered by that voxel.

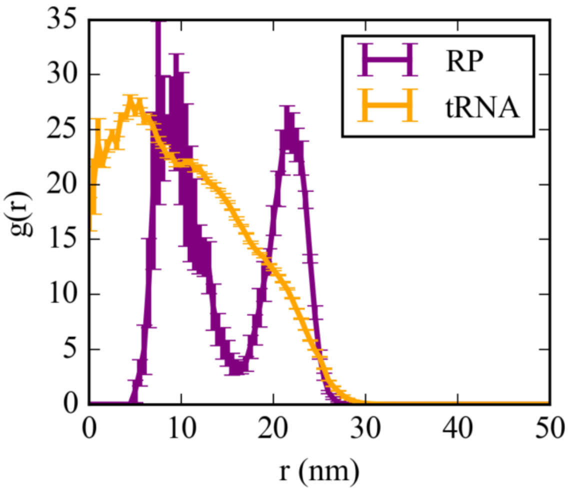


**Fig. S3.** Radial distribution curves for tRNA and RP in condensates from the center of their respective condensates.

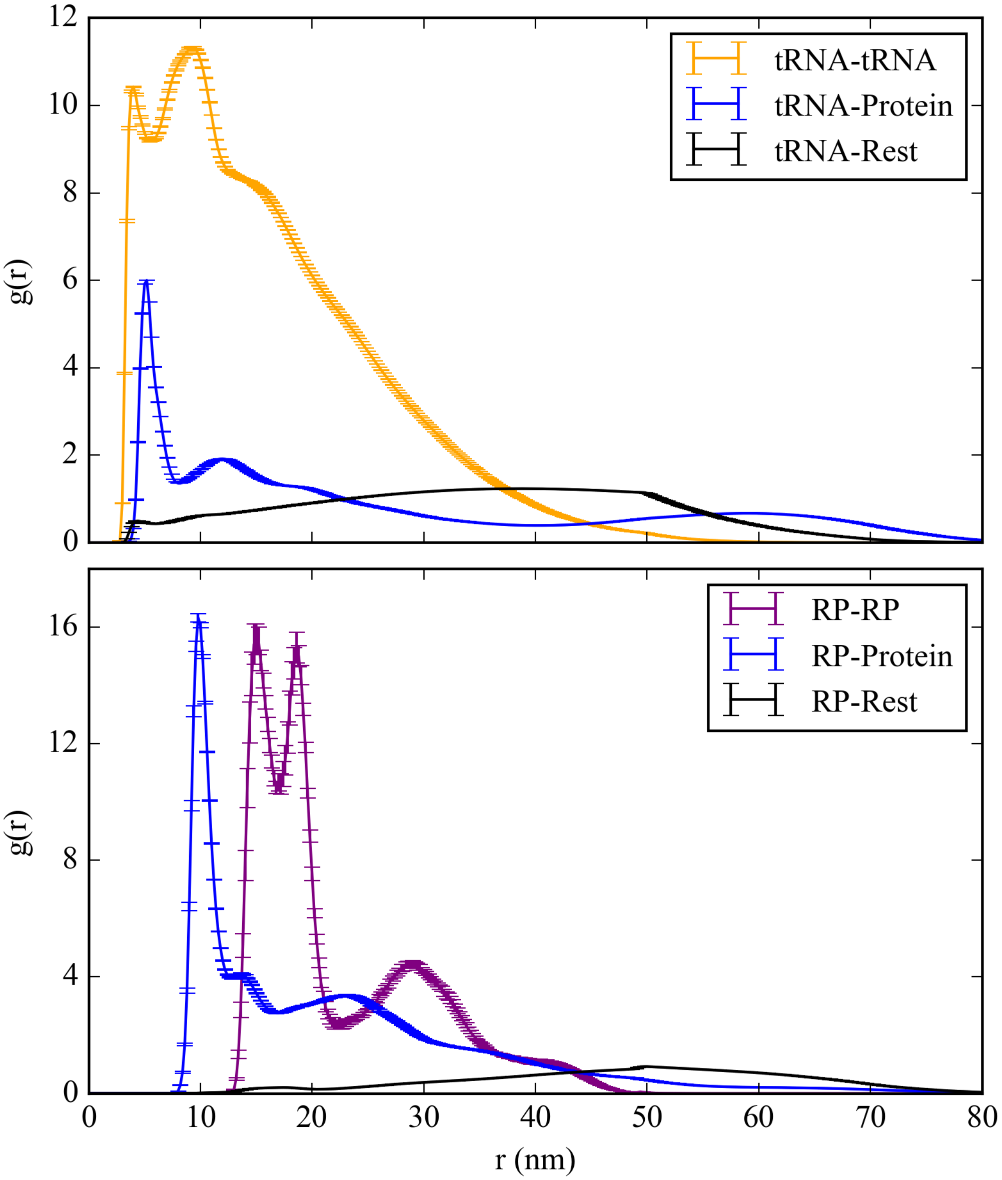


**Fig. S4.** Pairwise radial distribution functions between tRNA, RP, and positively charged protein particles and any other particles in the cytoplasmic model system.

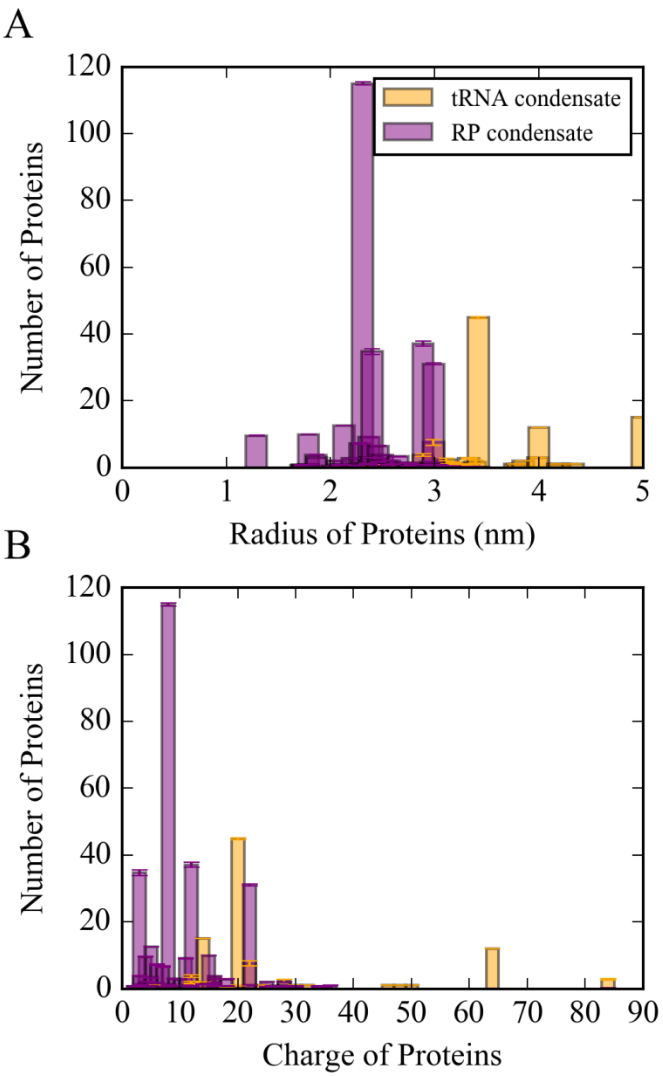


**Fig. S5.** Number of proteins in the tRNA and RP condensates vs. the radius (A) and charge (B) of the proteins found in the condensates. A 2.2 nm distance cutoff was used to identify molecules as part of the condensates (see SI Methods)

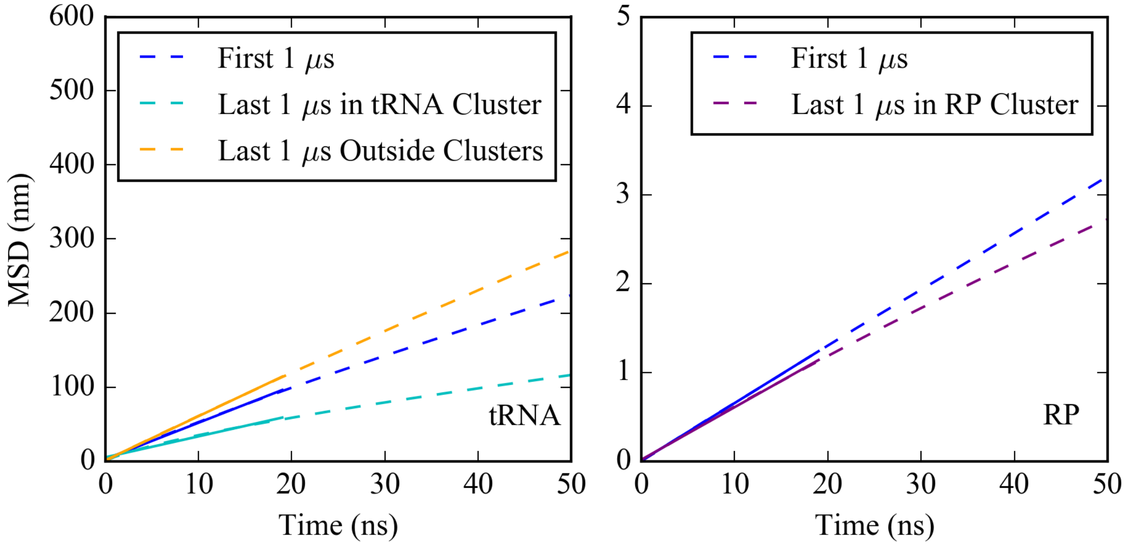


**Fig. S6.** Mean square displacement (MSD) for tRNA (left) and RP (right) particles during the first and last 1 µs of the cytoplasmic simulations.

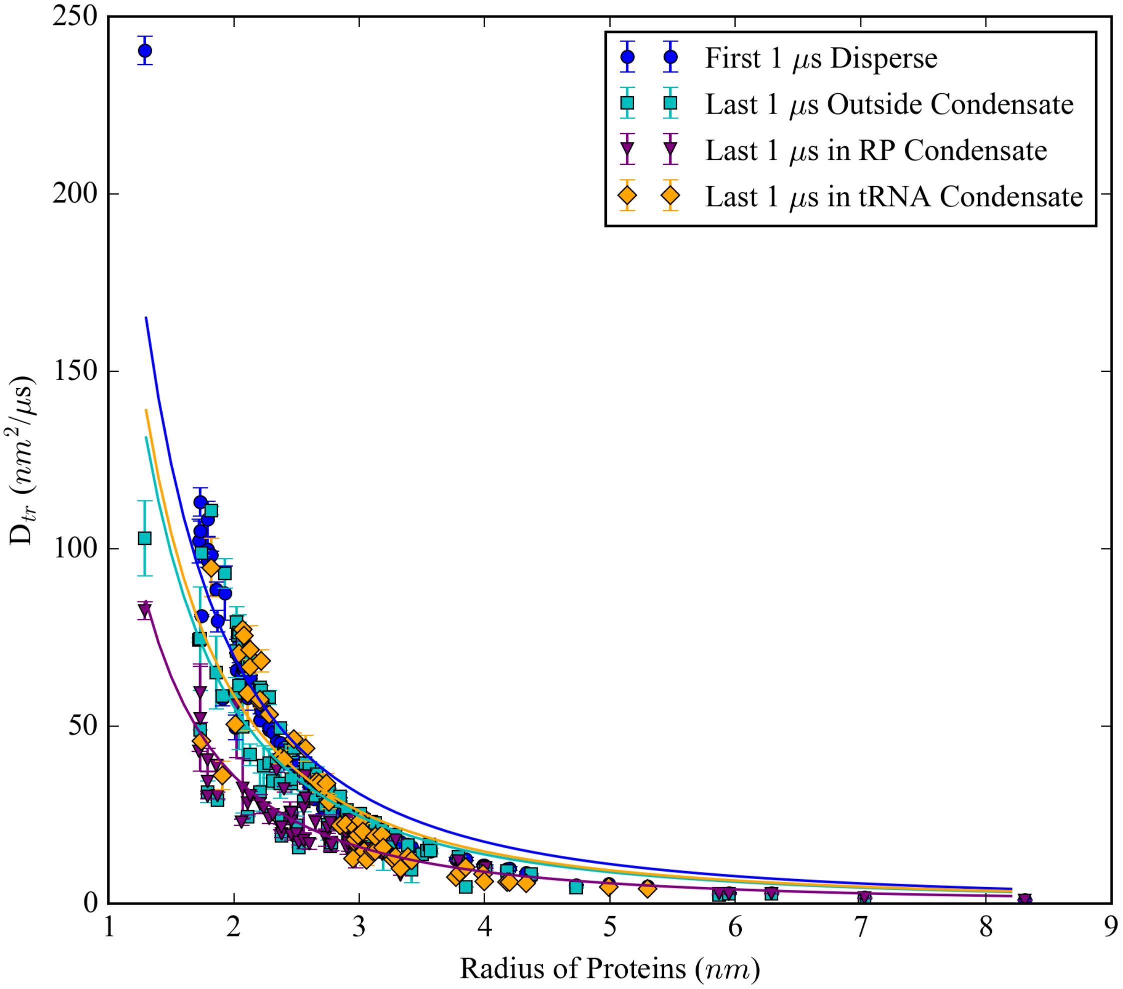


**Fig. S7.** Translational diffusion of macromolecules in the cytoplasmic system as a function of the radius of the macromolecules during the first and last 1 μs of the simulations. For the last 1 μs the diffusion coefficients were calculated separately for molecules inside and outside the tRNA and RP condensates. Solid lines depict fitting functions as a function of the particle radius for the dispersed system ($D_{tr}$= $279/r^2$), outside of condensates ($D_{tr}$= $222/r^2$), inside tRNA condensates ($D_{tr}$ = $235/r^2$), and inside RP condensates ($D_{tr}$ = $144/r^2$).

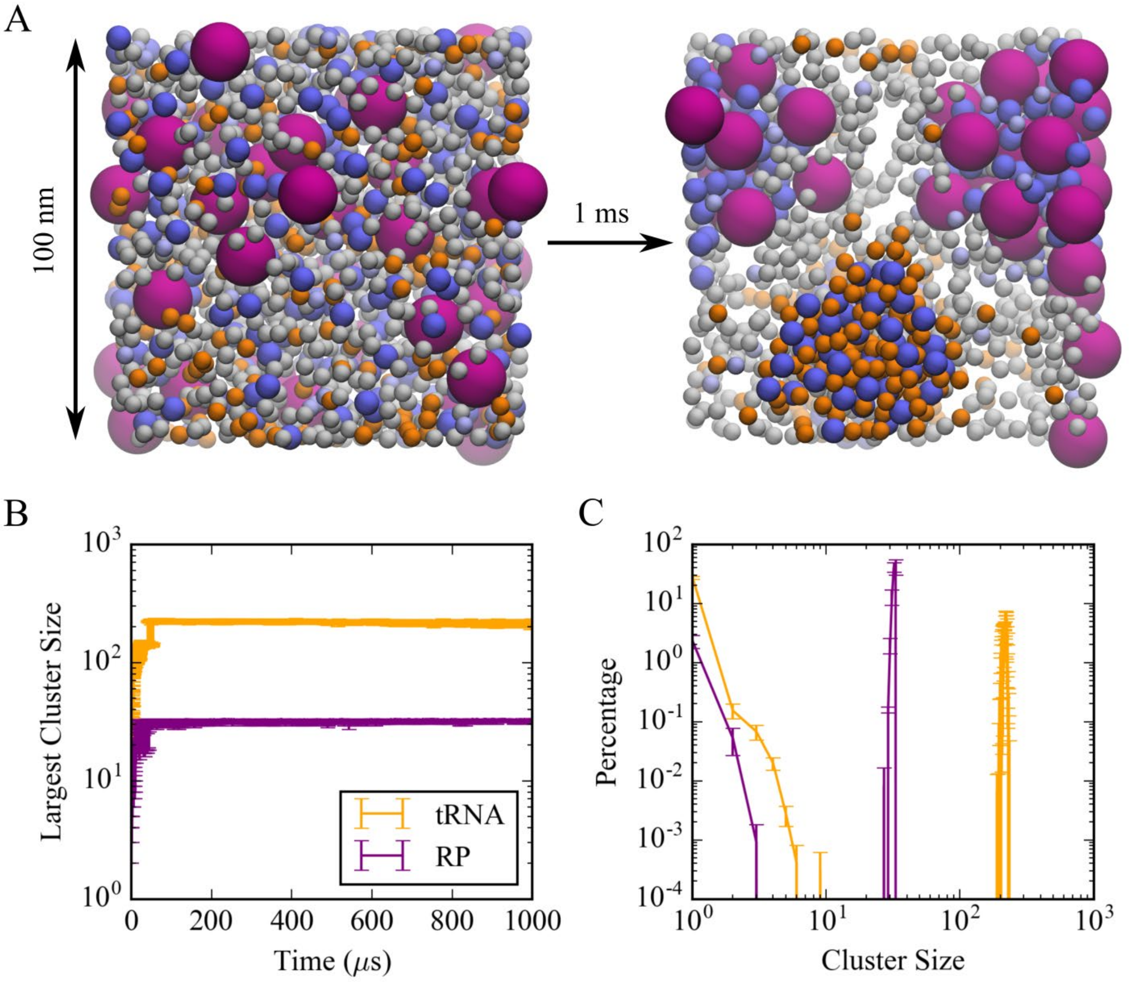


**Fig. S8.** Initial and final frames of the five-component model system simulation (A); time evolution of cluster formation for tRNA and RP clusters (B); and cluster size distributions (C).

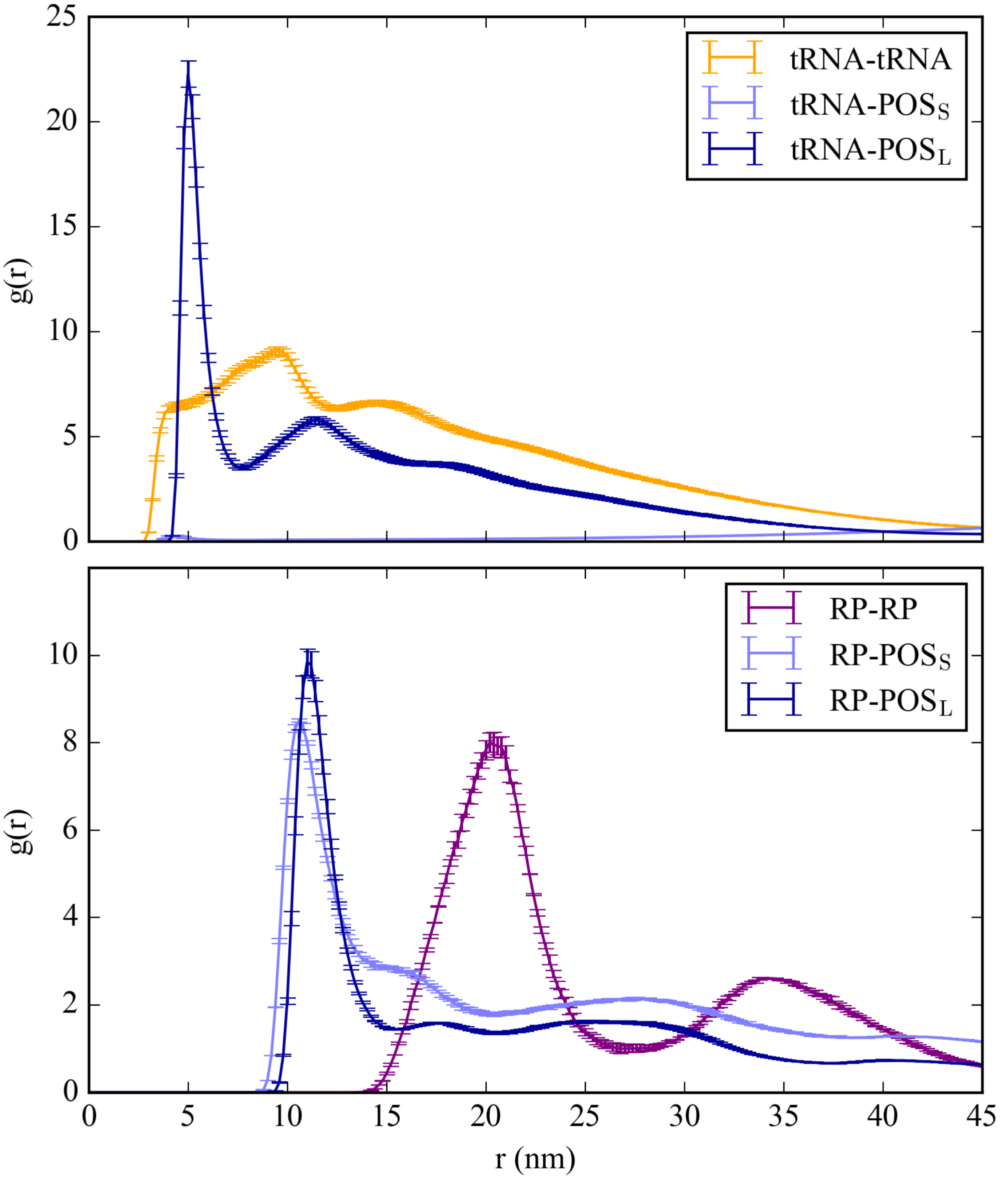


**Fig. S9.** Radial distribution functions for interactions between different particle types in the five-component model.

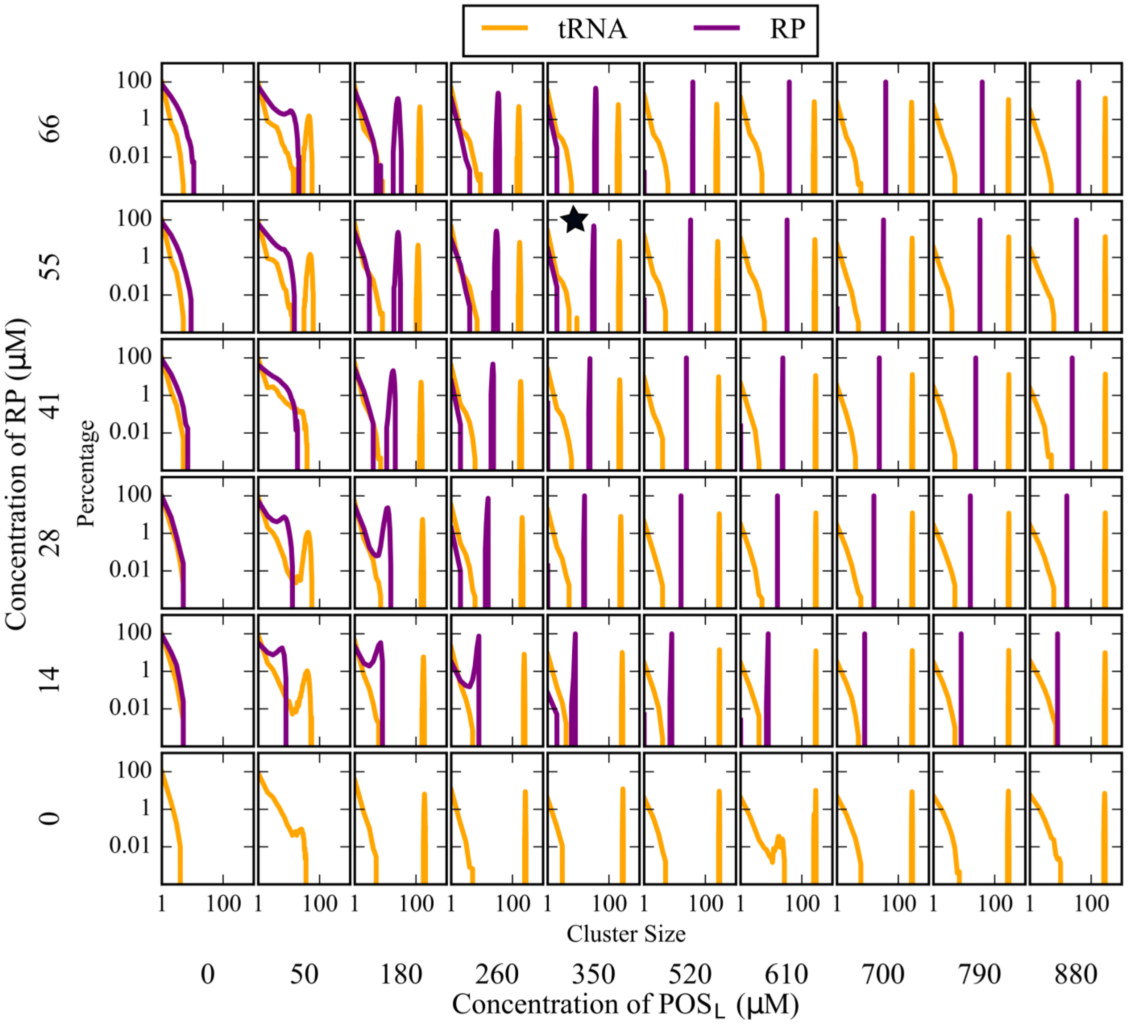


**Fig. S10.** Cluster size distribution of tRNA and RP as a function of [RP] and [$POS_L$] in the five-component model system. The black star indicates the conditions that match the full cytoplasmic model.

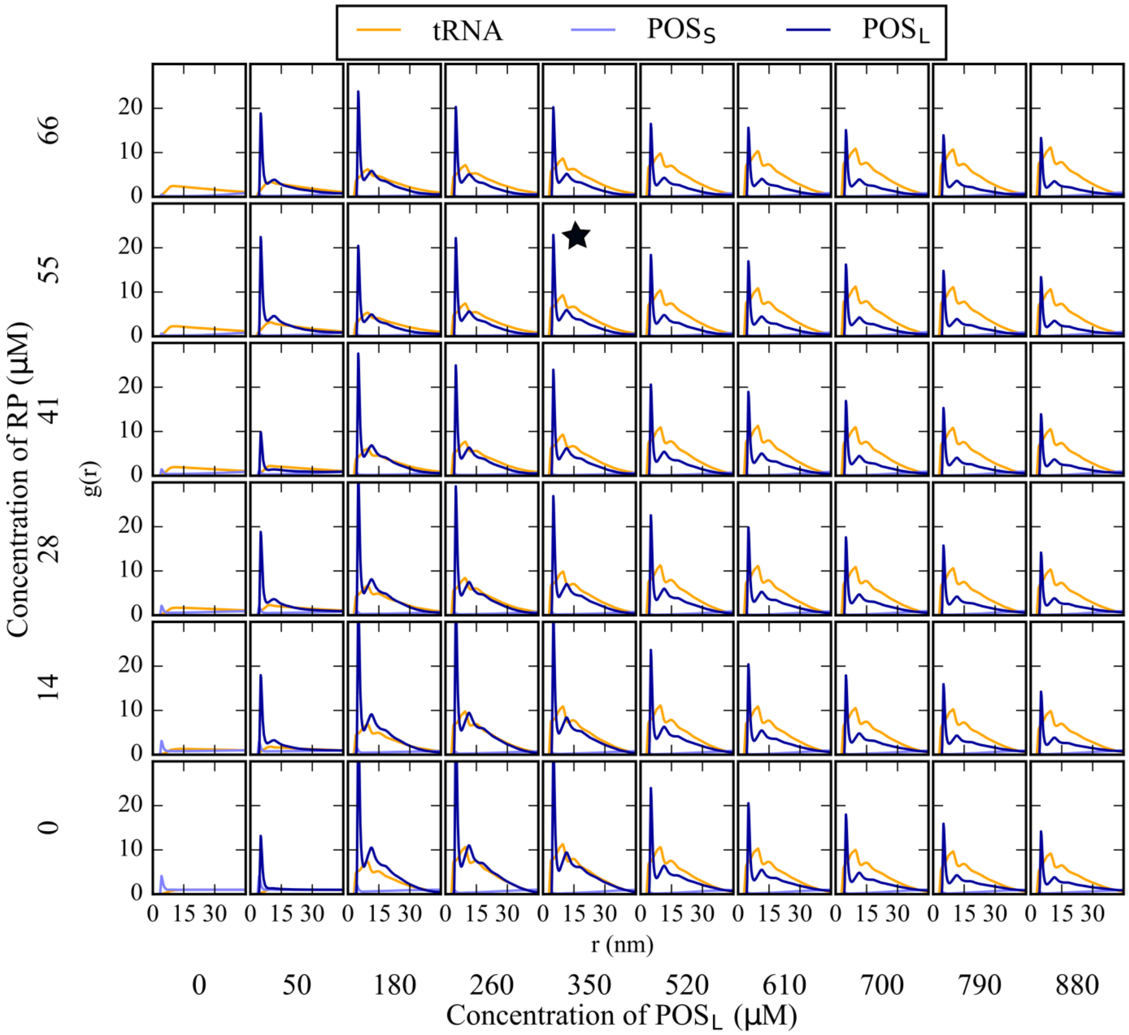


**Fig. S11.** Radial distribution functions for tRNA with tRNA, POS$_S$, and POS$_L$ as a function of [RP] and [POS$_L$] in the five-component model system. The black star indicates the conditions that match the full cytoplasmic model.

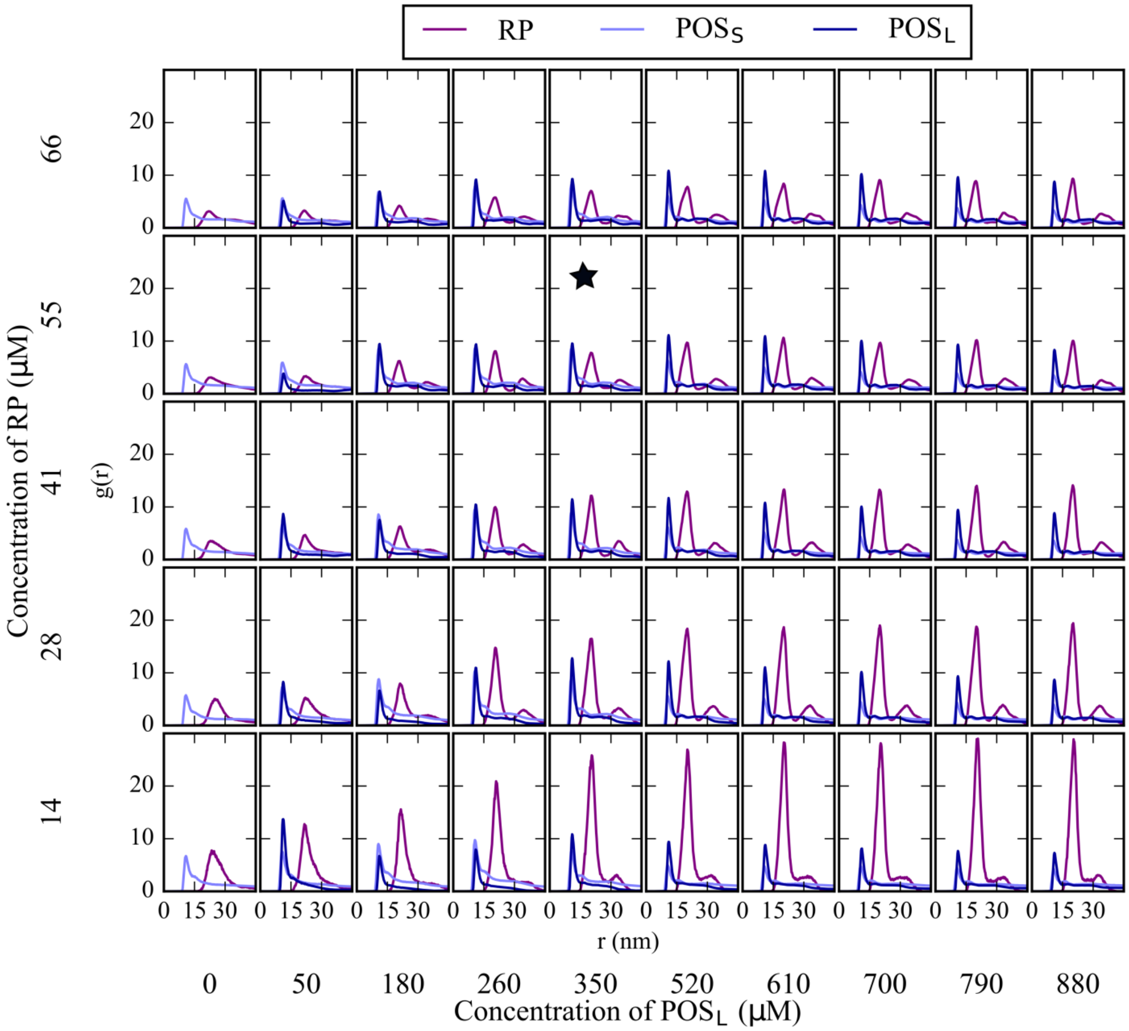


**Fig. S12.** Radial distribution functions for RP with RP, $POS_S$, and $POS_L$ as a function of [RP] and [$POS_L$] concentration in the five-component model system. The black star indicates the conditions that match the full cytoplasmic model.

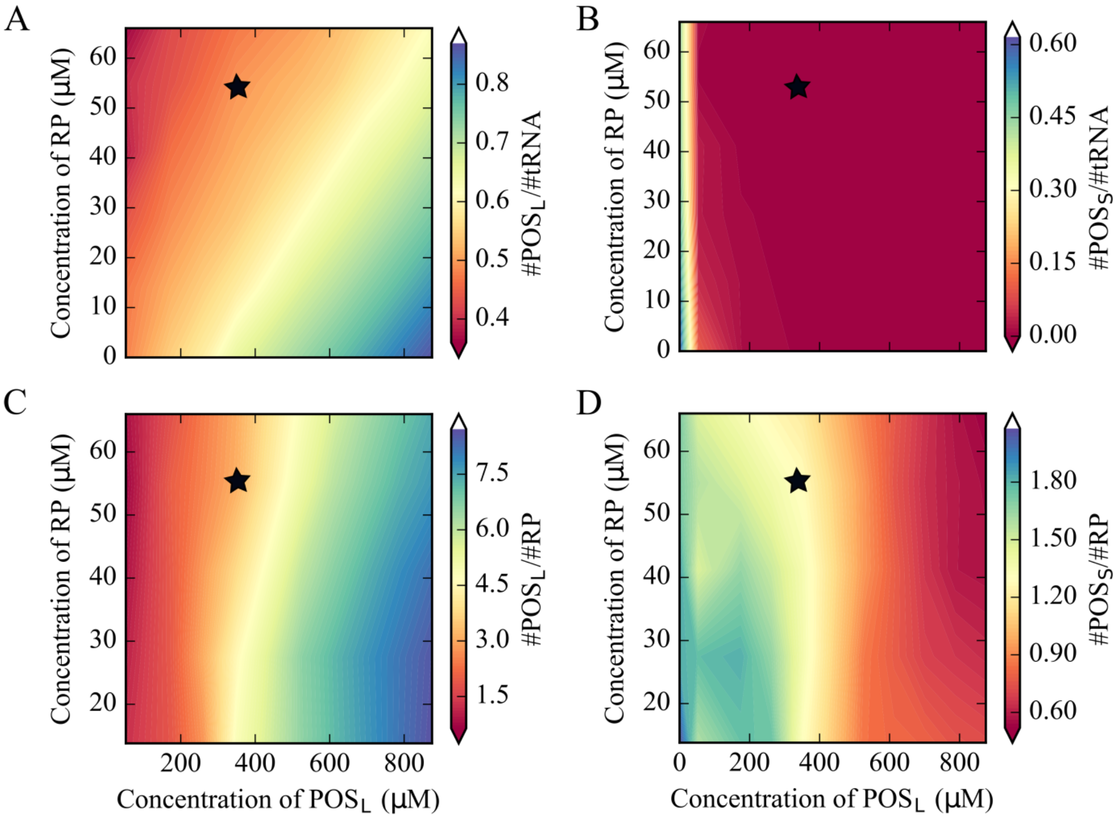


**Fig. S13.** Relative abundance of $POS_L$ and $POS_S$ in the largest tRNA and RP clusters with the five-component model as a function of [RP] and [$POS_L$]: (A) Ratio of $POS_L$ vs. tRNA; (B) ratio of $POS_S$ vs. tRNA; (C) ratio of $POS_L$ vs. RP; (D) ratio of $POS_S$ vs. RP. The black star indicates the conditions that match the full cytoplasmic model.

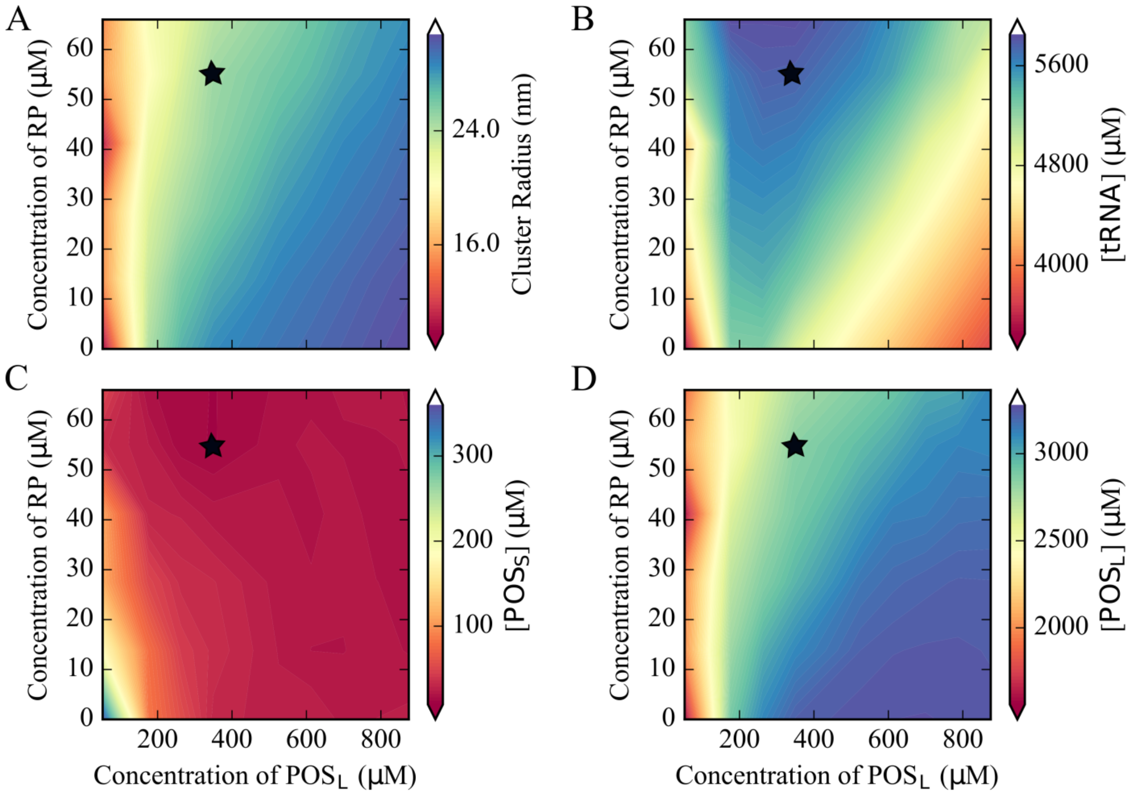


**Fig. S14.** Volume-equivalent radii for largest cluster in tRNA condensates with five-component model (A); macromolecular concentrations inside tRNA condensates for tRNA (B), $POS_S$ (C) and $POS_L$ (D). The black star indicates the conditions that match the full cytoplasmic model.

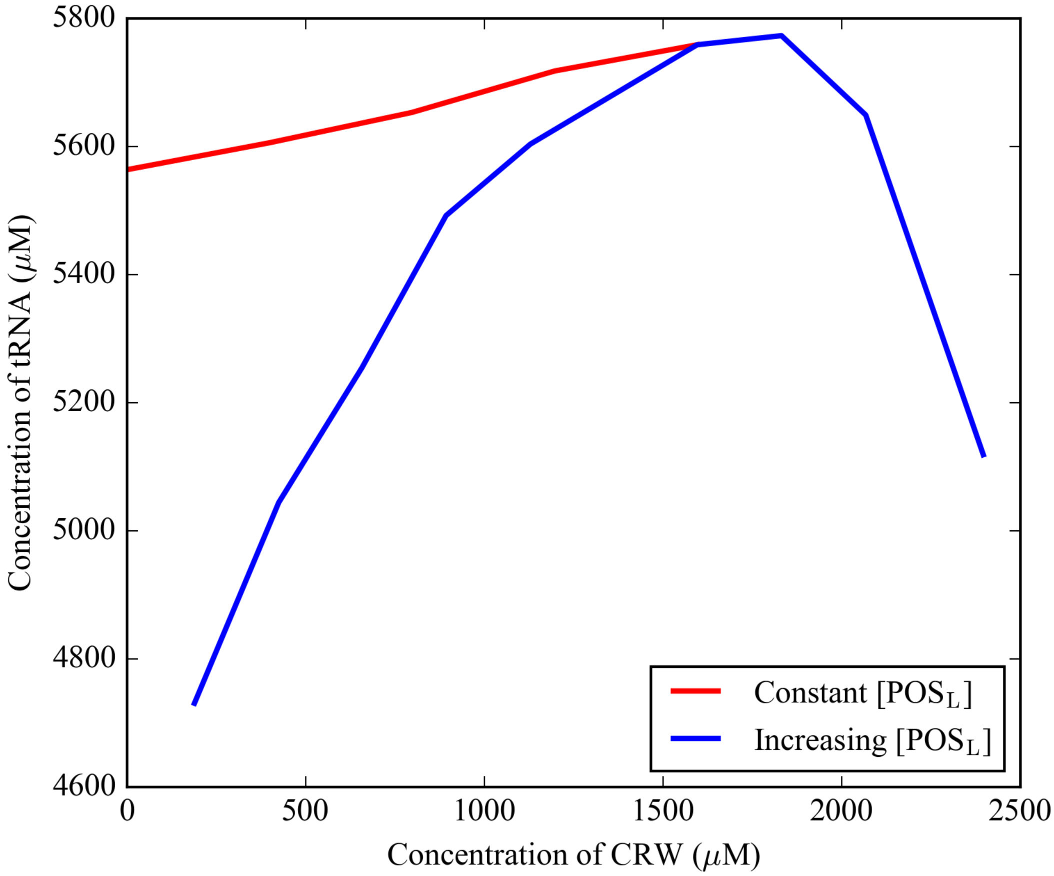


**Fig. S15.** Concentration of tRNA inside the tRNA condensates as a function of [CRW] at constant and increasing values of [$POS_L$] from simulations of the five-component model.

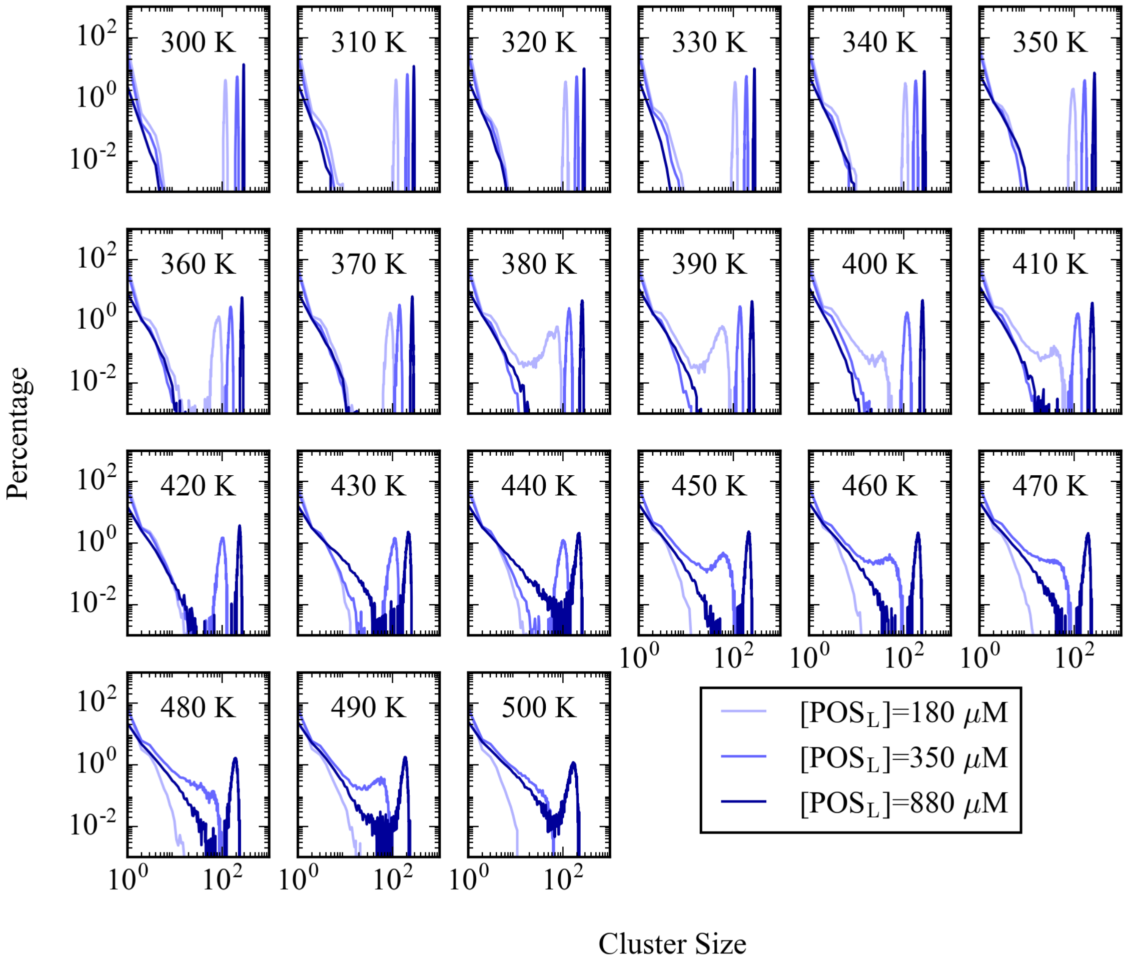


**Fig. S16.** Cluster size distributions of tRNA at [RP] = 55 μM and three $POS_L$ concentrations (see Legend) for temperatures between 300 and 500 K from simulations of the five-component model.

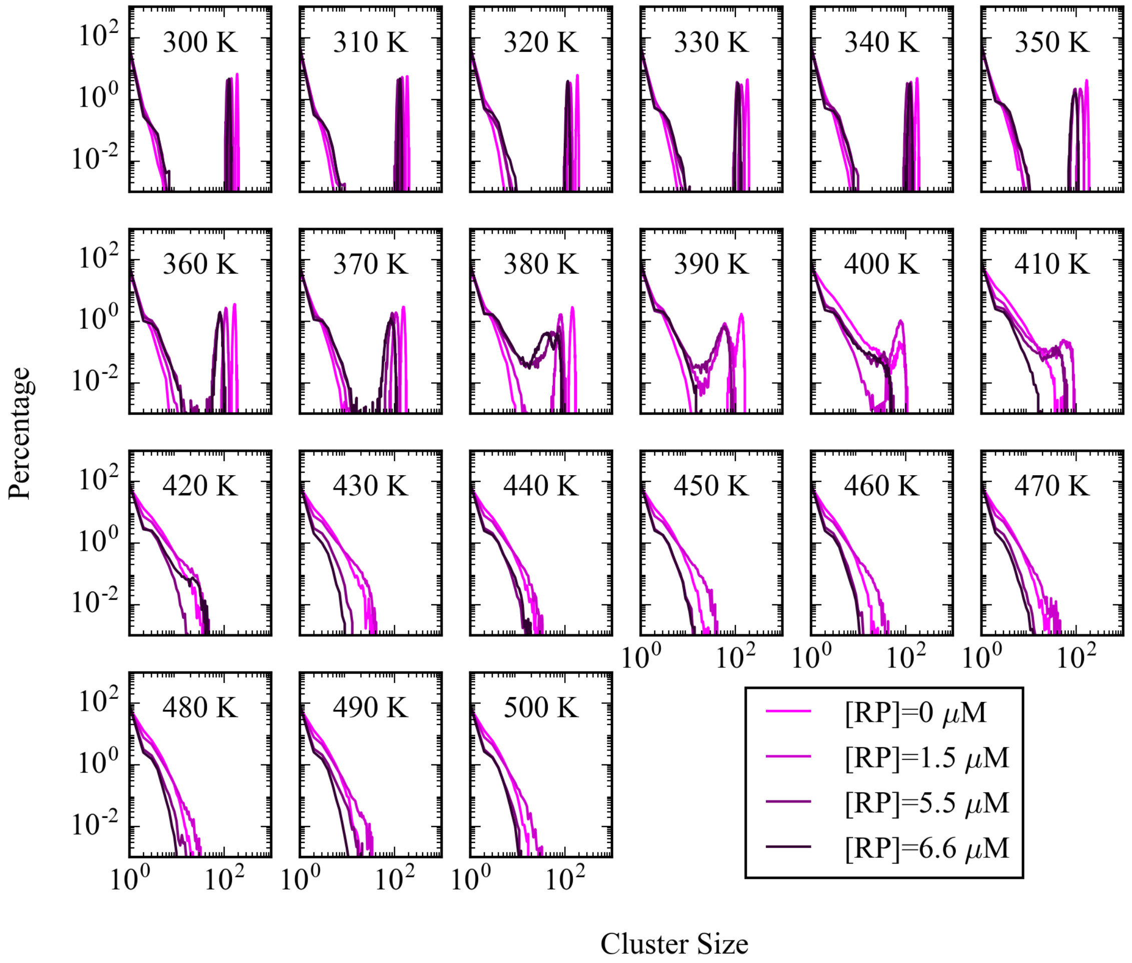


**Fig. S17.** Cluster size distributions of tRNA at [$POS_L$] = 180 μM and a range of RP concentrations (see Legend) for temperatures between 300 and 500 K from simulations of the five-component model.

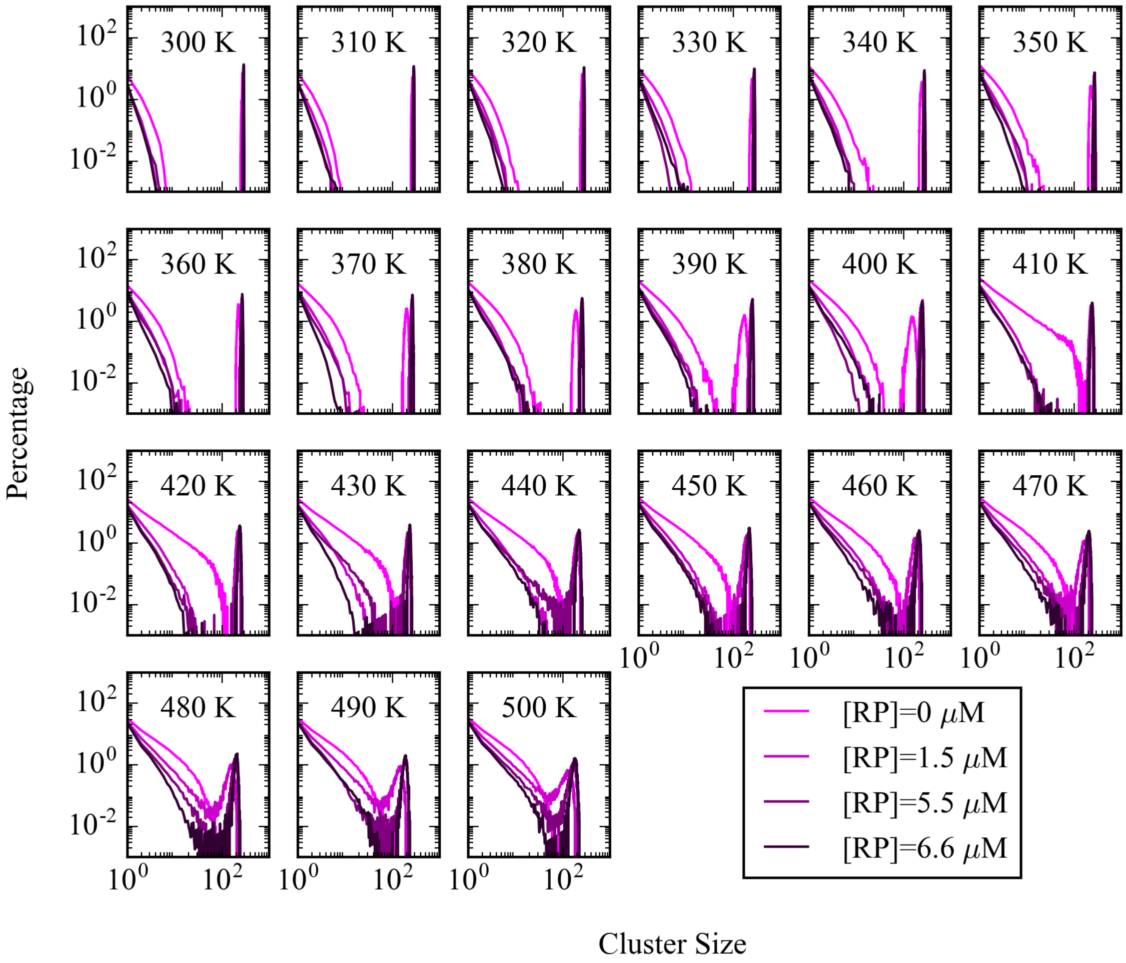


**Fig. S18.** Cluster size distributions of tRNA at [$POS_L$] = 880 μM and a range of RP concentrations (see Legend) for temperatures between 300 and 500 K from simulations of the five-component model.

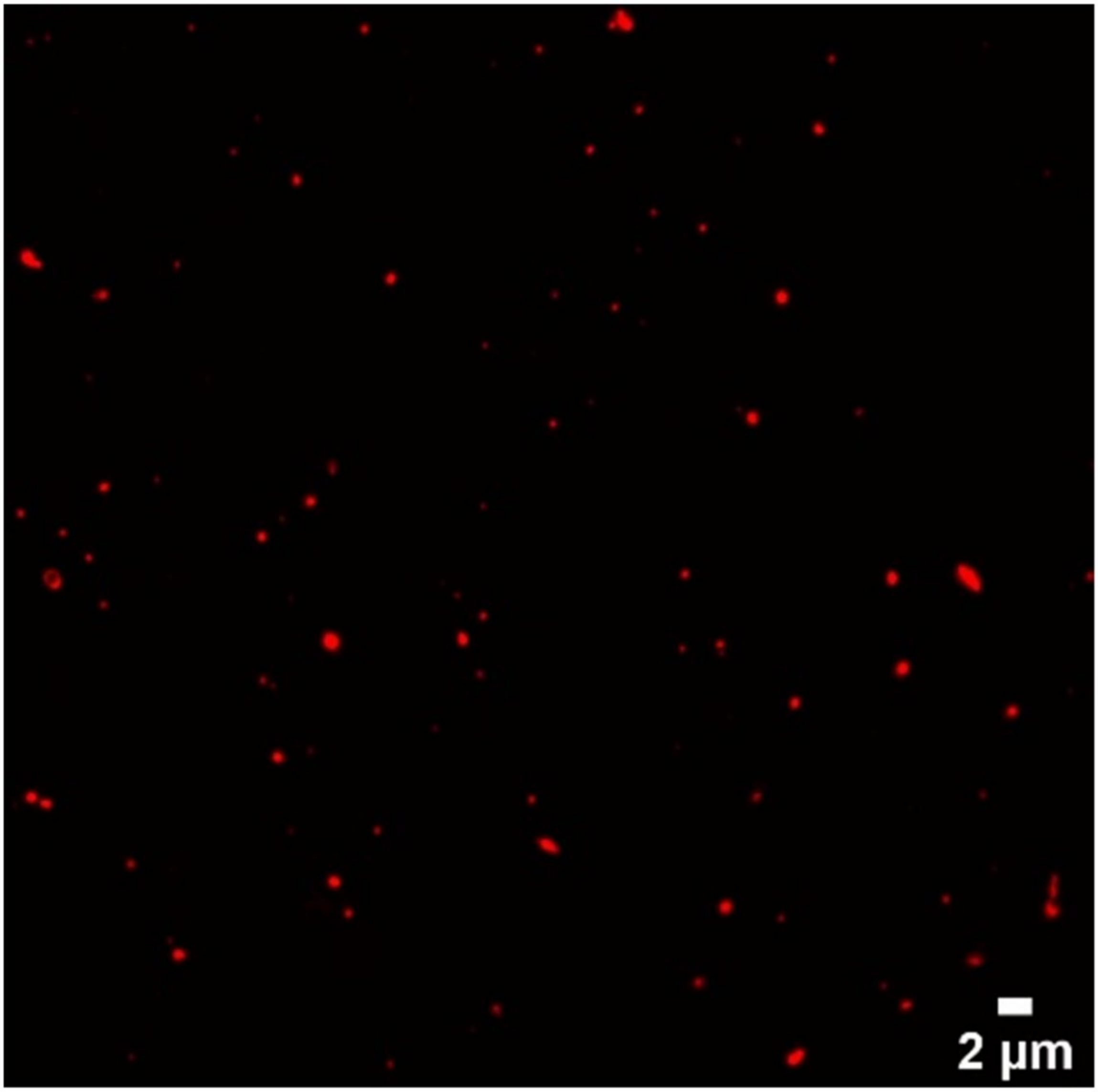


**Fig. S19.** Confocal microscopy of labeled J345 RNA for a mixture between J345 RNA at 0.45 mM and trypsin at 0.35 mM.

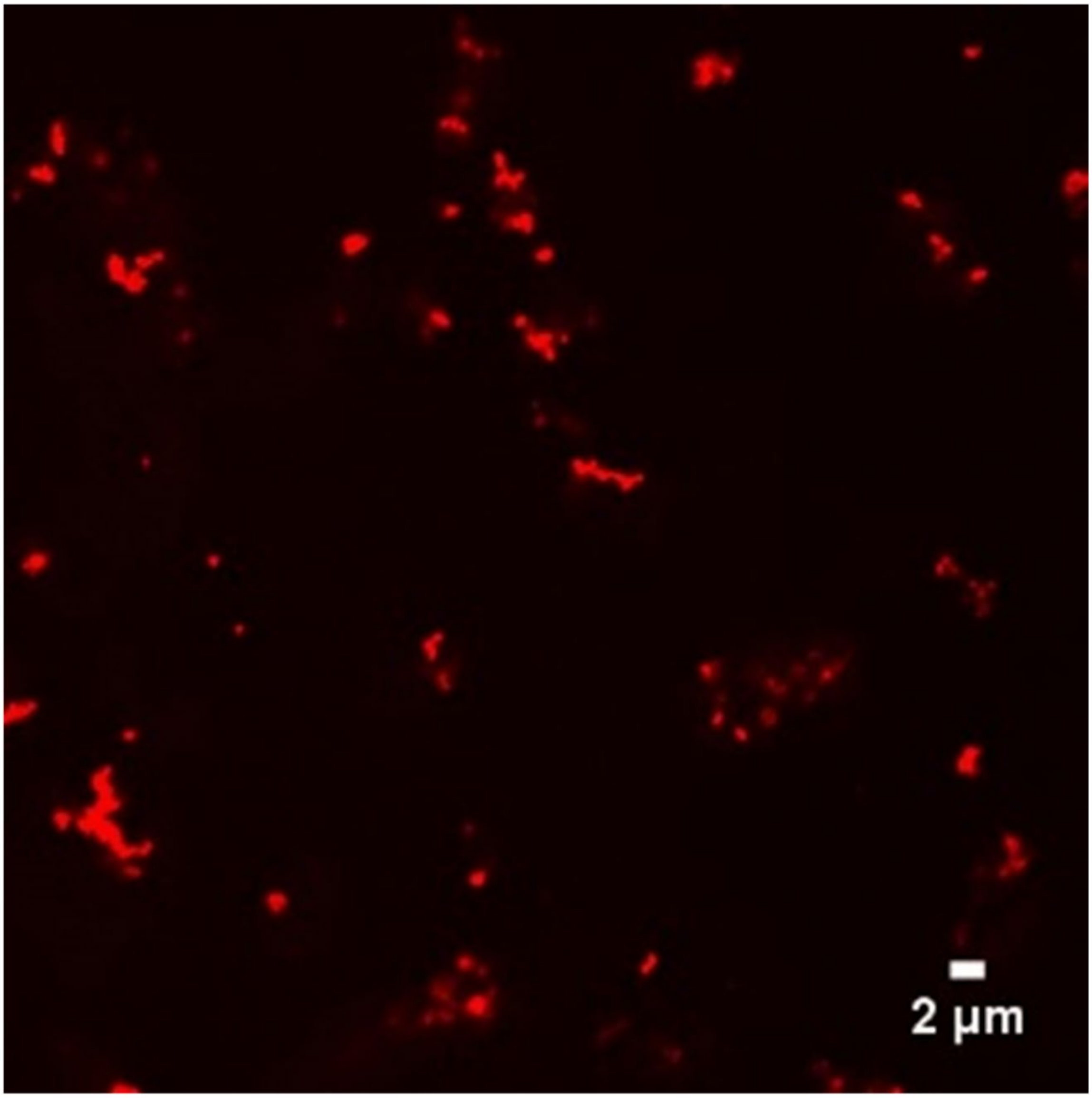


**Fig. S20.** Confocal microscopy of labeled J345 RNA for a mixture between J345 RNA at 0.45 mM and alcohol dehydrogenase at 0.35 mM.

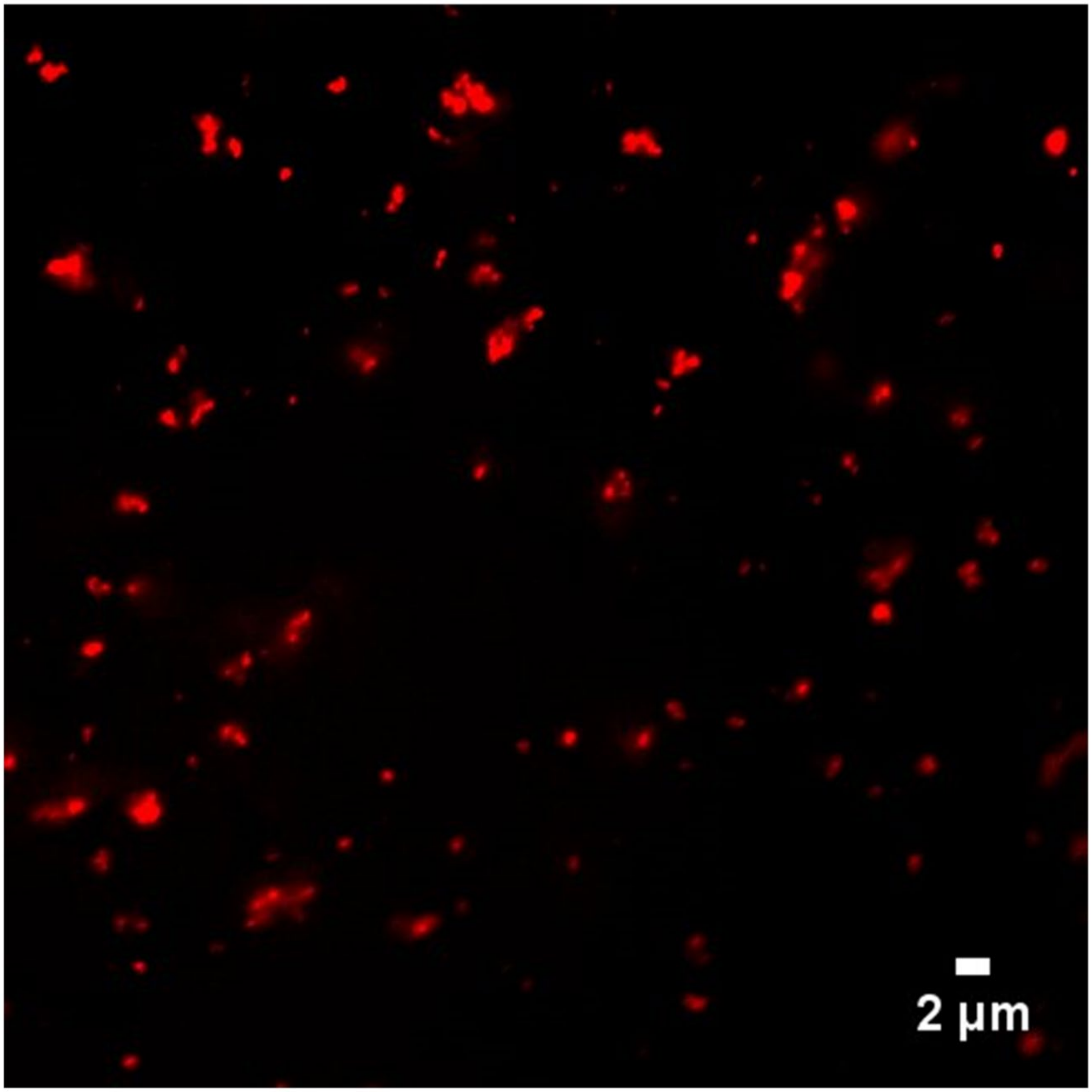


**Fig. S21.** Confocal microscopy of labeled J345 RNA for a mixture between J345 RNA at 0.45 mM and lysozyme at 0.35 mM.

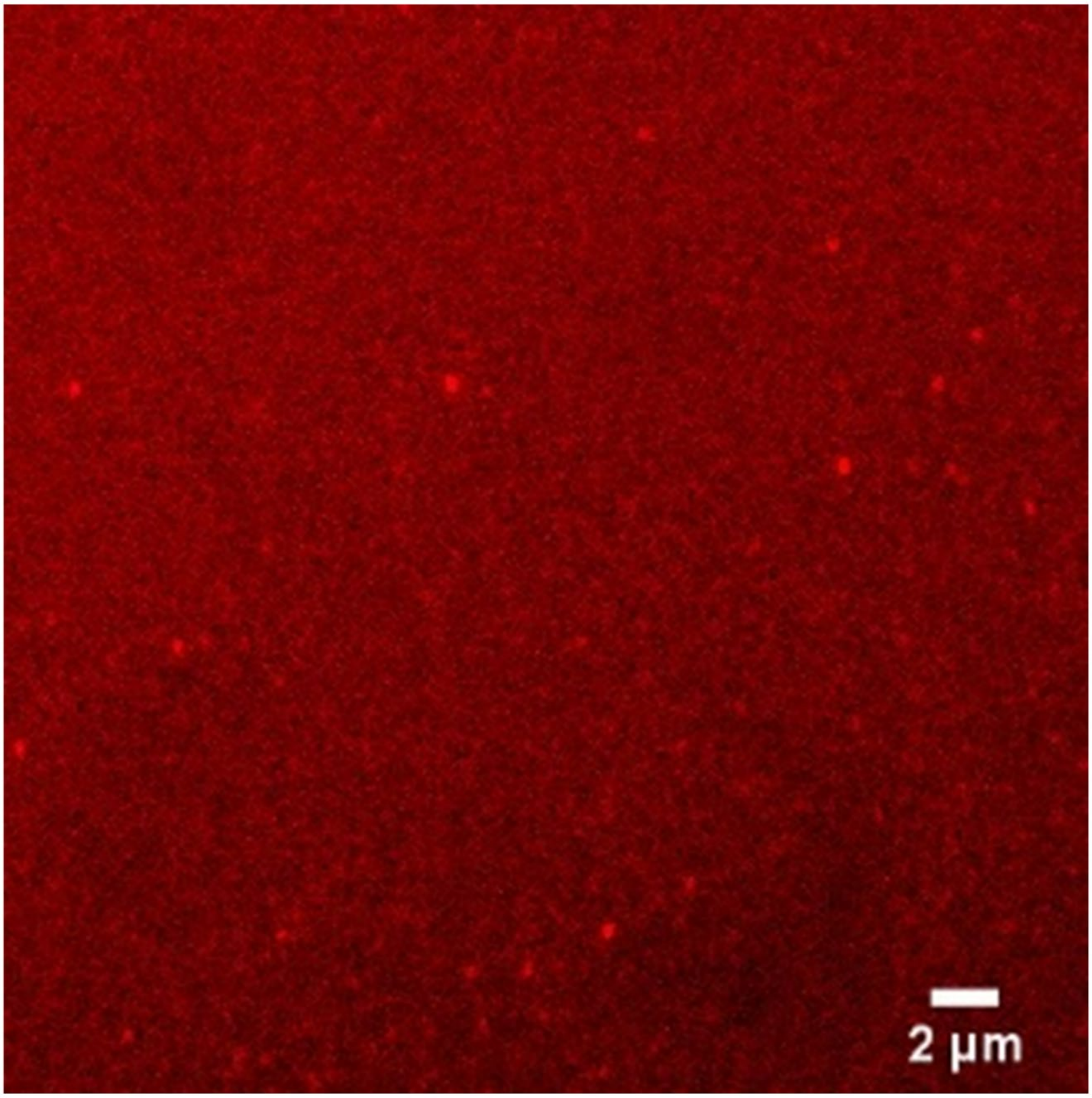


**Fig. S22.** Confocal microscopy of labeled J345 RNA for a mixture between J345 RNA at 0.45 mM and lactate dehydrogenase at 0.35 mM.

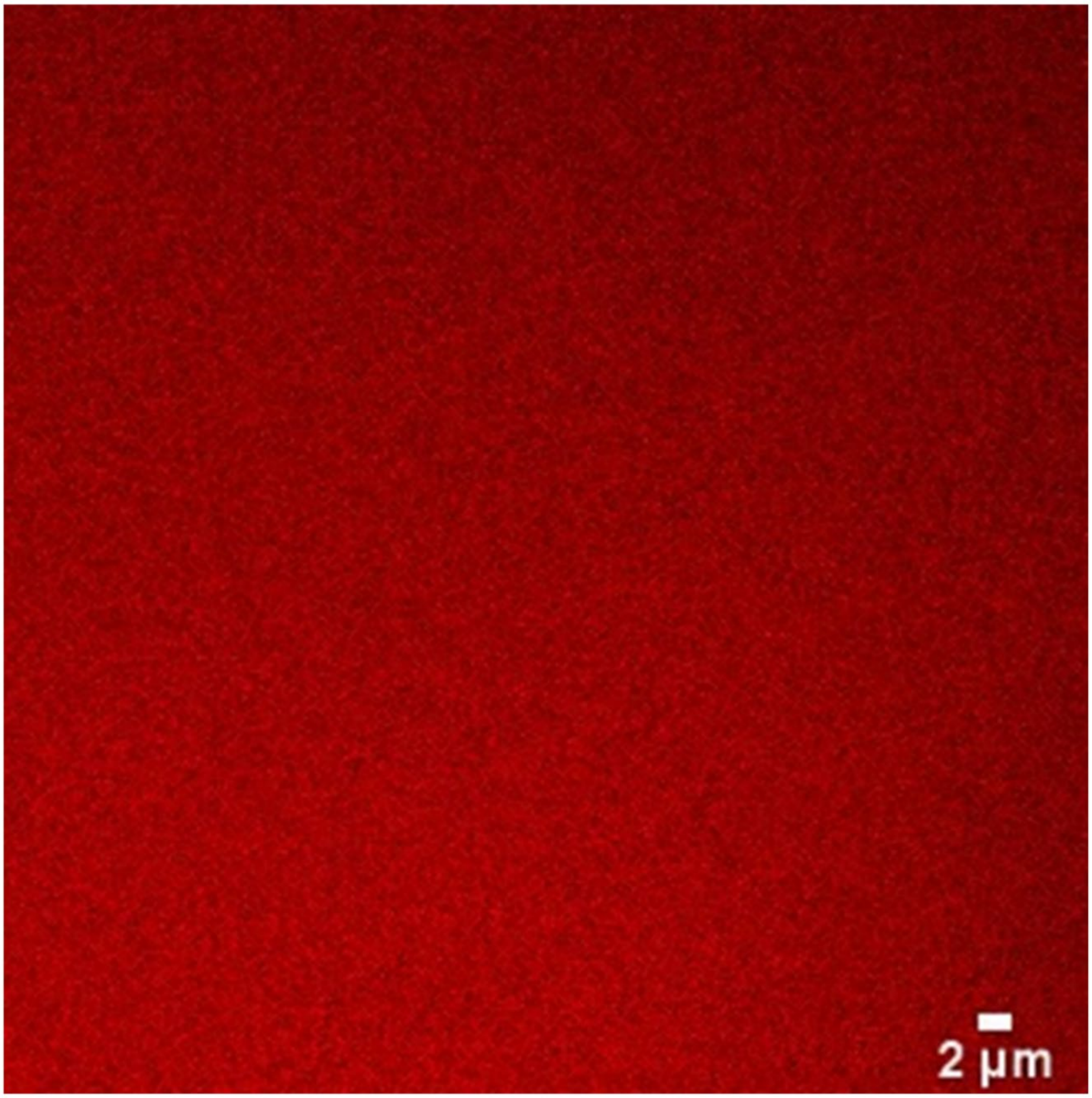


**Fig. S23.** Confocal microscopy of labeled J345 RNA for a mixture between J345 RNA at 0.45 mM and myoglobin at 0.35 mM.

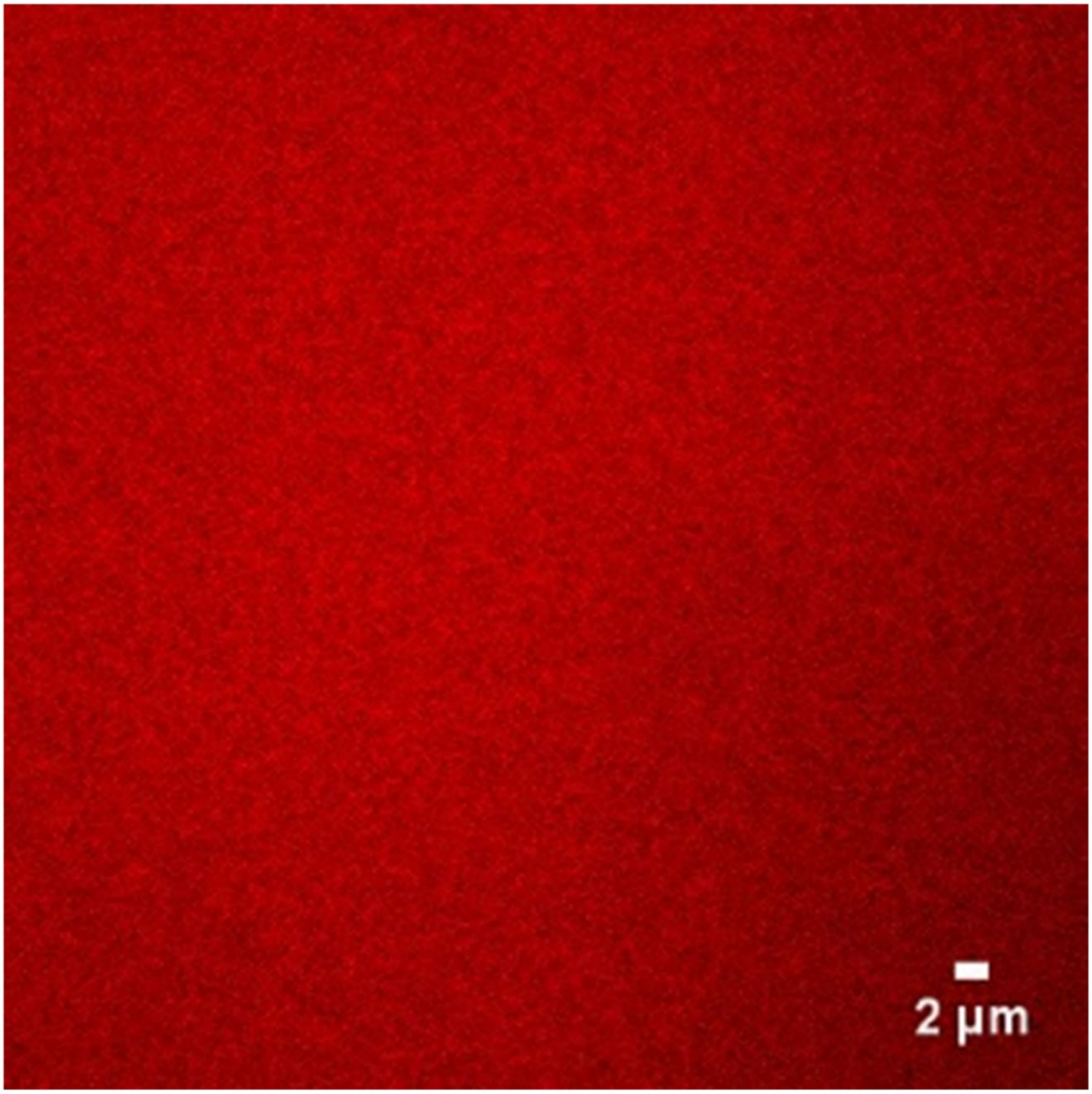


**Fig. S24.** Confocal microscopy of labeled J345 RNA for a mixture between J345 RNA at 0.45 mM and bovine serum albumin at 0.35 mM.

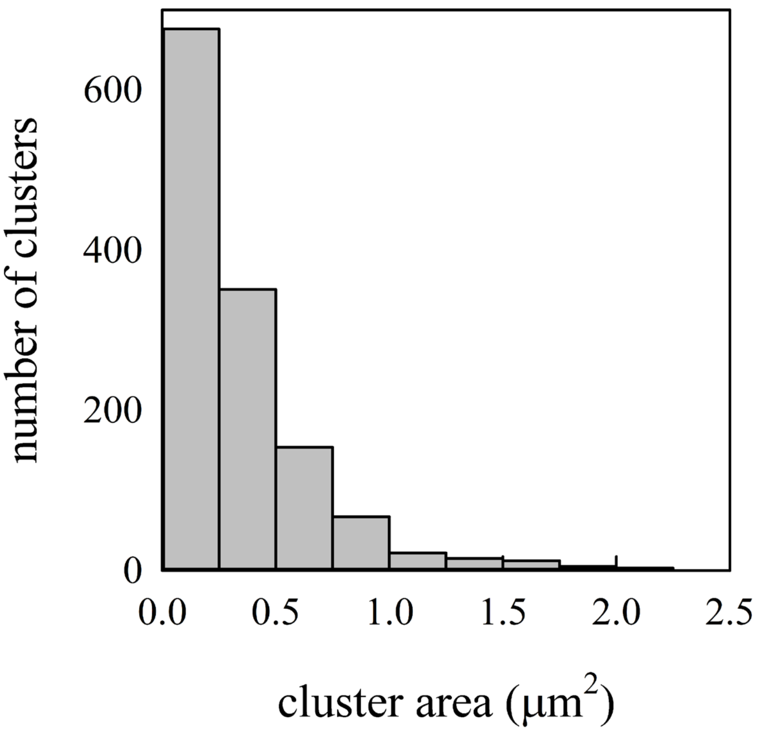


**Fig. S25.** Distribution of cluster sizes from confocal microscopy of labeled J345 RNA in mixtures between J345 RNA at 0.1 mM and trypsin at 0.25 mM. Note that the first bar represents clusters within the diffraction limit of the microscope.

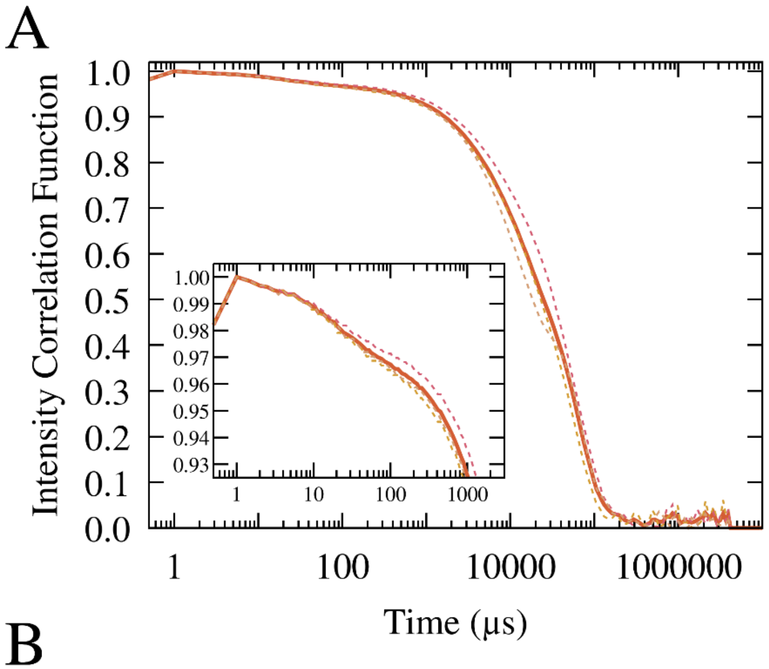


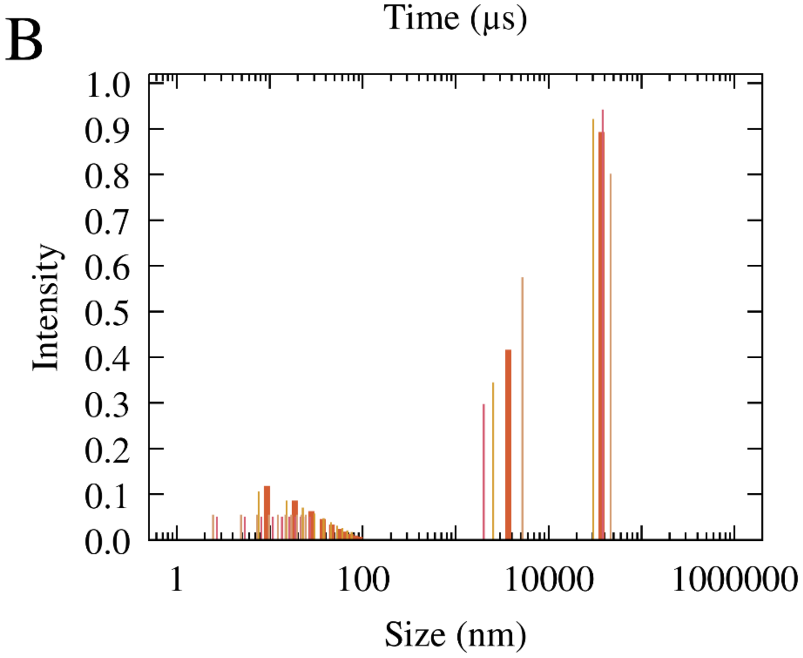


**Fig. S26.** Scattering intensity correlation functions (A) and intensities as a function of particle size from multi-exponential fits (B) from individual dynamic light scattering experiments (dashed/thin lines) of mixtures of 0.1 mM J345 RNA with 0.166 mM trypsin compared with the analysis based on averaged data (thick lines).

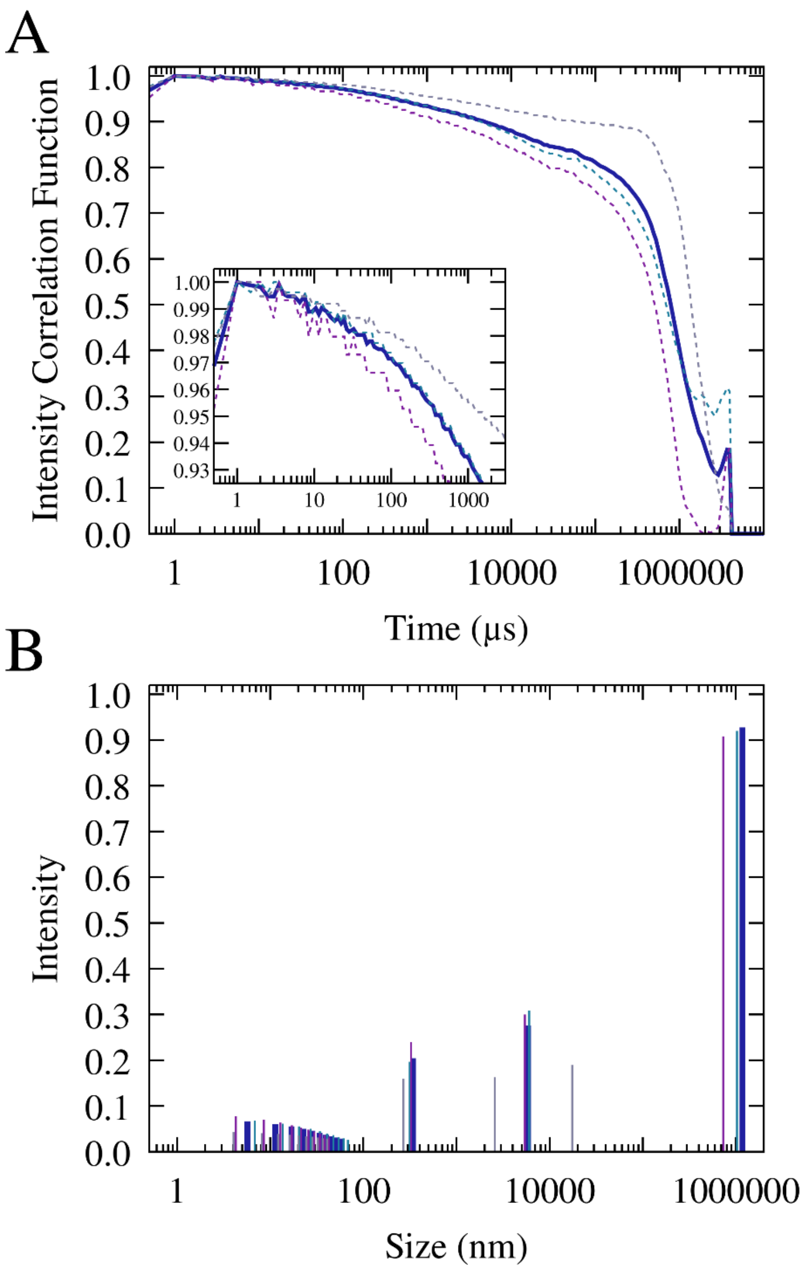


**Fig. S27.** Scattering intensity correlation functions (A) and intensities as a function of particle size from multi-exponential fits (B) from individual dynamic light scattering experiments (dashed/thin lines) of mixtures of 0.4 mM J345 RNA with 0.675 mM lysozyme compared with the analysis based on averaged data (thick lines).

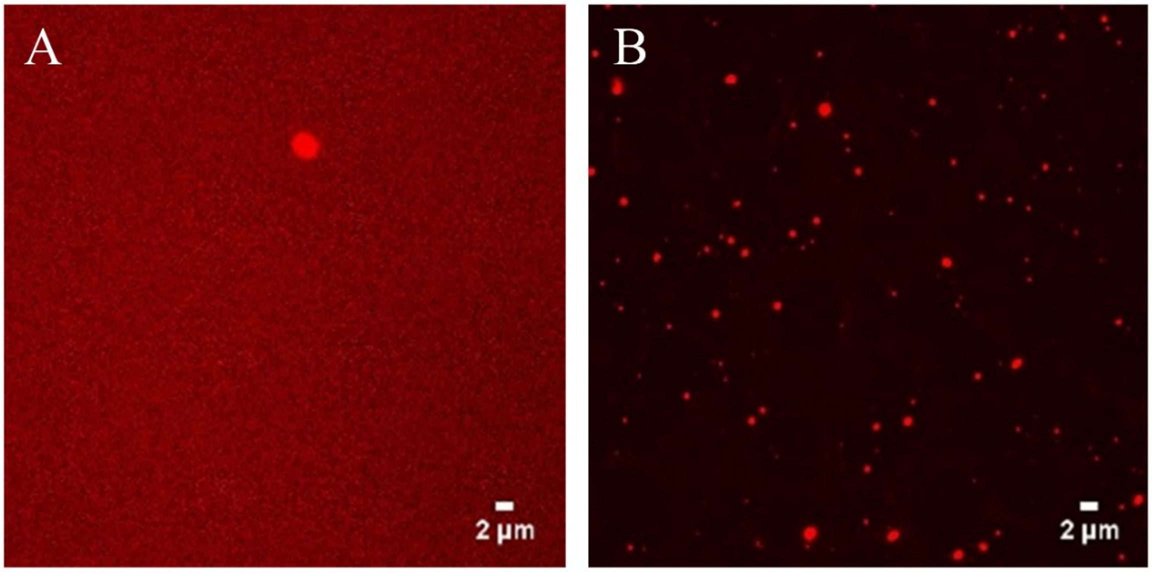


**Fig. S28.** Confocal microscopy of labeled J345 RNA for mixtures between J345 RNA at 0.1 mM and trypsin at 0.05 mM (A) and at 0.15 mM (B). The single bright spot in (A) is attributed to contamination rather than phase separation.

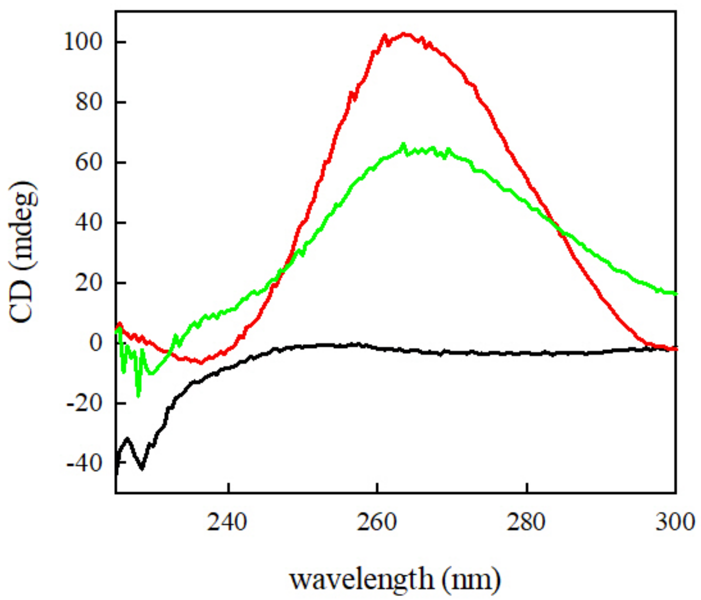


**Fig. S29.** Circular dichroism spectra of trypsin at 0.150 mM (black), J345 RNA, at 0.037 mM (red), and a mixture of trypsin at 0.150 mM and J345 RNA at 0.029 mM (green).

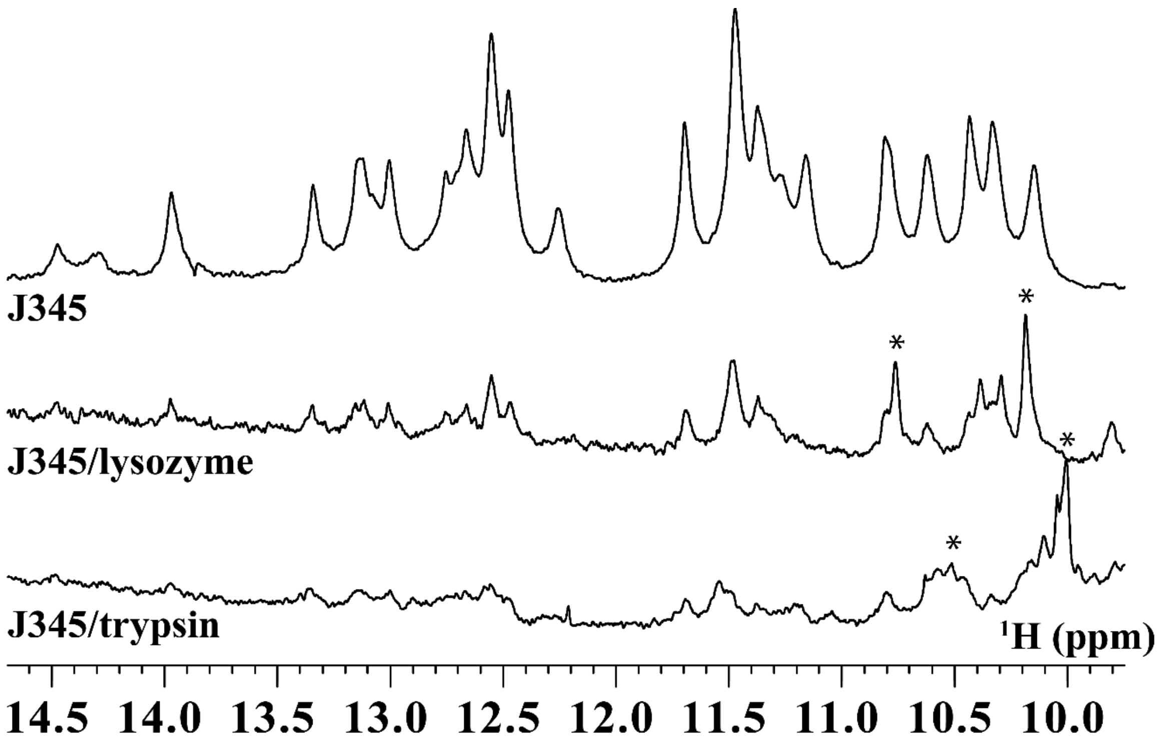


**Fig. S30.** 600 MHz $^{1}H$ NMR spectra in 90:10 $H_2O$:$D_2O$ for J345 RNA only (A) and mixtures of RNA with lysozyme (B) and trypsin (C). Spectral scaling was adjusted to account for higher RNA concentration in the RNA-only sample.

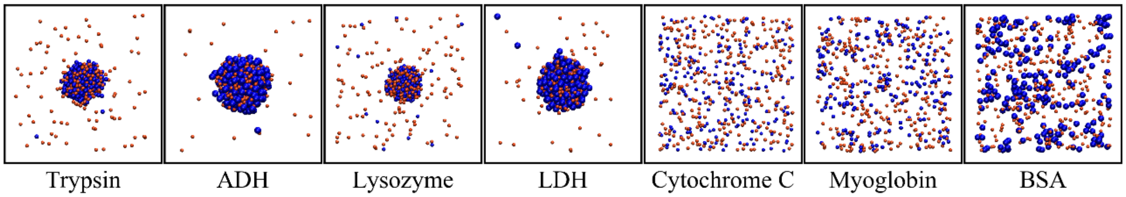


**Fig. S31.** Snapshots from CG simulations after 1 ms for binary RNA-protein mixtures at T = 298K, with $\kappa = 0.75$ using Eq. M-3 to obtain effective charges. [RNA] = 0.493 mM and [protein] = 0.350 mM. Orange and blue spheres show RNA and proteins, according to size. Concentrations inside the condensates were [RNA:lysozyme] = 18.6:18.4 mM; [RNA:trypsin] = 15.6:14.8 mM; [RNA:LDH] = 8.8:7.2 mM; [RNA:ADH] = 8.7:6.6 mM.

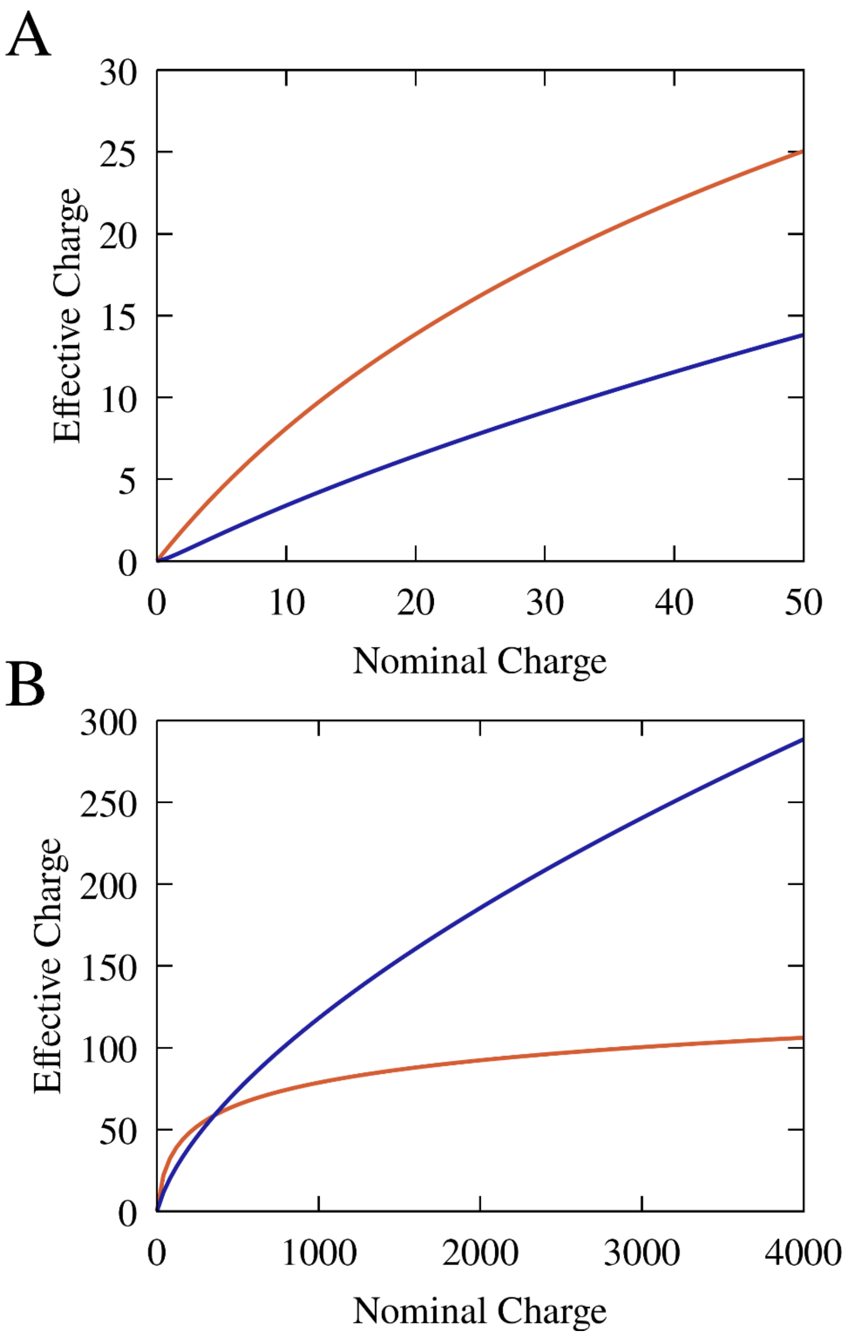


**Fig. S32.** Comparison of effective charge models that take into counterion condensation according to Eq. M-2 (orange) or Eq. M-3 (blue) for moderate (A) and high (B) nominal charges.

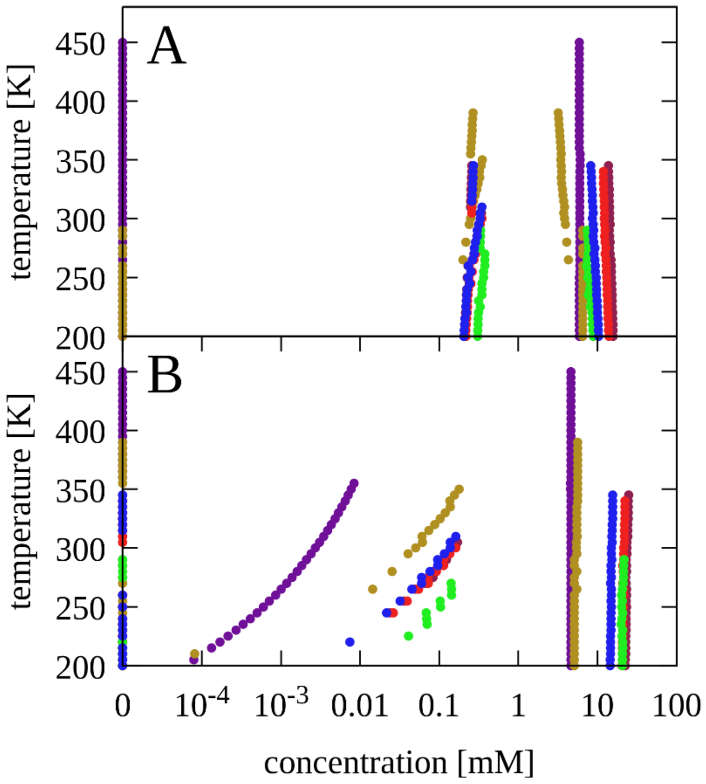


**Fig. S33.** Concentrations of RNA (A) and proteins (B) in dilute and condensed phases as a function of temperature with κ = 1.17. $r_{RNA}$ = 1.47 nm, $q_{RNA}$ = -46, [RNA] = 0.45 mM, [protein] = 0.35 mM. Colors indicate proteins: trypsin (blue), alcohol dehydrogenase (violet), lysozyme (red), lactate dehydrogenase (tan), myoglobin (green), cytochrome C (dark red).

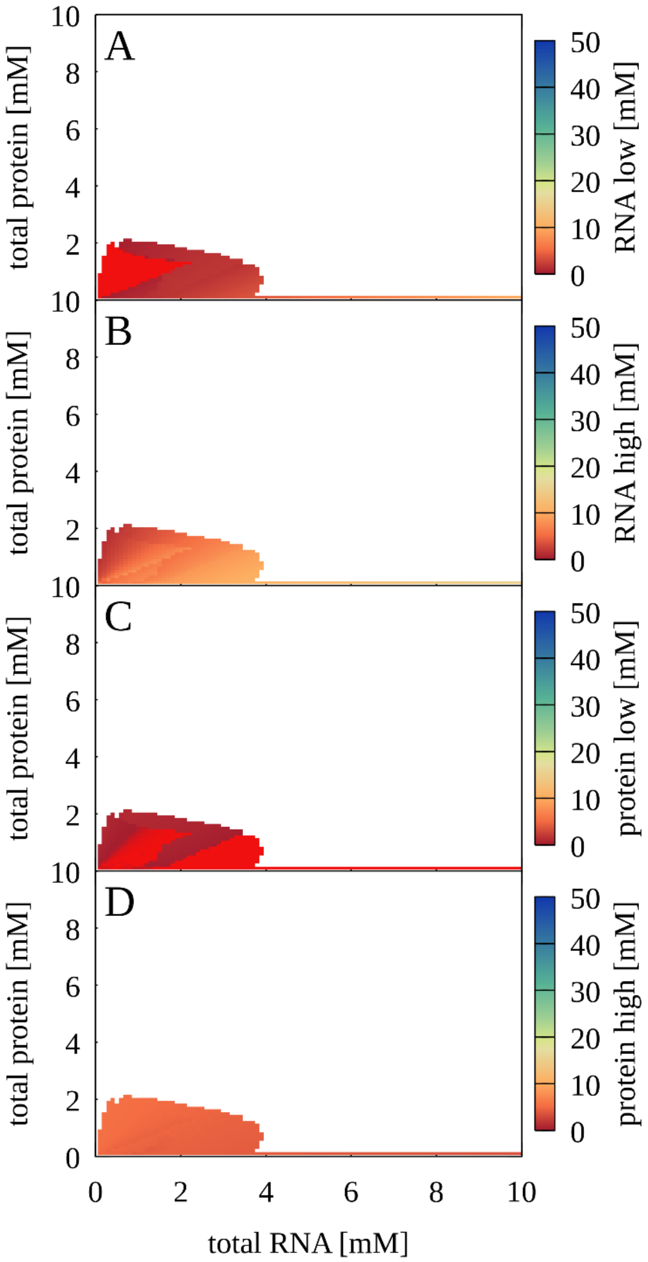


**Fig. S34.** Phase separation for mixtures between RNA and alcohol dehydrogenase as a function of total protein and RNA concentrations. Colors indicate predicted concentrations for RNA (A, B) and proteins (C, D) in dilute (A, C) and condensed (B, D) phases. Bright red color indicates zero concentration (*i.e.* no phase coexistence for that component); no phase separation is predicted for white areas. $r_{RNA}$ = 1.47 nm, $q_{RNA}$ = -46, $r_{protein}$ = 2.79 nm, $q_{protein}$ = 8. The Debye-Hückel screening term was set to κ = 1.17 and T = 298 K.

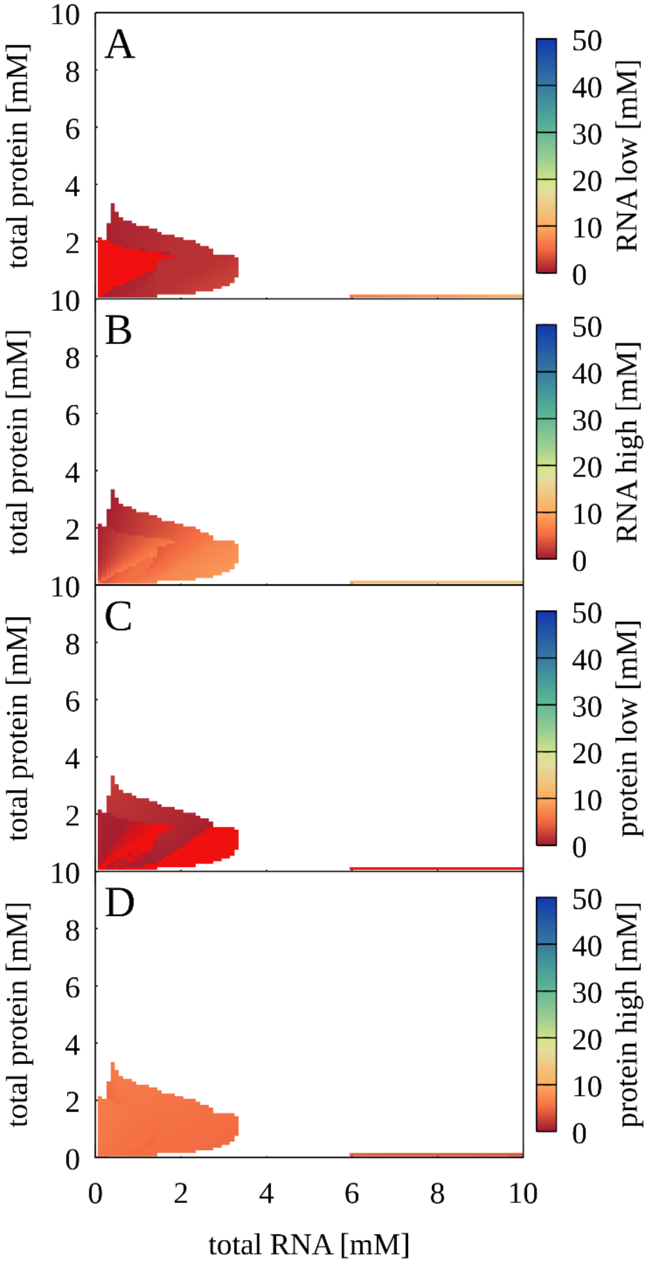


**Fig. S35.** Phase separation for mixtures between RNA and lactate dehydrogenase as a function of total protein and RNA concentrations. Colors indicate predicted concentrations for RNA (A, B) and proteins (C, D) in dilute (A, C) and condensed (B, D) phases. Bright red color indicates zero concentration (*i.e.* no phase coexistence for that component); no phase separation is predicted for white areas. $r_{RNA}$ = 1.47 nm, $q_{RNA}$ = -46, $r_{protein}$ = 2.68 nm, $q_{protein}$ = 4. The Debye-Hückel screening term was set to κ = 1.17 and T = 298 K.

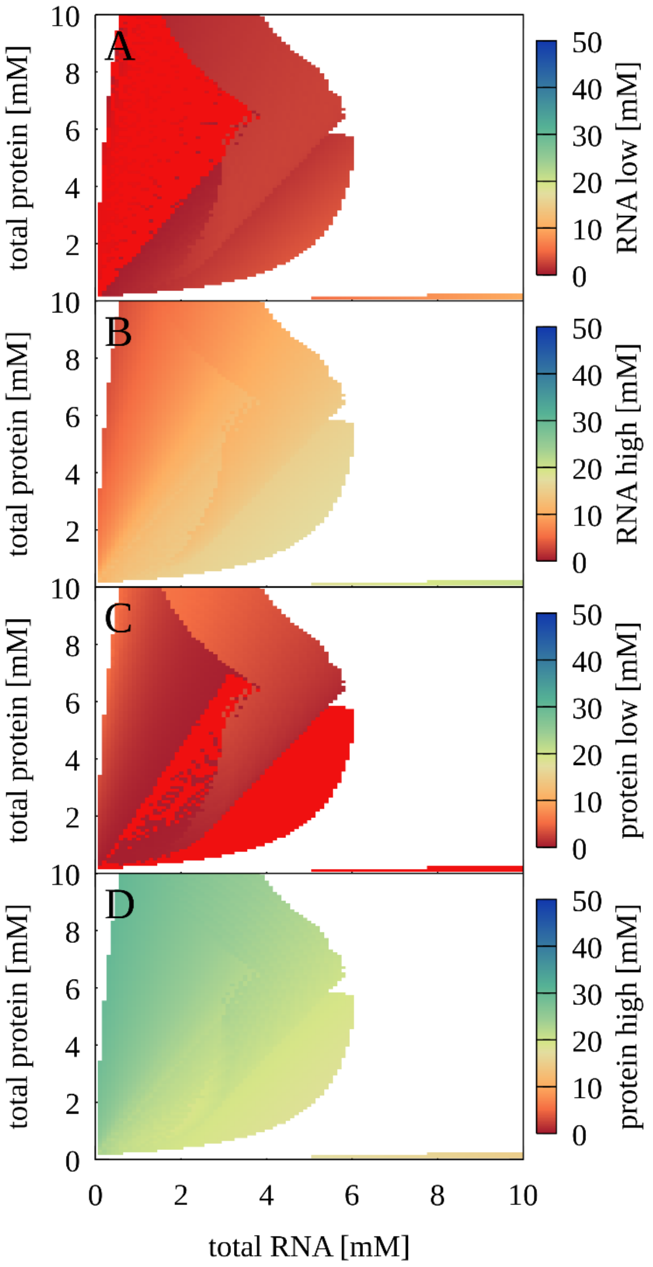


**Fig. S36.** Phase separation for mixtures between RNA and lysozyme as a function of total protein and RNA concentrations. Colors indicate predicted concentrations for RNA (A, B) and proteins (C, D) in dilute (A, C) and condensed (B, D) phases. Bright red color indicates zero concentration (*i.e.* no phase coexistence for that component); no phase separation is predicted for white areas. $r_{RNA}$ = 1.47 nm, $q_{RNA}$ = -46, $r_{protein}$ = 1.54 nm, $q_{protein}$ = 8. The Debye-Hückel screening term was set to κ = 1.17 and T = 298 K.

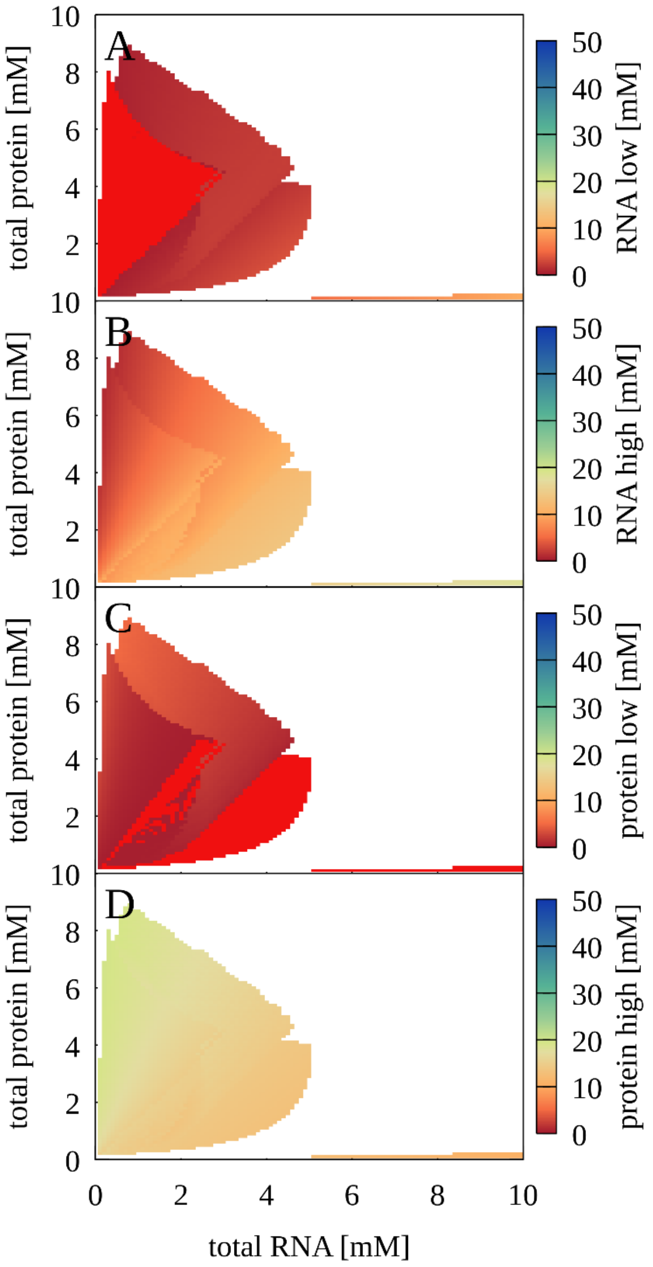


**Fig. S37.** Phase separation for mixtures between RNA and trypsin as a function of total protein and RNA concentrations. Colors indicate predicted concentrations for RNA (A, B) and proteins (C, D) in dilute (A, C) and condensed (B, D) phases. Bright red color indicates zero concentration (*i.e.* no phase coexistence for that component); no phase separation is predicted for white areas. $r_{RNA}$ = 1.47 nm, $q_{RNA}$ = -46, $r_{protein}$ = 1.81 nm, $q_{protein}$ = 6. The Debye-Hückel screening term was set to κ = 1.17 and T = 298 K.

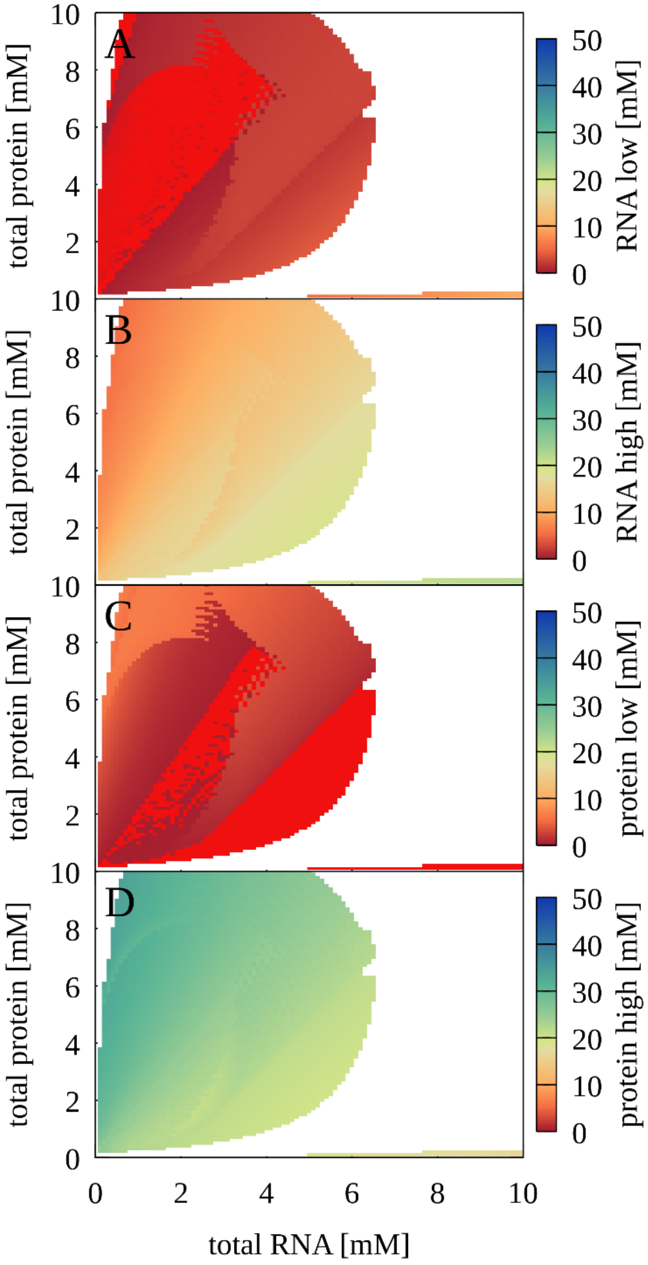


**Fig. S38.** Phase separation for mixtures between RNA and cytochrome C as a function of total protein and RNA concentrations. Colors indicate predicted concentrations for RNA (A, B) and proteins (C, D) in dilute (A, C) and condensed (B, D) phases. Bright red color indicates zero concentration (*i.e.* no phase coexistence for that component); no phase separation is predicted for white areas. $r_{RNA}$ = 1.47 nm, $q_{RNA}$ = -46, $r_{protein}$ = 1.45 nm, $q_{protein}$ = 11. The Debye-Hückel screening term was set to κ = 1.17 and T = 298 K.

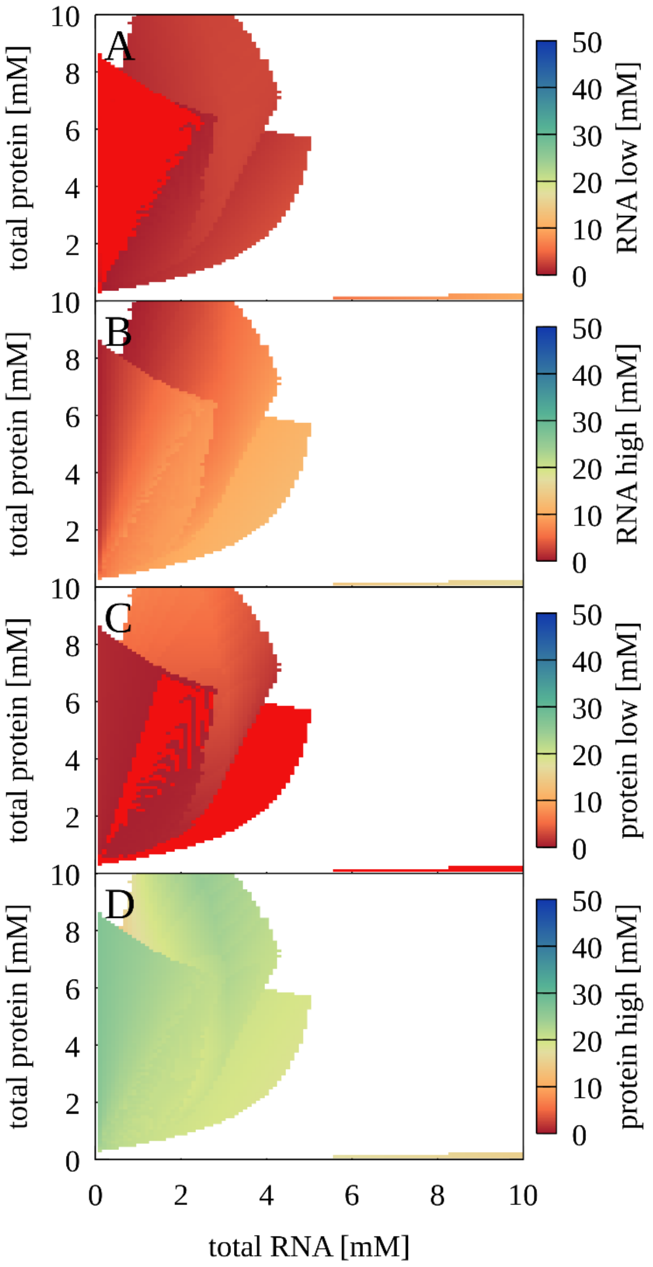


**Fig. S39.** Phase separation for mixtures between RNA and myoglobin as a function of total protein and RNA concentrations. Colors indicate predicted concentrations for RNA (A, B) and proteins (C, D) in dilute (A, C) and condensed (B, D) phases. Bright red color indicates zero concentration (*i.e.* no phase coexistence for that component); no phase separation is predicted for white areas. $r_{RNA}$ = 1.47 nm, $q_{RNA}$ = -46, $r_{protein}$ = 1.64 nm, $q_{protein}$ = 2. The Debye-Hückel screening term was set to $\kappa$ = 1.17 and T = 298 K.

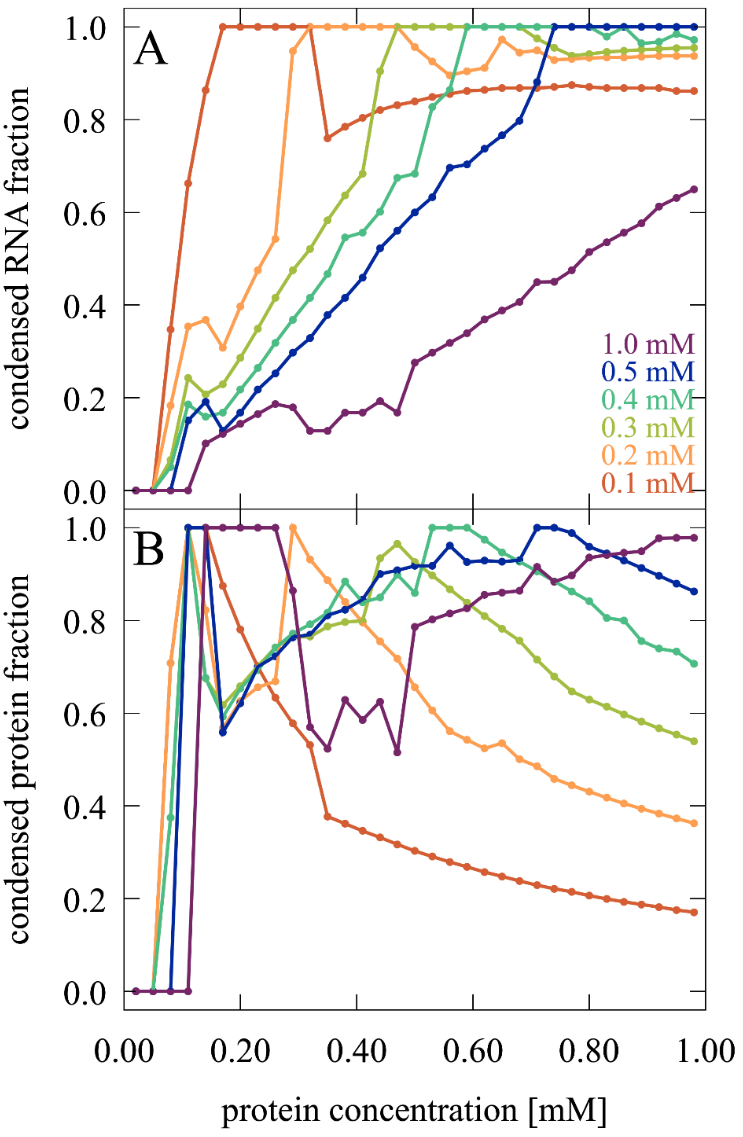


**Fig. S40.** Fraction of RNA (A) and protein (B) in the condensed phases predicted by the theory model for trypsin as a function of protein concentration at different total RNA concentrations. Results represent averages over three subsequent values from values obtained at protein concentrations at increments of 0.01 mM.

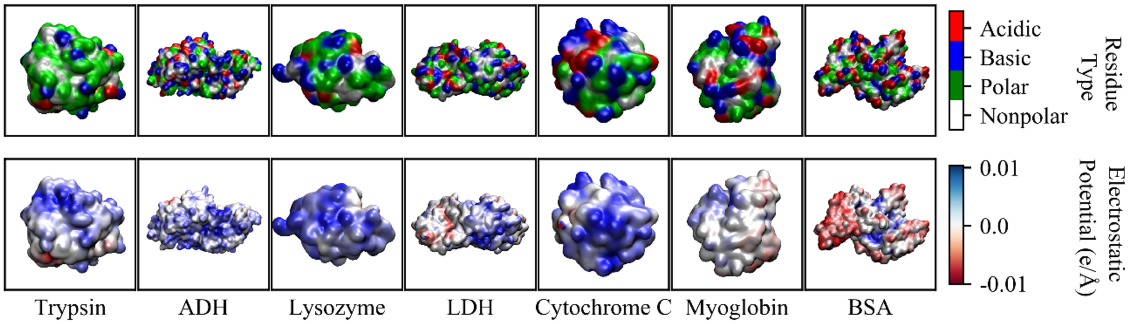


**Fig. S41.** Charge distribution on protein surfaces based on amino acid residue types (top; basic: blue, acidic: red, polar: green, hydrophobic: white) and electrostatic potentials calculated via a Poisson-Boltzmann continuum model (bottom) with coloring according to the sign of the potential (positive: blue, negative: red).

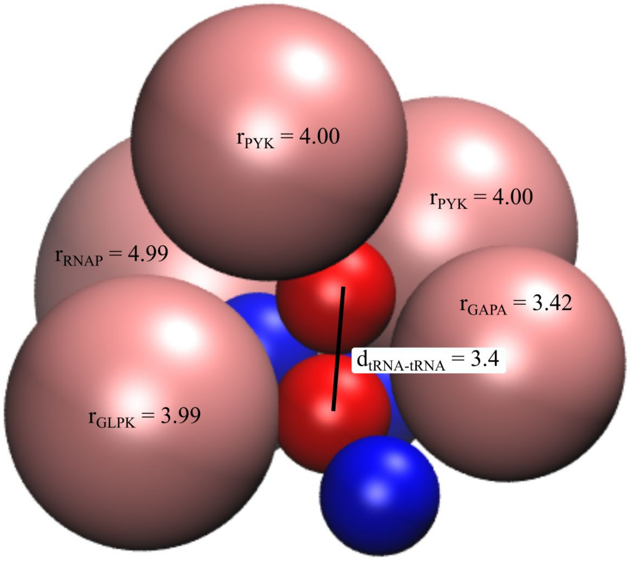


**Fig. S42.** An illustrative example of packing of a tRNA pair (red) in close contact with the positively charged proteins (pink) and other tRNAs (blue) in the cytoplasmic simulations based on the last snapshot after 1 ms simulation. Intermolecular distances (d) and molecular radii (r) are given in nm.

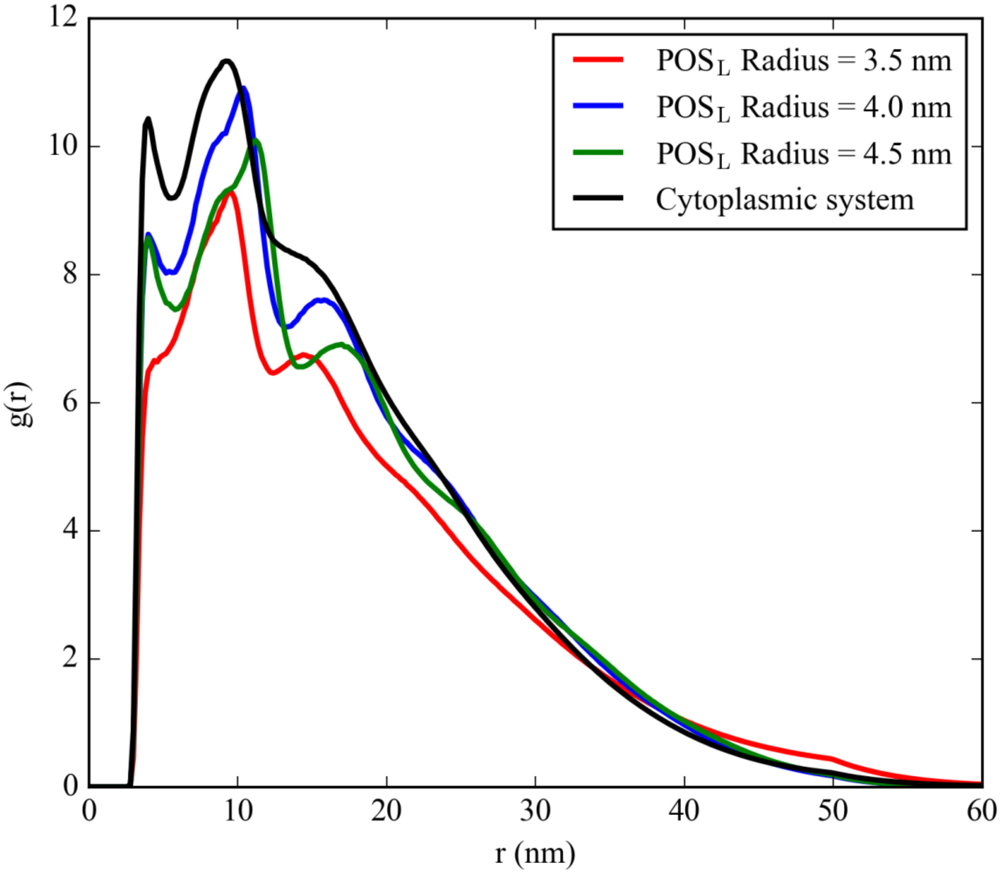


**Fig. S43.** Radial distribution functions for tRNA-tRNA interactions in the five-component model system with different $POS_L$ radii in comparison with the cytoplasmic system.

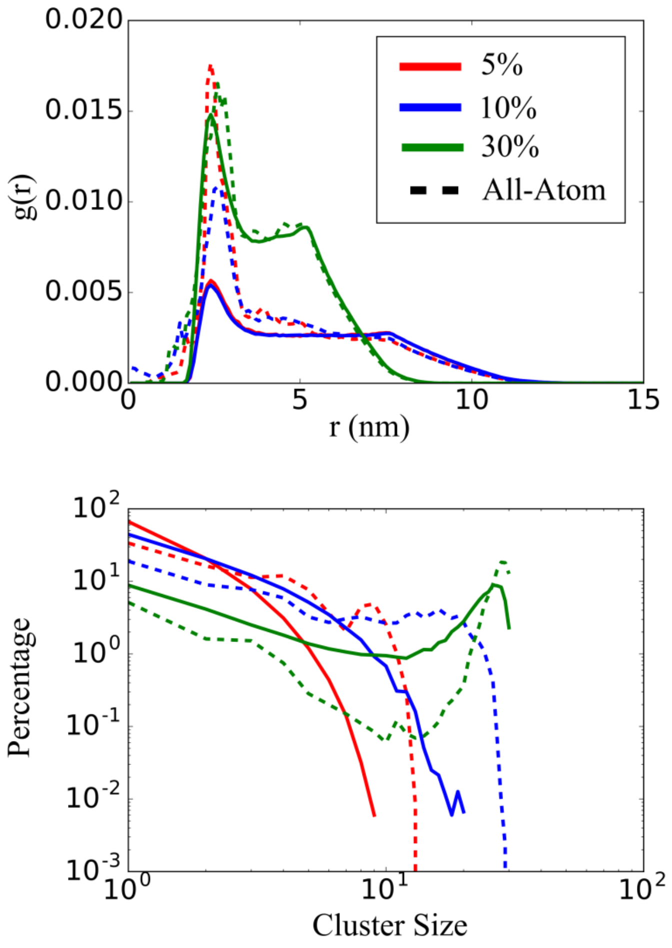


**Fig. S44.** Radial distribution functions of protein-protein pairs (top) and cluster size distributions for proteins (bottom) in simulations of mixtures of villin, protein G, and ubiquitin at volume fractions of 5, 10, and 30%. Dashed lines show results from previously published all-atom simulations (1). Solid lines show results from coarse-grained simulations with the spherical colloid-type model described in the Methods section. A value of $\kappa = 1.5$ was applied and T = 298 K.

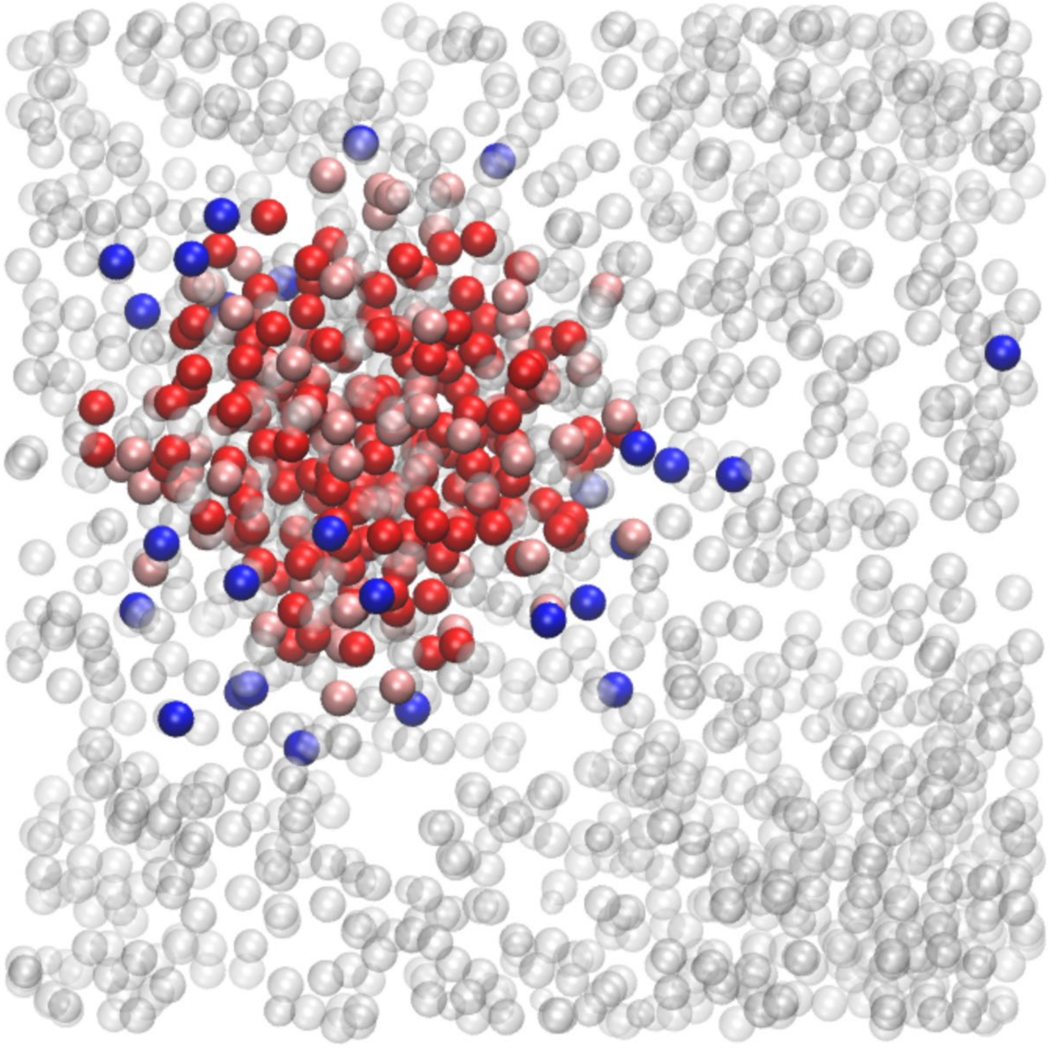

**Fig. S45.** The tRNA cluster at the final snapshot of the cytoplasmic system. tRNAs inside the cluster from pairs determined with a $\sigma_{ij}$+0.7 nm cutoff (see Methods) are shown in red. Additional tRNA molecules included in the cluster with a $\sigma_{ij}$+2.2 nm cutoff are shown in pink. Other tRNA molecules not considered to be part of the cluster are shown in blue, with the rest of the molecules shown in transparent white.

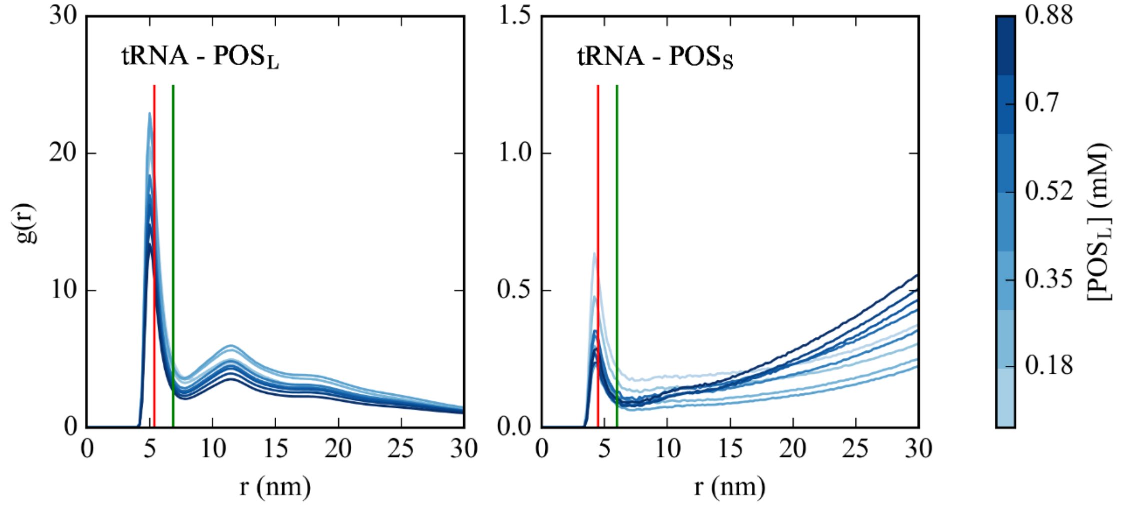


**Fig. S46.** Radial distribution functions between tRNA and $POS_S$ / $POS_L$ particles in simulations of five-component model at different $POS_L$ concentrations and [RP] = 55 μM. Cutoffs based on $\sigma_{ij}$+0.7 nm and $\sigma_{ij}$+2.2 nm are indicated as red and green vertical lines, respectively, with $\sigma_{RNA}$ = 1.55 nm, $\sigma_{POSL}$= 3.12 nm and $\sigma_{POSS}$ = 2.25 nm.

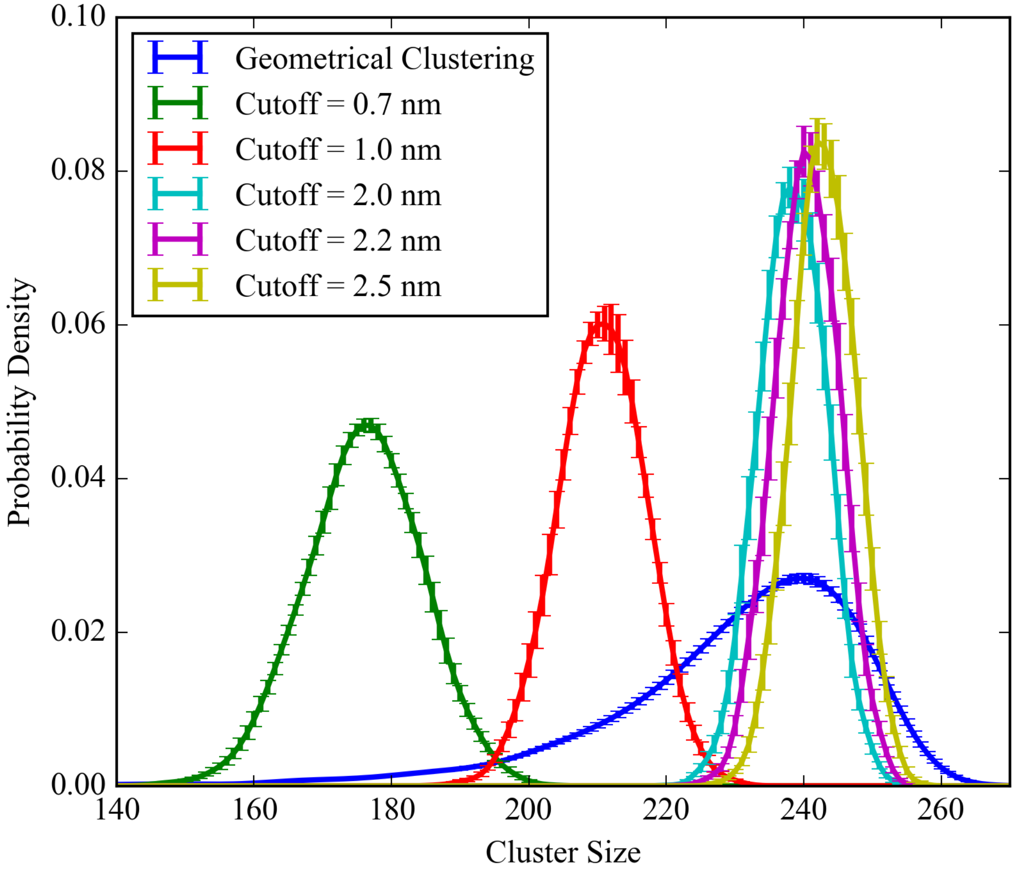


**Fig. S47.** Histograms of tRNA cluster sizes for the cytoplasmic system using the geometrical clustering and based on pairwise contacts using different distance cutoffs added to $\sigma_{ij}$.

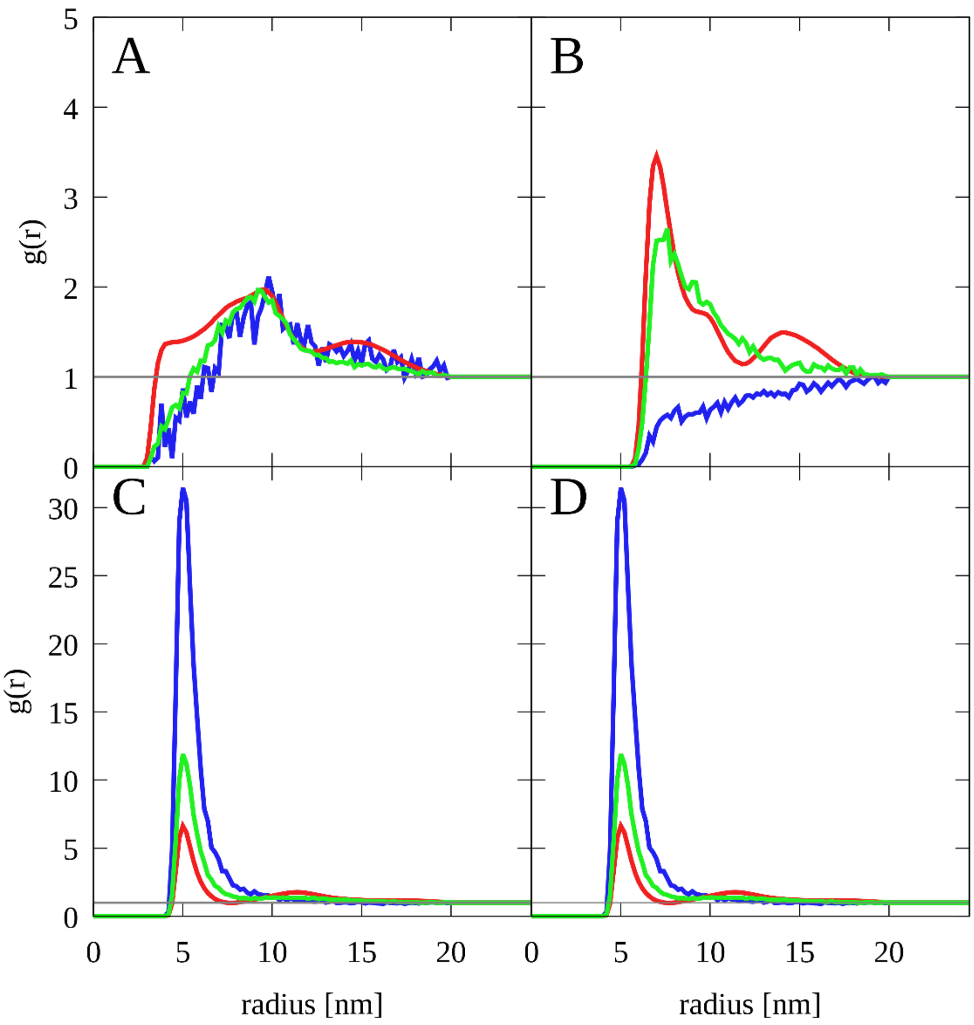


**Fig. S48.** Normalized radial distribution functions for tRNA-tRNA (A), $POS_L$-$POS_L$ (B), tRNA-$POS_L$ (C) and $POS_L$-tRNA (D) interactions in the condensed (red), dilute (blue), and disperse (green) phases used as input for the theory model.

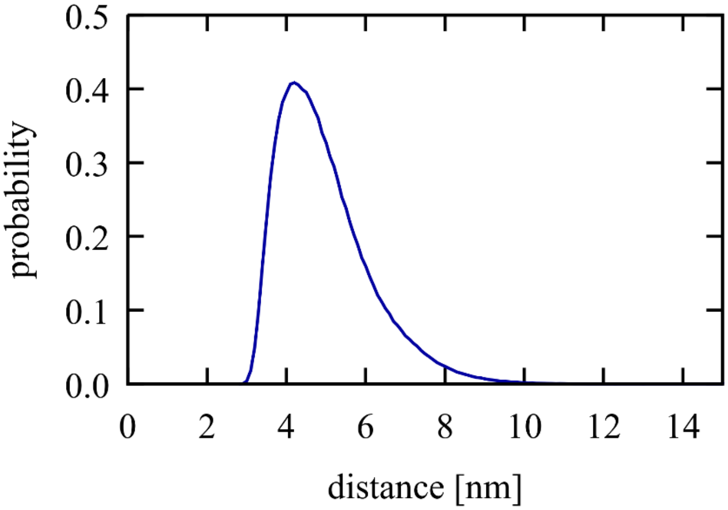


**Fig. S49.** Probability of minimum RNA-RNA distances in the condensed phase from coarse-grained simulations of the five-component model.

**Table S1. Simulation systems for coarse-grained model validation.**

| System | Villin | | | Protein G | | | Ubiquitin | | | Box (nm) |
|---|---|---|---|---|---|---|---|---|---|---|
| Volume Percentage | g/L | mM | $N_p$[1] | g/L | mM | $N_p$[1] | g/L | mM | $N_p$[1] | |
| 5% | 9.7 | 2.3 | 5 | 14.3 | 2.3 | 5 | 19.8 | 2.3 | 5 | 15.3 |
| 10% | 19.0 | 4.5 | 10 | 28.2 | 4.5 | 10 | 39.0 | 4.5 | 10 | 15.4 |
| 30% | 57.9 | 13.8 | 30 | 85.7 | 13.8 | 30 | 118.6 | 13.8 | 30 | 10.6 |

[1]number of proteins.

**Table S2. Multi-exponential fits of dynamic light scattering correlation functions**

| **System**[1] | **Clusters** | | | **Size 1** | | **Size 2** | | **Size 3** | | **Size 4** | | $\chi^2$ |
|---|---|---|---|---|---|---|---|---|---|---|---|---|
| | $D_c$ *(nm)* | $a_c$ | $t_c$ | $D_1$ *(nm)* | $a_1$ | $D_2$ *(nm)* | $a_2$ | $D_3$ *(nm)* | $a_3$ | $D_4$ *(nm)* | $a_4$ | |
| Lysozyme #1 | 6.8 | 0.076 | 9.4 | 314.5 | 0.197 | 6,061 | 0.309 | 1,037,030 | 0.919 | | | 0.000362 |
| Lysozyme #2 | 4.3 | 0.085 | 10.5 | 325.4 | 0.240 | 5,416 | 0.300 | 730,401 | 0.908 | | | 0.00091 |
| Lysozyme #3 | 4.0 | 0.045 | 21.7 | 270.0 | 0.160 | 2,585 | 0.163 | 17,553 | 0.190 | 28,373,600 | 0.948 | 0.00013 |
| **Lysozyme avg.** | **5.6** | **0.073** | **10.4** | **339.4** | **0.204** | **5,848** | **0.275** | **1,184,660** | **0.927** | | | **0.00016** |
| Trypsin #1 | 7.6 | 0.129 | 5.0 | 2,544 | 0.345 | 30,167 | 0.921 | | | | | 0.0016 |
| Trypsin #2 | 2.7 | 0.051 | 186625 | 2,003 | 0.297 | 38,323 | 0.942 | | | | | 0.00146 |
| Trypsin #3 | 2.4 | 0.055 | 106796 | 5,210 | 0.575 | 46,527 | 0.801 | | | | | 0.00105 |
| **Trypsin avg.** | **9.3** | **0.162** | **3.2** | **3,680** | **0.417** | **36,967** | **0.893** | | | | | **0.00231** |

[1]all systems are mixtures between protein and J345 RNA

**Movie S1. Simulation of bacterial cytoplasm model.**

Trajectory of the 100 nm system during the last 1 μs of a 1 ms simulation with tRNAs in orange, ribosomes in magenta, and other molecules colored according to their charges (blue towards positive charges; red towards negative charges). Sphere sizes are shown proportional to molecular sizes. Large pink spheres correspond to GroEL particles.

**Movies S2-S3. Merging of trypsin-RNA liquid condensate droplets**

Video of two representative examples of liquid droplet dynamics in trypsin-RNA mixtures with J345 RNA at 0.45 mM and proteins at 0.35 mM from confocal microscopy of fluorescent-labeled RNA (left) and corresponding bright-field imaging (right). Time evolution is accelerated 25x (i.e. the movies correspond to about 100 s in real time).

**Spreadsheet 1. Components in cytoplasmic model, CG parameters, and self-diffusion.**

Macromolecular components in a previously established model of the bacterial cytoplasm of *Mycoplasma genitalium* (12) with their molecular properties and corresponding CG model parameters. Self-diffusion rates extracted from the CG simulations are reported for different parts of the trajectory and different parts of the system (tRNA clusters, RP clusters, outside clusters).

**Spreadsheet 2. Five-component model simulations.**

List of the five-component model simulations with molecular compositions, CG parameters, and simulation conditions.
